# A hybrid model for multiscale prediction and generation of phase separating protein regions

**DOI:** 10.1101/2025.01.27.635039

**Authors:** Amir M. MohammadHosseini, Hossein Teimouri, Amir M. Cheraghali, Julian Bulssico, Hannah McLaughlin, Daniela Alvarez Robledo, Youssef Abdelmaksoud, Rayan Najjar, Ariel B. Lindner, Vincent Gureghian, Amir Pandi

## Abstract

Liquid-liquid phase separation (LLPS) is emerging as a fundamental process supporting multiple facets of biological systems. This phenomenon enables the dynamic compartmentalization of biomolecules contributing to a wide range of cellular functions, though in many instances its precise role and evolution remain unclear. Protein phase separation naturally occurs within cells and is prevalent across all species. Despite a recent surge in protein LLPS discovery, current predictive models lack generalizability and fail to identify the full spectrum of phase-separating proteins. To address this shortcoming, we developed Phaseek, a hybrid model integrating contextual sequence encoding with statistical graph representations to score LLPS propensity of amino acid sequences. Phaseek accurately identifies phase-separating proteins across diverse biological contexts, predicting key functional regions and the effects of point mutations. Proteome-wide predictions for 18 species highlight important physicochemical features. Gene Ontology enrichments recapitulate known processes (*e.g.*, nucleic acid binding, nuclear localization, chromatin organization) and suggest novel areas of investigation. Phylogenetic analysis of orthologs further suggests that LLPS is evolutionarily conserved beyond sequence similarity. In addition, we used Phaseek to design de novo phase-separating peptides and achieved a 70% success rate *in vivo*. Provided with a user-friendly implementation, Phaseek serves as a multipurpose LLPS predictor for advancing both fundamental and applied LLPS research.

## Introduction

Liquid-liquid phase separation (LLPS) is a physicochemical phenomenon by which molecular entities undergo demixing to form two coexisting phases of different chemical composition^1^. In protein LLPS, specific proteins condensate at nanometer scale and may fuse to form sub-cellular compartments, often referred to as membrane-less organelles that can reach a few micrometers and maintain stable shape and size. Cells typically involve thousands of chemical reactions but a limited number of specialized compartments. Membrane-less organelles provide a means for the cell to achieve spatial organization and functional compartmentalization without the need to synthesize and regulate lipid bilayers^2^.

LLPS was first studied in polymer chemistry in the mid-20th century and only later observed within live cells, when cytoplasmic regions, called P granules, were shown to play a role in the development of *Caenorhabditis elegans* in 2009^3^. The concept of LLPS has since been retrospectively investigated to explain earlier observations of sub-cellular organisations leading to the identification of new specialized sub-compartments, in both the nucleus (nucleolus, nuclear speckles and paraspeckles) and the cytoplasm (the centrosome, stress granules, or germ granules).

Numerous phase-separating proteins involved in various biological contexts and cellular functions have since been identified, highlighting the ubiquity of LLPS in biology^4,5^. As a prime example of a fundamental biological process involving LLPS, RNA polymerases, which catalyze the synthesis of RNA, were shown to form molecular condensates in both prokaryotes and eukaryotes^6,7^. In photosynthetic organisms, the Rubisco protein contributes to the formation of phase-separated sub-structures, the pyrenoids, within the chloroplastic compartment to enhance CO2 fixation^8^. Viruses can form phase-separated viral factories to facilitate the replication of their genome and its packaging into viral particles^9^. Such ubiquitous and important roles of LLPS in biological processes are reshaping our understanding of cellular organization and functions^10^.

Importantly, LLPS plays a crucial role in human health and aging^11^. Dysregulation of phase separation has been associated with multiple pathological conditions, underlying different hallmark processes in cancer but also the formation of protein aggregates in neurodegenerative diseases, of inflammosomes in inflammatory diseases, and of viral factories during infections^12–15^. Thus, condensates constitute promising therapeutic targets^16^ and the prediction of phase-separating proteins can contribute to both basic and clinical research.

A significant fraction of proteins known to undergo LLPS are enriched in intrinsically disordered regions (IDRs), which typically contain fewer hydrophobic amino acids necessary to form rigid tertiary structures and therefore provide the flexibility and conformational plasticity required for multivalent interactions^17^. However, phase separation is not an exclusive property of fully disordered proteins^18,19^. LLPS can also be driven by structured domains or by proteins that contain both folded domains and disordered regions. This sequence and structural heterogeneity, where the driving grammar of LLPS is not confined to a single protein class, poses a significant challenge for sequence-based prediction.

In proteins, LLPS is driven by weak multivalent interactions such as charge-charge, hydrogen bonds, cation-π, π-π, dipole-dipole interactions and hydrophobic contacts^4,5,17–19^. Amino acids, through their side chain, favor different types of interaction: glycine typically engages in hydrogen bonds, while valine favors hydrophobic interactions and more complex amino acids can contribute to multiple interaction types: charge-charge, hydrogen bonds but also hydrophobic interactions in the case of Arginine. Previous studies on IDRs suggested a hierarchy between interactions with charged residues promoting long-range interactions and dipoles contributing to short-range interactions^20^ while aromatic groups may influence the fluidity of LLPS structures^21^.

Machine learning models with ever-growing capabilities have been successfully deployed to address multiple biological problems. Technical progress combined with an increasing number of reported LLPS proteins has enabled the development of LLPS predictive models^22–29^ which often rely on pre-defined physicochemical and sequence features. Such models inherently struggle to generalize, as they fail to capture the complex, context-dependent, and non-local sequence patterns driving LLPS across diverse protein families and species.

To address these limitations, we develop Phaseek, a sequence-based model combining transformers architecture with statistical protein graphs. By integrating deep contextual embeddings (from the transformer) with global statistical properties (captured by the graphs), our approach is designed to restitute the complex, non-local sequence grammar that feature-based models miss, thereby enhancing generalizability. We demonstrate that our model achieves superior performance compared to existing models in predicting diverse phase-separating proteins. We showcase Phaseek ability to detect LLPS-drivers, *i.e.*, key protein regions and to assess the effects of point mutations. We predict phase-separating proteins across 18 organisms and analyze physicochemical and Gene Ontology terms associated as well as their conservation among orthologs. Beyond prediction, we show that the model can be used to generate de novo peptides which when fused to a super-folder Green Fluorescent Protein (sfGFP) achieve phase separation with 70% success in *Escherichia coli* (14 out of 20 new candidates).

Phaseek is provided with a user-friendly implementation through a Google Colab notebook allowing users to predict LLPS score of protein sequences and their key regions or to generate novel LLPS peptide sequences (**Code availability**).

## Results

### Phaseek, a generalizable model for accurate prediction of LLPS proteins

The first of the two core modules in the model is a classic encoder based on Generative Pre-trained Transformer (GPT) architecture^30^ (**Fig. 1a**). For training, we compiled and curated published data (**Methods**), totaling 1,275 positive and 5,258 negative phase-separating sequences. For the positive data, after filtering out sequences exceeding 512aa in length and excluding those with high similarity we obtained a set of 659 positive sequences. We then augmented this set to 4,151 using a trained Hidden Markov Model (**Data availability**) and split the data into training and test sets with a 9:1 ratio. This module alone achieved an AUC (area under curve) score of 0.84 on the test set (**Supplementary Fig. 1**). To enhance the generalization power of the model, we parallelized it with a statistical learning framework.

**Fig. 1:**
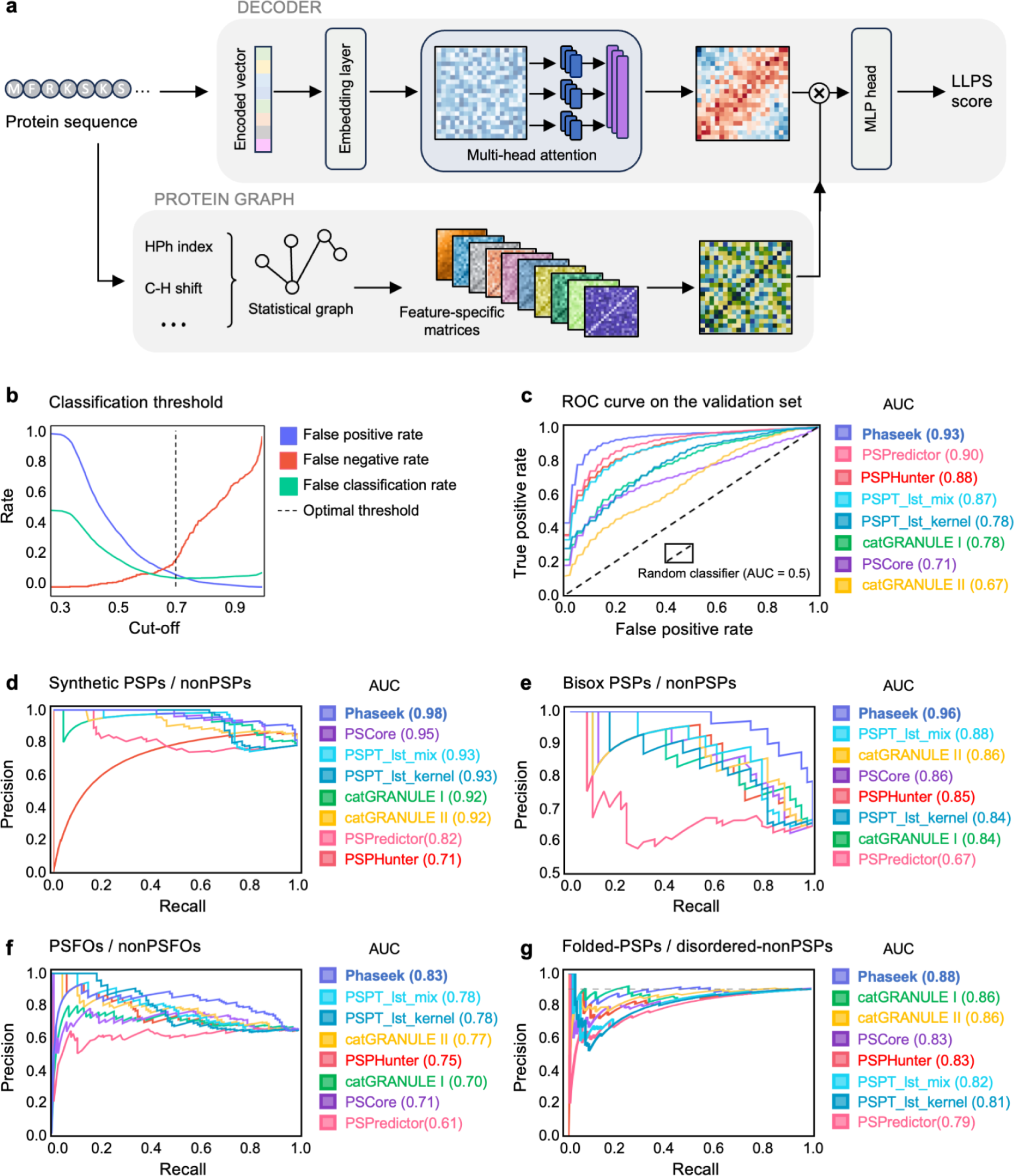
Phaseek architecture and model evaluation. **(a)** The core model comprises two modules: a GPT-based encoder extracting contextual sequence features, and a protein graph generator leveraging feature-specific matrices. The output from both modules is integrated to produce residue-level LLPS scores. For clarity, the XGBoost-based module, used to boost the contribution of LLPS-enriched regions to the final prediction, is not shown. **(b)** False positive, false negative and false classification rate evolution with threshold values used for classification of LLPS proteins based on LLPS score **(c)** ROC (Receiver Operating Characteristic) curves of Phaseek and other models on the evaluation set comprising phase-separating (n = 300) and non-phase-separating (n = 337) sequences. Comparison of the models generalizability with four external datasets: **(d)** Synthetic proteins, including phase-separating proteins (synthetic PSPs, n = 97) and non-phase-separating proteins (synthetic nonPSPs, n = 27). **(e)** Biotinylated isoxazole (B-isox)-identified proteins: phase-separating (Bisox PSPs, n = 43) and non-phase-separating (Bisox nonPSPs, n = 24). **(f)** Fusion oncoproteins: phase-separating (PSFOs, n = 115) and non-phase-separating (nonPSFOs, n = 63). **(g)** Folded phase-separating proteins (Folded-PSPs, n = 175) and disordered non-phase-separating proteins (Disordered-nonPSPs, n = 19). Models are color-coded in the figure: Phaseek (periwinkle blue), PSPhunter (deep salmon), catGRANULE I (Caribbean green), PScore (medium purple), catGRANULE II (saffron yellow), PSPredictor (coral pink), PSTP-LST-mix (bright sky blue), and PSTP-LST-kernel (deep cerulean).

The second core module is a protein graph generator that leverages sequence properties encoded as protein graphs, *e.g.*, average relative probability of inner helix. Feature Extraction based on Graphical and Statistical Features (FEGS) method^31^ generated 158 unique such graph representations corresponding to different protein properties within the training set. To optimize the complexity of using these graph representations as statistical matrices, we applied a random forest model and calculated the SHapley Additive exPlanations (SHAP)^32^ value of each matrix to quantify their predictive power (**Methods**). We selected the top 10 matrices with the highest SHAP values to produce residue-level scores in parallel with our GPT architecture (features listed in **Supplementary Table 1**). Combined, this core model achieved an AUC score of 0.91 on the test set (**Supplementary Fig. 1**).

The third module further enhances the contribution of LLPS sub-regions. Short regions within a protein can carry multiple binding sites, fulfilling the multivalency condition necessary for phase separation, and promote LLPS when fused to other proteins^33,34^. For each sequence, an XGBoost-based module receives amino acid-level scores consolidating them into one, and linearly combines it with the final score of the core model (**Methods, Supplementary Fig. 2**). Transitioning to the three-module version (referred as PhaseekV2 throughout the **Data availability**), the AUC score improved to 0.93 for the test set (**Supplementary Fig. 1**).

We benchmarked Phaseek using an arbitrary threshold of 0.7 balancing false positive and false negative rates (**Fig. 1b**) and compared its performance to six LLPS predictors, including PSPHunter^26^, catGRANULE 1.0^35^, catGRANULE ROBOT 2.0^27^, PScore^36^, PSPredictor^28^, and PSTP^29^ (for information about each model, see **Methods**). Using our test data set, Phaseek resulted in the highest AUC score: 0.93 (PSPHunter: 0.88, catGRANULE 1.0: 0.78, and CatGranule 2.0: 0.67, PScore: 0.71, PSPredictor: 0.90, and PSTP: 0.87 and 0.78) (**Fig. 1c, Supplementary Fig. 3**). We challenged the versatility and generalizability of Phaseek by comparing the models performances in four distinct biological contexts, not seen by any of the models. The first dataset consisted of 97 phase-separating and 27 non-phase-separating reported synthetic proteins^37–41^. Phaseek outperformed all models with an AUC value for the Precision-Recall plot of 0.98 (PSPHunter: 0.71, catGRANULE 1.0: 0.92, and catGRANULE 2.0: 0.92, PScore: 0.95, PSPredictor: 0.82, and PSTP: 0.93 and 0.93) (**Fig. 1d**). The second dataset corresponds to cell extracts from *Arabidopsis thaliana* seedlings and inflorescences^42^ in which intrinsically disordered proteins were precipitated using biotinylated isoxazole (B-isox) and their LLPS behavior was experimentally confirmed (43 phase-separating and 24 non-phase-separating proteins). Phaseek showed the greatest AUC with a score of 0.96 (PSPHunter: 0.85, catGRANULE 1.0: 0.84, and catGRANULE 2.0: 0.85, PScore: 0.86, PSPredictor: 0.67, and PSTP: 0.84 and 0.88) (**Fig. 1e**). The third data set consists of fusion oncoproteins (FOs), which may facilitate oncogenesis through LLPS^33^, totaling 115 phase-separating and 63 non-phase-separating FOs. Phaseek, again, achieved the best AUC with a score of 0.83 (PSPHunter: 0.75, catGRANULE 1.0: 0.70, and catGRANULE 2.0: 0.77, PScore: 0.71, PSPredictor: 0.61, and PSTP: 0.78 and 0.78) (**Fig. 1f**). Since not all intrinsically disordered regions (IDRs) promote LLPS while folded proteins can also display LLPS^18,43^, the fourth dataset contains 19 proteins predicted as disordered but LLPS-negative and 175 folded (non-disordered) but LLPS-positive proteins (**Methods**). Phaseek classified phase-separating and non-phase-separating proteins with an AUC value for the Precision-Recall plot of 0.88, higher than all other models (PSPHunter: 0.83, catGRANULE 1.0: 0.86, catGRANULE 2.0: 0.86, PScore: 0.83, PSPredictor: 0.79, and PSTP: 0.81 and 0.82) (**Fig. 1g**).

Across four diverse biological contexts, Phaseek achieved superior performance in predicting phase-separating and non-phase-separating proteins.

### Phaseek identifies LLPS key regions and predicts the effect of point mutations

Multiple diseases and pathological conditions are associated with mutations occurring in LLPS-promoting regions, so-called key regions, of phase separation proteins^33,34,44–47^. Thus, the identification of such regions is important to both fundamental and translational research.

To assess whether Phaseek identifies key regions, we challenged its ability to predict the effect of truncation for experimentally validated sequences^48–51^. Comparing the score profiles of amino acid sequences at the residue-level, Phaseek (followed by PSTP) performed best in distinguishing LLPS from non-LLPS regions (**Fig. 2a, b, Supplementary Fig. 4a, b**) as measured by a custom match score (**Methods**). For the ORF1 sequence of the human transposable element LINE-1^48^ encoding an RNA-binding protein which has been shown to phase-separate, Phaseek assigned an overall LLPS score of 0.82 (**Fig. 2c**). The model also detected previously identified LLPS key regions^48^ and captured their overall effect as removing residues 47-53 (Region A) or 132-152 (Region B) from the protein sequence reduced the LLPS score to 0.43 and 0.37, respectively (**Fig. 2c**, additional examples for RNA pol II and Tau protein **Supplementary Fig. 5**).

**Fig. 2:**
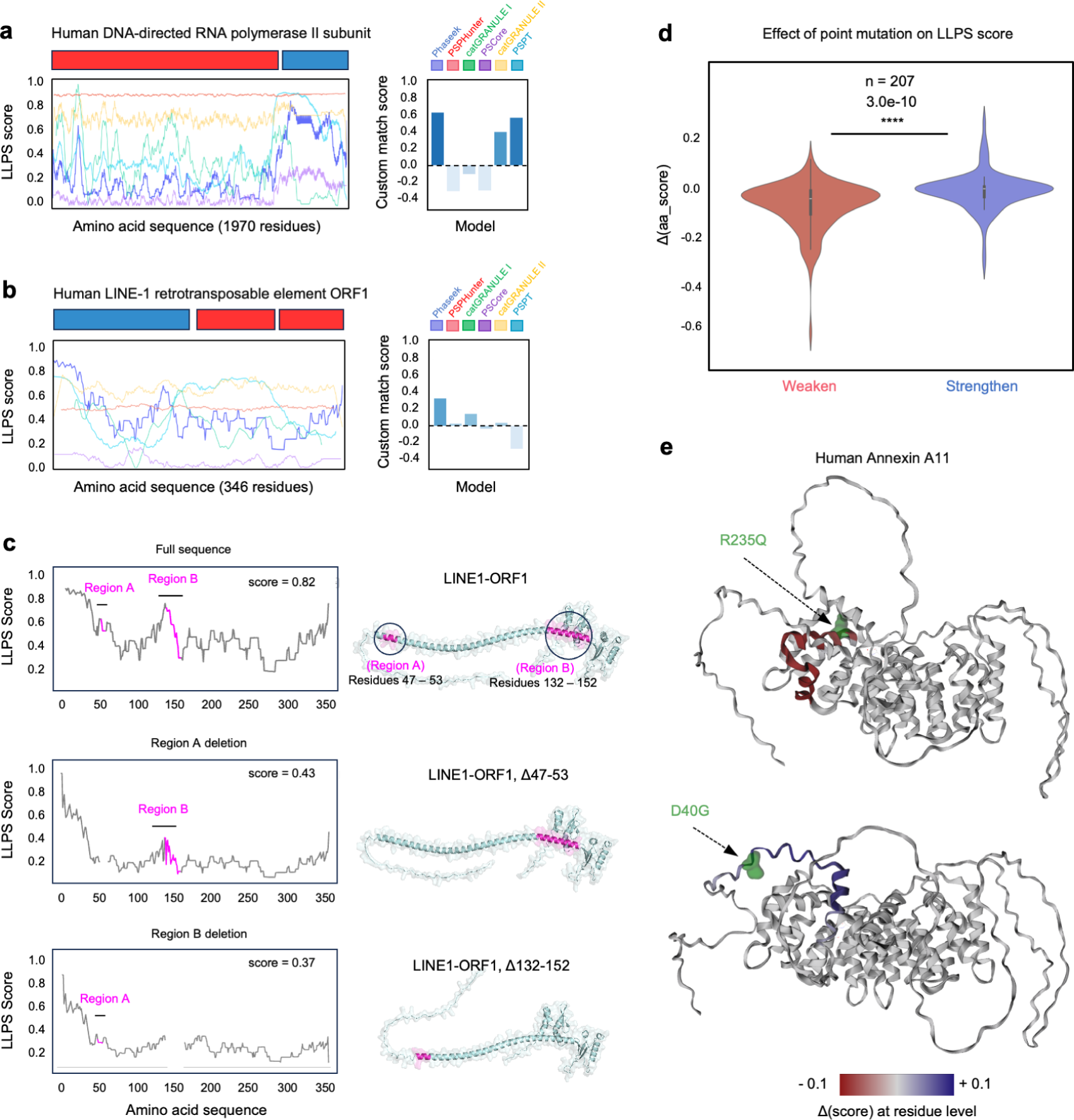
Phaseek identifies LLPS key regions and predicts mutation effects. **(a)** Comparison of LLPS predictions, from different models generating residue-level LLPS profiles, for human RNA polymerase II subunit with phase-separating (blue) and non-phase-separating (red) regions. **(b)** Residue-level LLPS predictions from various models for LINE1-ORF1 containing two LLPS segments: residues 47–53 (Region A) and 132–152 (Region B) **(c)** Comparisons of Phaseek results for the wild-type form (LLPS score: 0.82), deletion of Region A (LLPS score: 0.43), and deletion of Region B (LLPS score: 0.37). Structures were predicted using AlphaFold2. **(d)** Effects on LLPS scores of 90 strengthening mutations and 111 weakening mutations (violin plots on the right) **(e)** Predicted effects of two point mutations, LLPS-weakening R235Q and LLPS-strengthening D40G, on Human Annexin A11 (UniProtKB: P50995). Changes in residue-level LLPS scores are shown on AlphaFold2-predicted structures, with decreased scores in red, increased scores in blue, and mutated residues in green. Predictions by catGRANULE I and PScore in panels (a, b) were scaled to a 0–1 range. Model colors: Phaseek (periwinkle blue), PSPhunter (deep salmon), catGRANULE I (Caribbean green), PScore (medium purple), catGRANULE II (saffron yellow), PSTP-LST-kernel (deep cerulean). The statistical significance was assessed with one-sided Mann–Whitney U tests (****p < 0.0001).

To investigate whether the model would capture the translational fusion of an LLPS region, we compared sequence predicted scores between native non-phase-separating proteins and their fusion to phase-separating peptides ^52 37^. For interferon-alpha A (IFN-alpha A) fused to an artificially designed elastin-like protein (ELP)^52^, all models except catGRANULE 2.0 correctly predicted LLPS scores below a threshold of 0.5 for IFN-alpha A and above for ELP and IFN-alpha A-ELP (**Supplementary Fig. 4c**). For the LLPS peptide HERD2,2, fused to mCherry and a flavin-containing monooxygenase (FMO)^37^, only Phaseek correctly predicted LLPS scores below 0.5 for mCherry (0.49) and FMO (0.23), and above for HERD2,2 (0.89) and HERD2,2-mCherry-FMO (0.80) (**Supplementary Fig. 4d**).

To challenge the model ability to capture point mutations, we collected data for 2,529 LLPS-determining mutations for 137 phase-separating proteins from multiple studies^53–55^ (**Data availability**) and simulated 20 random mutations for each of the 137 proteins, yielding a total of 2,740 random mutations (**Methods**). We calculated three custom metrics: Δ(score), Δ(aa_score), and Σ(Δ(aa_score)) representing the changes in score at the sequence, the amino acid, and the regional level (**Supplementary Fig. 6 and Methods**). For missense mutations, the global Δ(score) and the local Σ(Δ(aa_score)) resulted in significant differences (p-value = 1.3×10^-24^ and 7.6×10^-13^) while the Δ(aa_score), which focuses solely on the mutated residue, showed no significant difference (p-value = 0.70) (**Supplementary Fig. 7**). This difference between global, local and residue score suggests that the model considers contextual information beyond single amino acid. These observations hold true when considering separately mutations happening in folded or IDR regions, noteworthy the statistical significance was greater for folded regions (**Supplementary Fig. 8**).

To evaluate if the model is able to predict the effects of point mutations, we used a set of 207 sequences with experimentally characterized 90 LLPS-promoting and 111 -weakening mutations. Changes in LLPS scores were significantly different between the two categories for all scores (residue level Δ(aa_score) **Fig. 2d**, global Δ(score) and local Σ(Δ(aa_score)) **Supplementary Fig. 9**). Some LLPS-weakening mutations had more impact on the score than promoting mutations. As an example, we examined mutations in human Annexin A11^56^, which facilitates RNA transport in neurons and has been implicated in amyotrophic lateral sclerosis^56^. Phaseek captured the effects of both weakening (R235Q, H395P, and R456H) and strengthening (D40G, R346C, and G38R) mutations on LLPS (**Fig. 2e and Supplementary Fig. 10**).

Together, these results show that Phaseek, which has been trained on whole sequences, identifies key functional sub-sequences driving LLPS and predicts the effect of mutations at residue level.

### Phaseek captures LLPS determinants and the prevalence of IDRs in key regions

LLPS is driven by amino acid sequences with specific physicochemical and structural properties^57^. To investigate the determinants of LLPS, we analyzed the proteomes of 18 organisms: *Escherichia coli*, *Saccharomyces cerevisiae*, *Plasmodium falciparum, Arabidopsis thaliana*, *Anopheles gambiae*, *Drosophila melanogaster*, *Caenorhabditis elegans*, *Danio rerio*, *Xenopus laevis*, *Gallus gallus*, *Rattus norvegicus*, *Mus musculus*, *Canis lupus familiaris*, *Bos taurus*, *Sus scrofa*, *Macaca mulatta*, *Pan troglodytes*, and *Homo sapiens*. We used Phaseek to score coding sequences, both manually curated sequences from Swissprot and computationally predicted sequences from trEMBL (**Supplementary Fig. 11, Data availability**)

We analyzed the correlations between the predicted LLPS score and 12 physicochemical features across proteomes (**Fig. 3a**). Among these features, the IDR score which reflects the average disorder propensity as predicted by AIUPred^58^, showed the strongest positive correlation (Spearman ρ = 0.45). This aligns with disordered regions facilitating effective multivalent inter-/intra-molecular interactions^59^. Conversely, hydropathy, the fraction of hydrophobic residues, and the fraction of aromatic residues exhibited inverse correlations (ρ = −0.41, −0.40, and −0.35, respectively). Polyproline II (PPII) helix propensity and glycine/proline fraction were positively correlated (ρ = 0.33 and 0.27). Meanwhile, Aromatic clustering, net charge, and Kappa charged amino acid patterning (which quantifies the tendency of positive/negative amino acids to form clusters) did not correlate. These correlations were coherent across species besides *E. coli* having weaker correlations and *H. sapiens* less correlated features (**Supplementary Fig. 12**).

**Fig. 3:**
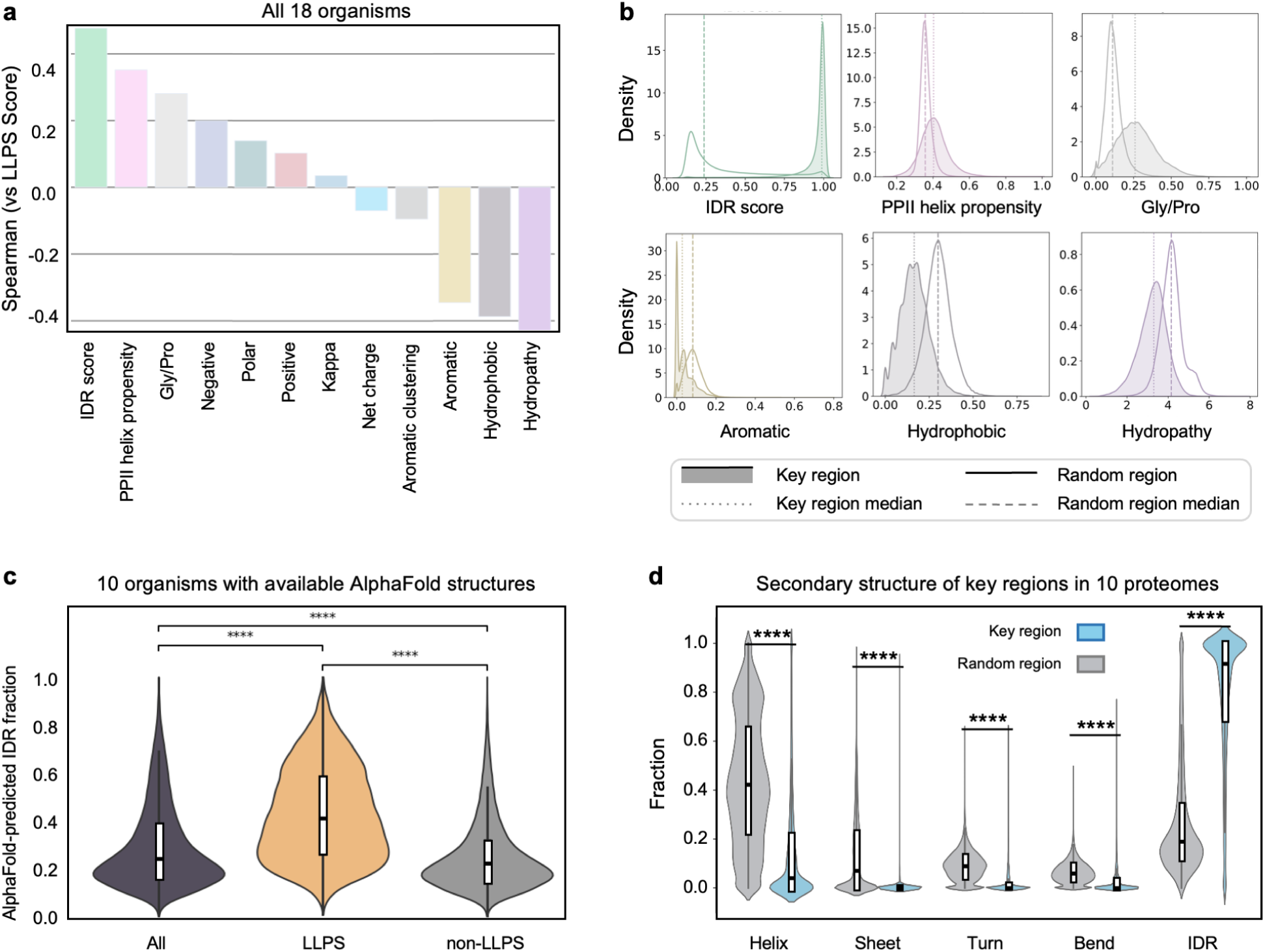
Intrinsically disordered regions discriminate between key and random regions. **(a)** Spearman correlation coefficients between physicochemical features and Phaseek LLPS scores of the same 10 proteomes. **(b)** Density plots of physicochemical feature distributions in random versus key regions for the same 10 proteomes, with medians shown. One-sided Mann–Whitney U tests yielded p < 0.0001 for all comparisons shown. **(c)** Violin plots of AlphaFold-predicted IDR fraction for all proteins of the 10 proteomes, LLPS positive and LLPS negative proteins. **(d)** Violin plots compare secondary structure distributions for random and key regions across the 10 proteomes with AlphaFold data (*E. coli*, *P. falciparum*, *S. cerevisiae*, *A. thaliana*, *D. melanogaster*, *C. elegans*, *D. rerio*, *R. norvegicus*, *M. musculus*, and *H. sapiens*). A one-sided Mann–Whitney U test produced p < 0.0001 for all five structural classes. Boxes indicate the interquartile range (Q1–Q3) and medians. Feature labels in (a, b) use shorthand: aromatic (fraction of F, W, Y), hydrophobic (A, V, I, L, M), Gly/Pro (G and P), polar (S, T, N, Q, C, H), positive (K, R), and negative (D, E).

We compared the distributions of these features between key regions identified by Phaseek and random regions (**Methods**). To quantify differences, we analyzed the common-language effect size (CLES), which is the probability of having a greater score when drawing at random one observation from each distributions^60^. Among the most correlated features, the IDR score exhibited the greatest separation between key and random regions. Key regions showed a CLES = 0.956, indicating that random regions had greater values in 95.6% of pairwise comparisons and lower values in only 4.4% (**Fig. 3b** and **Supplementary Fig. 13** for species-specific plots). In contrast, other correlated features like PPII helix propensity and Gly/Pro as well as the negatively correlated features, Aromatic, Hydrophobic and Hydropathy, exhibited less pronounced differences. Their CLES values were 0.576, 0.807, 0.215, 0.115 and 0.139 respectively. Notably, these distributions remained consistent across species (Supplementary Fig. 14-31)

Using AlphaFold^61^ structure predictions for 10 organisms, we explored the relationships between the secondary structures and LLPS scores assigned by Phaseek (**Supplementary Fig. 32** and **Supplementary table 3**). Indeed, LLPS proteins showed a greater proportion of disordered regions, with a greater median IDR fraction for LLPS proteins than for non-LLPS proteins (**Fig. 3c** and **Supplementary Fig. 33** for species-specific plots). We further compared secondary structures (helices, sheets, turns, bends, and IDRs) between random and key regions. Key regions were largely depleted of ordered structures as compared to random regions, with median fraction of 0.035 vs 0.419 for helices, respectively (**Fig. 3d**). Conversely, key regions were significantly enriched in disordered structures, median of 0.914 vs 0.189 in random regions. Notably, key regions contained very few sheets, turns or bends, with *E. coli* showing the highest proportion among species (species-specific plots **Supplementary Fig. 34**)

Overall, these results show that the model captures different physicochemical properties supporting phase separation and highlights the prevalence of IDRs in LLPS key regions.

### Phaseek identifies Gene Ontology terms associated with LLPS

To shed light on a potential association of LLPS to biological functions, we enriched Gene Ontology (GO) terms in the LLPS positive fraction of 17 organisms with an available OrgDb R package (excluding *P. falciparum* **Supplementary table 3)**. We further tightened the results by applying semantic similarity to filter a representative subset of parent terms, mimicking the REVIGO project methodology^62^ (**Methods**).

Enriched terms reported across multiple organisms are coherent for the three GO categories and align with previous observations that LLPS involves protein complexes which can be located in the nucleus and interact with nucleic acids to regulate chromatin organization. Across these three ontologies, most enriched terms were species-specific: 2,480 terms appeared in only one species before applying semantic similarity, and 628 remained after. In contrast, only 16 and 1 terms respectively, were shared across all 17 species (**Supplementary Fig. 35**). To pinpoint fundamental aspects of LLPS, we focused on GO terms most broadly shared across organisms, with *E. coli* being the only prokaryotic organism in our panel and having fewer shared terms (**Fig. 4**). Gene set sizes were similar across species, supporting meaningful cross-species comparisons. We further provide a detailed species-specific analysis for further exploration beyond the premise of this work (**Supplementary Fig. 36-52**).

**Fig 4:**
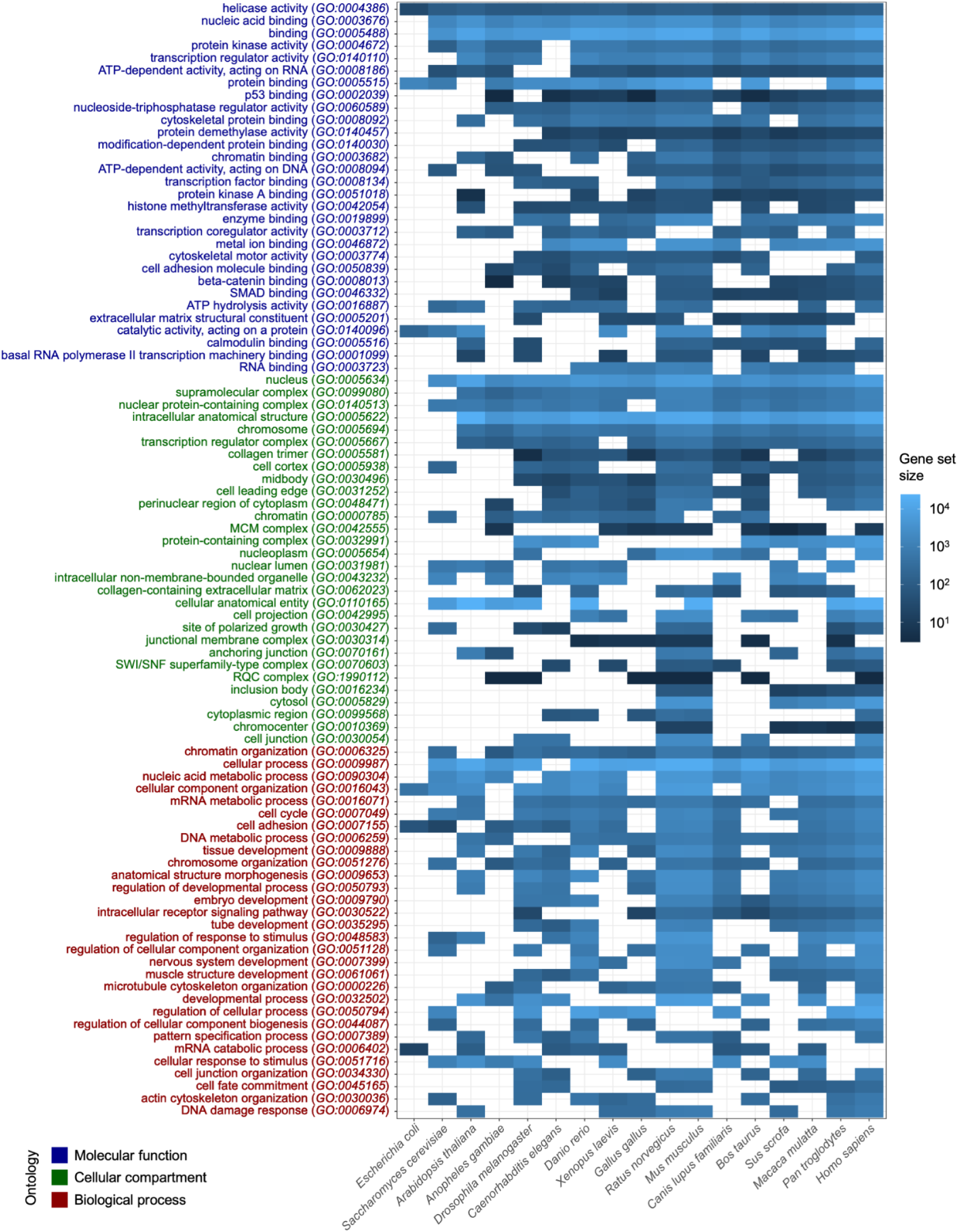
Gene Ontology (GO) terms enriched in the LLPS positive fraction of 17 organisms proteome. Enriched terms for each organism were summarized using semantic similarity as implemented in the rrvgo R package. Summarized terms are ranked by occurrences (top to bottom) across organisms ranked by evolutionary complexity (left to right). Terms are colored according to the ontology to which they belong, blue for Molecular function, green for Cellular compartment and red for Biological Process.

Within the **Molecular Function** category (*blue*, **Fig. 4),** the top result reported for all organisms is helicase activity (GO:0004386, n=17), in accordance with their known involvement in membrane-less organelles such as P bodies and stress granules ^63^. Terms enriched across multiple organisms are related to binding of multiple biological components such as nucleic acid (GO:0003676, n=16), protein (GO:0005515, n=13), chromatin (GO:0003682, n=12), transcription factor (GO:0008134, n=11) and enzymes (GO:0019899, n=11); similarly captured by the more generic term binding (GO:0005488, n=16). Terms, as nucleoside tri-phosphate regulator activity (GO:0060589, n=13) and modification-dependent protein binding (GO:0140030, n=12), are in line with the role of ATP and phosphorylation in LLPS regulation^7,64–66^.

In the **Cellular Compartment** category (*green,* **Fig. 4)**, the top result compartment, nucleus (GO:0005634, n=16), is enriched in all organisms but *E. coli* and is coherent with the existence of multiple non-membrane organelles in the nucleus ^67^. Four terms are related to protein complexes such as supramolecular complex (GO:0099080, n=15), nuclear protein-containing complexes (GO:0140513, n=15) and transcription regulator complexes (GO:0005667, n=14) in agreement with the role of LLPS in enhancers and transcription regulation ^6,7, 68^.

In the **Biological Process** category (*red*, **Fig. 4)**, the top result, chromatin organization (GO:0006325, n=15), is in line with phase separation observed during nucleosome assembly and the formation of chromatin loops ^69^. Terms enriched across multiple organisms are also related to intracellular organization such as cellular component organization (GO:0016043, n =14) and chromosome organization (GO:0051276, n=12). Three terms are related to metabolic processes such as nucleic acid metabolic process (GO:0090304, n=14), mRNA metabolic process (GO:0016071, n=13) and DNA metabolic process (GO:0006259, n=13). More surprisingly, five terms are related to intercellular phenomena such as tissue development (GO:0009888, n=12), anatomical structure morphogenesis (GO:0009653, n=12), regulation of developmental process (GO:0050793, n=11), embryo development (GO:0009790, n=11), and developmental process (GO:0032502), suggesting a yet-to-be discovered potential role of LLPS in developmental processes.

### Phaseek suggests evolutionary conservation of LLPS among orthologs

Little is known about the conservation of phase separation across the tree of life^70,71,72^. To assess whether LLPS is conserved among orthologs, we analyzed LLPS scores among orthogroups which are sets of sequences derived from a single ancestral gene. Our hypothesis was that, if phase separation is a conserved trait, scores should remain consistent within the orthogroup, with variations arising solely by evolutionary divergence among orthologs^70^.

We categorized manually curated sequences from SwissProt using OrthoFinder^73–75^, identifying 24,438 orthogroups. Of these, only 216 included contributions from all 18 organisms (**Supplementary Fig. 53**), which is coherent with the number of sequences available across organisms (**Supplementary Fig. 11**). Noteworthy, one species can encode multiple paralogs corresponding to the same orthogroup. *E. coli* exhibited the lowest fraction of proteins with predicted phase-separating regions (2%), consistent with the observation that eukaryotes have more disordered proteins than prokaryotes^76^. In contrast, *S. cerevisiae* and *A. thaliana* each reached 12%, and *H. sapiens* 27% (**Fig. 5a**).

**Fig. 5.**
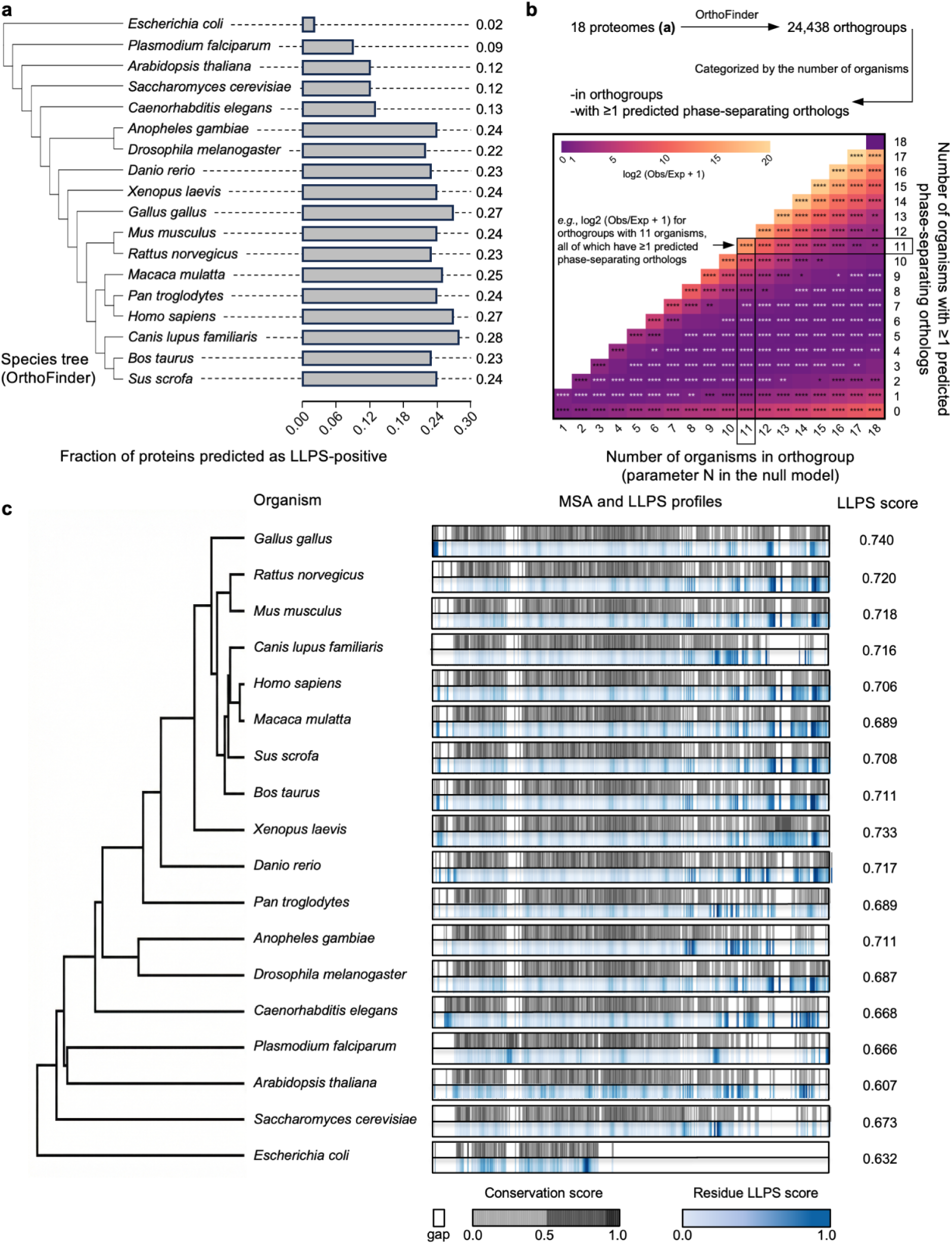
LLPS is conserved among orthologs. **(a)** Fractions of proteins predicted to be phase-separating by Phaseek are shown along with the species tree generated by OrthoFinder^73–75^. **(b)** OrthoFinder was applied across the 18 proteomes, identifying 24,438 orthogroups with representatives from 1 to 18 organisms. To evaluate whether LLPS propensity is conserved beyond chance, a binomial null model was applied in which each protein has an independent probability of being LLPS-positive equal to the global average LLPS rate across all proteins predicted by Phaseek. For each cell defined by orthogroup size (number of organisms, x-axis) and number of organisms with ≥1 LLPS-predicted protein (y-axis), the expected number of orthogroups under this null hypothesis was computed, and enrichment was quantified as log₂(observed/expected). Statistical significance was assessed using two-sided exact binomial tests with Benjamini–Hochberg FDR correction. The heatmap displays enrichment (pink/dark) or depletion (light) relative to the null, with asterisks denoting cases that significantly deviate from expectation after FDR correction. **(c)** Conserved LLPS propensity in Topoisomerase II orthologs across diverse species. Phylogenetic tree of 18 Topoisomerase II orthologs spanning major eukaryotic and prokaryotic lineages, constructed from multiple sequence alignment using biopython. Each row shows the residue-level LLPS scores predicted by Phaseek under the corresponding conservation scores. Despite divergence in amino acid sequence, particularly in the C-terminal domains (CTDs), several orthologs retain high LLPS propensity (Phaseek-predicted LLPS score > 0.7), suggesting that phase separation may represent a conserved functional feature of this enzyme family. Notably, orthologs in *S. cerevisiae* and *H. sapiens* have been experimentally validated to undergo LLPS, supporting the predictive relevance of the Phaseek model. Conservation, gap positions, and residue-level LLPS scores are shown as indicated in the color bars.

We compared the number of organisms having at least one LLPS-predicted ortholog to a null model (**Fig. 5b**). The null model uses the LLPS frequency across all predicted proteins as the probability for an organism to have an LLPS-predicted ortholog and a binomial law to compute the expected number of positive orthologs when considering N organisms. We evaluated enrichment or depletion in LLPS positive proteins using the log₂(observed/expected+1). Orthogroups in which all or none of the member organisms contain LLPS proteins are significantly enriched, as reflected by high scores along the diagonal and lower horizontal margin of the heatmap, respectively. Conversely, intermediate tiles, where only a subset of organisms possess an LLPS-positive ortholog, are significantly depleted. This pattern indicates that LLPS propensity tends to be either broadly shared or uniformly absent across organisms within orthogroups, suggesting that phase separation is a functionally conserved property throughout evolution. Additionally, we tested for over-represented species among orthogroups in which only one organism was absent and among those in which only one organism was present and found cases of such outlier organisms, namely *D. rerio* and *C. familiaris* (**Supplementary Fig. 54a and 54b**). Further, variations in phase separation propensity among positively enriched orthologs could be missed, for example, the BuGZ protein of *D. melanogaster* forms hydrogel-like material unlike its orthologs which are liquid condensates^70^.

Among the orthogroups with contributions from all 18 organisms, the topoisomerase II enzyme family (Topo II) included 14 members with LLPS scores > 0.7. Topo II resolves tensions from DNA supercoiling and entanglements^77^. In *S. cerevisiae* and *H. sapiens*, Topo II (TOP2α/TOP2β) undergoes LLPS via its less-conserved disordered C-terminal domain (CTD) to couple catenation with chromatin condensation^77,78^. Multiple sequence alignment further illustrates how phase-separating CTDs can retain high LLPS scores despite low conservation scores, suggesting that LLPS behavior can be conserved despite changes in the amino acid sequence (**Fig. 5c**).

LLPS scores among members of an orthogroup are coherent and suggest evolutionary functional conservation of the phenomenon across orthologs, across different sequence solutions.

### Phaseek generates de novo phase-separating peptides

The emergence of LLPS as a fundamental mechanism for spatial organization has opened new avenues in synthetic biology, where engineered membraneless organelles can be used to sequester cellular pathways or to create responsive intracellular compartments^79,80^. Naturally occurring LLPS sequences are often long, repetitive, or strongly context-dependent, which constrains their use as standalone engineering parts. Short peptides are particularly attractive in this context, as they can serve as modular tags for organizing proteins without the structural complexity of full-length sequences.

To meet this need, we expanded the Phaseek functionality from scoring existing sequences to generating new ones. We implemented a differentiable sequence optimization approach, with Phaseek as the scoring function (**Methods**). The first token of each peptide was constrained to be a methionine to mimic natural protein initiation, while the remaining positions were iteratively adjusted to maximize the LLPS score. By varying the target length, we generated a set of 20 candidate peptides ranging from 25 to 80 amino acids, all with high predicted LLPS propensity for experimental validation (**Supplementary Table 4**).

To visualize phase separation *in vivo*, we fused the short polypeptides to the C-terminus of super-folder GFP (sfGFP) and expressed them in *E. coli* under an inducible promoter (**Fig. 6a**). Using fluorescence microscopy, we observed that 14 out of the 20 tested candidates formed distinct condensates, achieving a 70% success rate (**Fig. 6b**). The 14 condensate-forming peptides produced visibly distinct phenotypes in cells. Some peptides drove the formation of one or two foci at the cell pole (*e.g.*, Phaseek_Gen_4, _9), or middle (*e.g.*, Phaseek_Gen_6, _18), a pattern characteristic of nucleoid-exclusion in *E. coli*^81,82^. A second group exhibited peripheral phenotype (*e.g.*, Phaseek_Gen_7, _8), suggestive of distinct localization or material behavior.

**Fig. 6.**
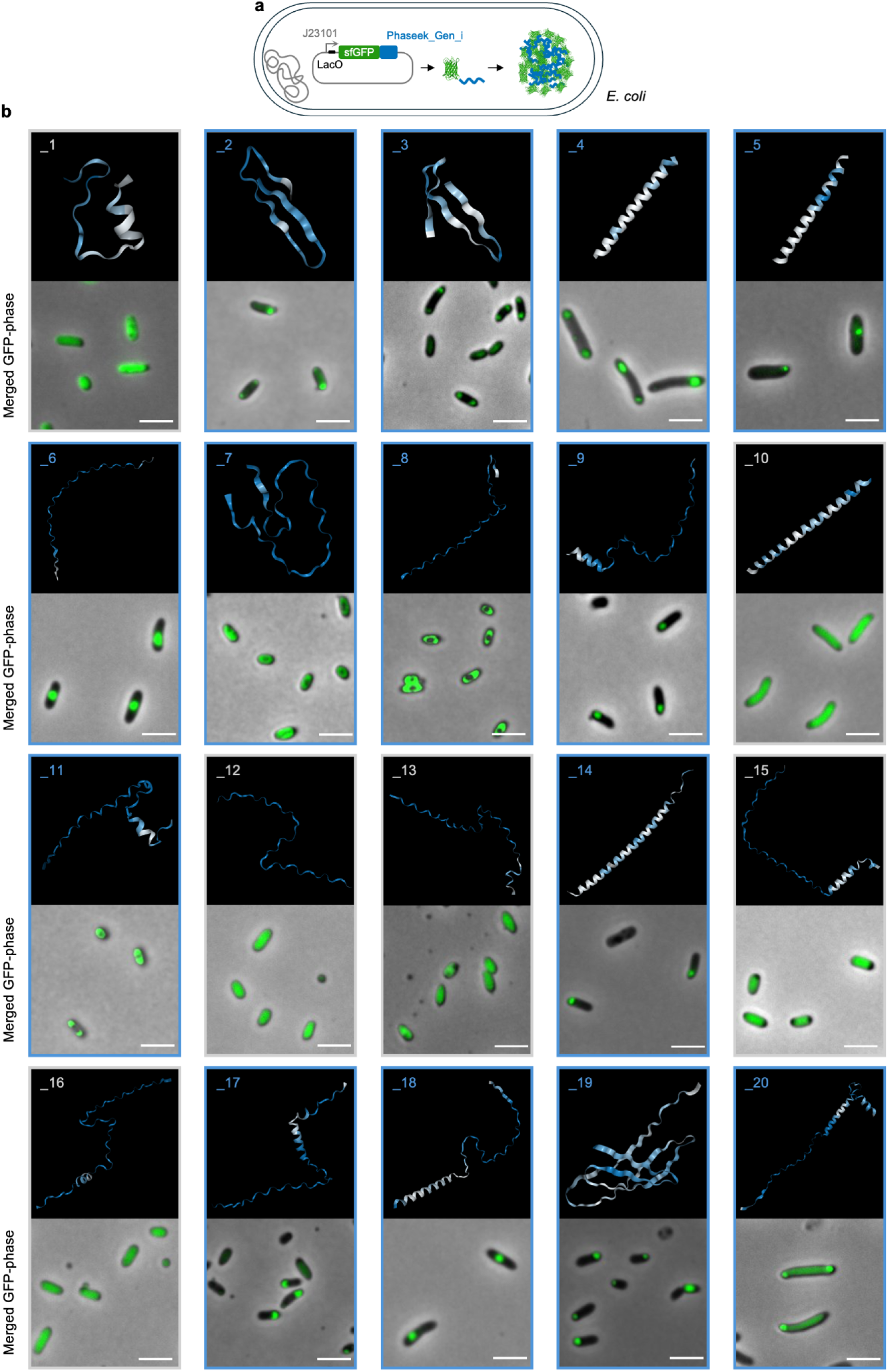
Experimental validation of de novo-designed phase-separating peptides. **(a)** Schematic representation of the genetic constructs in *E. coli* producing sfGFP with C-terminus fusion of each of the designed peptides. The fusion construct is expressed under a J23101 sigma-70 promoter followed by a LacO operator site. **(b)** Alphafold 3-predicted structure for each peptide where the color gradient shows the residue-level score (white=0, blue=1) for each of the 20 peptides along with merged phase and GFP images. The IDs of phase-separating peptides are highlighted in blue. Phaseek_Gen_1, _10, _12, _13, _15, and _16 did not show a phase separation phenotype (IDs in white).

We predicted the structure of active peptides using AlphaFold and observed a diverse structural landscape including both ordered and disordered regions, such as β-sheets (Phaseek_Gen_2, _3, _19), single α-helical rods (Phaseek_Gen_4, _5, _14), disordered (Phaseek_Gen_6, _8), and mixed helical and disordered conformations (Phaseek_Gen_9, _11, _17, _18, _20). This is in line with Phaseek ability to recognize LLPS across different structures and with prior reports that LLPS is not restricted to disordered proteins (**Fig. 1g**). This phenotypic diversity is relevant for synthetic biology applications, since condensate properties, including material state, morphology, and localization, are recognized as key determinants of condensate function^83^. The smallest active candidates, 25 amino acids of non-repetitive sequences, minimize the structural burden imposed on fusion partners, giving a practical advantage over longer or repetitive LLPS scaffolds.

Together, these results show that Phaseek can be used to design phase-separating sequences. The resulting peptide repertoire offers a versatile toolkit for engineering biomolecular condensates with applications in synthetic biology and biotechnology.

## Discussion

We introduced Phaseek, a sequence-based LLPS prediction model that integrates transformers architecture, feature-specific matrices, and a region-boosting bottleneck to compute sequence and residue level scores (**Fig. 1a, Supplementary Fig. 2**). It must be noted that the scores produced by the model are not indicative of condensate properties such as size, dynamics, or critical concentration but rather reflect the likelihood that a sequence harbors phase-separating region(s) and may phase-separate under physiological conditions. We demonstrated that Phaseek produces sequences and residue level scores with high accuracy in various biological contexts making it a versatile and reliable tool for the prediction of LLPS proteins.

Phaseek identifies key regions driving phase separation (**Fig. 2a, b, c**) despite being trained on amino acid sequences without explicit annotation of subsequences. We showcased this feature on the ORF1 protein of the human transposable element LINE-1, for which the model captured the effect of truncating two key regions. Inversely, we showed that fusion of phase-separating peptides to non-phase-separating proteins significantly increases the score of the original sequence (**Supplementary Fig. 4c and d**). The ability of the model to detect key regions makes it a valuable tool for hypothesis-driven study of LLPS proteins and could be used for example, to identify proteins of the endogenous retrovirus HERV-K113 driving phase separation as its gag polyprotein sequence, which codes for elements of viral particles, is predicted to phase separate (**Supplementary Fig. 55**).

Phaseek predicts the effect of point mutations and whether these promote or weaken LLPS, as exemplified for two mutations on the structure of human Annexin A11 (**Fig. 2d**). Among the 2,529 mutations studied, amino acid substitutions from polar to aromatic, particularly serine to phenylalanine and serine to tyrosine, resulted in the highest score increase while aromatic to hydrophobic substitution, especially tyrosine to isoleucine, were associated with the largest decrease of LLPS propensity (**Supplementary Fig. 6**). These changes in score correspond to substitutions involving aromatic residues suggesting that the model captures the importance of π-effects (cation–π and π-stacking) in phase separation. Mapping LLPS scores on protein structures predicted by AlphaFold revealed that the model assigns greater scores to unstructured and exposed regions. Residue level scores for point mutation, such as Annexin A11 R346C, suggest that single amino acid substitutions can impact surrounding regions relatively distant in both the primary sequence and the 3D structure (**Supplementary Fig. 10**).

The proteome-wide predictions of LLPS proteins for 18 organisms highlight physicochemical properties captured by the model and their respective importance in key regions. In line with the role of intrinsically disordered regions in phase separation, the model predictions are well correlated with the IDR score which is highly predictive of key regions (**Fig. 3b and d**). GO terms enriched for the LLPS positive fraction and shared across organisms recapitulate LLPS-prone processes, such as chromatin binding, transcription factor binding and chromatin organisation (**Fig. 4**), supporting a fundamental role of phase separation across the tree of life. Phylogenetic analyses suggest the evolutionary conservation of phase separation property within orthogroups (**Fig. 5b**) and a more detailed analysis of these sequences suggests that this conservation goes beyond sequence similarity (**Fig. 5c**).

Finally, Phaseek can be used to design de novo LLPS peptides. For 20 candidates between 25 and 80 amino acids in length, we experimentally confirmed the phase-separating property of 14 peptides, achieving 70% success in vivo (**Fig. 6**). These diverse peptides (**Supplementary Table 4**) showed different phenotypes of phase-separation, *i.e.*, diverse shapes and sizes and expectedly fluidity/viscosity. Interestingly, this list includes both disordered and folded peptides as we show that Phaseek is able to distinguish them in the context of LLPS (**Fig. 1g**). These provide a versatile repertoire of phase-separating peptide tags for various applications in biochemistry, synthetic biology and biotechnology, including protein purification, pathway sequestration, programmable on/off systems, and delivery platforms^84^.

Phaseek serves as a multi-scale, multi-purpose model enabling researchers to predict LLPS proteins, as well as the sub-sequences driving LLPS, or the effects of mutations at the amino acid resolution. Phaseek can also be used to generate de novo LLPS peptides with different lengths and structures and associated with distinct LLPS phenotypes.

## Methods

### Train, test and evaluation data

To construct a dataset of LLPS positive sequences, we gathered multiple available resources, including LLPSDB^85^, DrLLPS^86^, PhaSeDB^87^, and PhaSePro^88^, along with a manually curated set of 611 sequences from the literature, yielding a total of 1,275 LLPS-positive polypeptides (**Data availability**). For negative data, we used a set of 5,258 sequences previously compiled for PSPredictor^28^.

We filtered out sequences longer than 512aa, exceeding the model capacity to capture long-range dependencies, and applied CD-HIT^89^ (Cluster Database at High Identity with Tolerance) clustering with a similarity threshold of 0.9 to eliminate highly similar sequences, resulting in a set of 659 positive and 4,859 negative sequences (**Data availability**).

Due to the limited number of LLPS positive sequences, we employed an oversampling technique to augment the size and diversity of our training set. IDRs are widely recognized as important drivers of LLPS^17^. To preserve these regions during data augmentation, we introduced mutations only within the structured (folded) regions of protein sequences, leaving disordered regions intact. To do so, we trained a categorical HMM (hidden Markov model) on a synthetic set of folded regions comprising residues with AIUPred IDR scores inferior to 0.5, and considered as folded, from the set of LLPS positive sequences using the Python hmmlearn library (https://github.com/hmmlearn/hmmlearn). The trained HMM was used to sample new amino acid sequences of the same length as each original sequence. We repeated the augmentation process multiple times, ultimately increasing the size of the positive set from 659 to 4,151 sequences. Finally, the positive and negative training data sets were splitted with a 9:1 ratio.

The dataset used for evaluation included positive sequences which were not used during training nor tuning, as well as synthetic sequences compiled from five studies^37–41^. For negative sequences, the evaluation set comprised sequences from the training split, supplemented with the PSPHunter^26^ negative test set to increase diversity.

### Model architecture and training

Our model consists of two core modules, an encoder and a protein graph generator. The encoder module takes as input a protein sequence, encoded as an array of one-dimensional token indices, and outputs an array of token embeddings. It is built based on a classical GPT architecture with 6 attention heads per layer and 6 layers in total. We employed a token embedding size of 192 and a block size of 512. The protein graph module operates in parallel with the encoder to generate graphical space representations of protein sequences. Each sequence is encoded into 10 distinct statistical matrices based on sorting methods derived from the FEGS paper^31^ (see Matrix selection and Protein graph sections below). After padding, these vectors are merged via weighted summation, producing a single extended matrix. In the final pooling step, outputs from the encoder and protein graph modules are integrated. The resulting output is then passed through a multi-layer perceptron model which reduces the dimensionality of each token, followed by a median operation to yield a 2-dimensional probability vector (Supplementary Methods). The integrated core model, comprising the encoder and the protein graph generator, was trained on the augmented training set (see Train, test and evaluation data section). Optimization was performed using the AdamW optimizer with the cross entropy loss function applied to quantify prediction errors. The model demonstrated convergence after 35 epochs, achieving the minimum test loss at this point (**Code availability**).

### Matrix selection

Each protein sequence in our dataset was encoded by the FEGS method^31^ into 158 matrices, whose eigenvalues were concatenated into a 158-dimensional feature vector. A random forest classifier was subsequently trained on the training set based on the resulting vectors (**Code availability**). To evaluate the contribution of each of the 158 matrices in the scoring process, we employed the SHAP method, a game theoretic approach for interpreting the outputs of machine learning models^32^. This analysis enabled us to quantify the relative contribution of each matrix by calculating its SHAP value. Based on this, we identified and selected the ten most predictive matrices during the scoring process (**Supplementary Table 1**).

### Score boosting module

We introduced a boosted scoring mechanism to better capture the contribution of key regions to LLPS. A gradient boosting classifier (XGBoost) was trained on residue-level LLPS scores generated by the core model to leverage local phase separation propensities.

The final score is obtained by combining the scores of the core model and the XGBoost bottleneck module. This strategy enhances the accuracy of LLPS prediction by identifying and emphasizing the contributions of key regions involved in phase separation (**Supplementary Fig. 1**).

### Other LLPS predictors used in this study

PSPHunter is a random forest model that utilizes 60 sequence-derived features including RNA-DNA binding site affinity and post-translational modification sites, and is specialized in identifying essential key residues in LLPS^26^. catGRANULE 1.0 is a statistical algorithm that considers the contributions of structural disorder, nucleic acid binding propensities, sequence length, and amino acid content^35^. catGRANULE ROBOT 2.0 can predict LLPS at single amino acid resolution using a multi-layer perceptron model based on physicochemical and structural features^27^. PScore is a machine learning model trained to specifically capture pi-pi and pi-cation interactions that promote LLPS^36^. PSPredictor is a word2vec-coded model trained using a gradient-boosting decision tree^28^. Finally, PSTP (with two versions available) is a multi-layer perceptron model that uses the ESM-2 protein language model^90^, and ALBATROSS IDR prediction model^91^ to encode the input sequences^29^.

### Evaluation parameters

To evaluate the model performance and interpret the results, we employed several standard metrics (TN, TP, FN and FP denote true negative, true positive, false negative, and false positive, respectively):

- AUC (area under the ROC curve) assesses the ability of the classifier to differentiate positive from negative classes regardless of the classification threshold. Higher AUC indicates better performance.

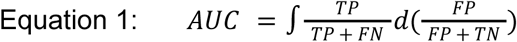

- Recall measures the capability of the model to identify the actual positive cases. It is defined as the proportion of true-positives out of all actual positive cases.

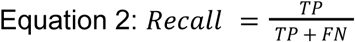

- Precision, or positive predictive value, measures the proportion of predicted positive cases that are actually positive.

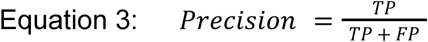

- F1-Score combines precision and recall into a single metric, offering a balanced measure of performance, especially useful when dealing with imbalanced data.

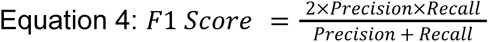

### Confidence intervals

Using our validation set, we swept decision thresholds on Phaseek scores and selected cut points where the empirical precision (PPV) of scores met predefined target thresholds, thereby defining the “Possible,” “Probable,” and “High-confidence” bands. For each band, we report the score range, sample size, and observed precision with 95% binomial confidence intervals, and re-estimate thresholds periodically to track shifts.

|  | lower_inclusive | upper_inclusive | n_samples | empirical_precision |
| --- | --- | --- | --- | --- |
| Possible | 0.2752 | 0.5557 | 132 | 0.28 |
| Probable | 0.5555 | 0.8890 | 213 | 0.87 |
| High-confidence | 0.889 | 0.967 | 63 | 1.00 |

### Validation with folded/disordered (non-)phase-separating data

To identify folded phase-separating (folded PSPs) and disordered non-phase-separating (disordered-nonPSPs), we utilized AIUPred to calculate the average disorder score associated with each protein sequence across DrLLPS, PhaSePro, and PhaSeDB databases. Using a strict cut off of 0.7 on the average IDR score from AIUPred, we filtered out the non-phase-separating sequences. For Folded-nonPSPs we used a threshold of 0.3 and selected only the phase-separating sequences with IDR score below 0.3. Sequences overlapping with the train/test sets or failing CD-HIT filtering were excluded from further analysis.

### Validation with mutation data

We collected data for 2,328 mutations annotated as missense mutations from the DisPhaseDB⁶⁶ database. These mutations influence LLPS in disease-associated proteins, although they have not been specifically classified as either LLPS-strengthening or LLPS-weakening in the database. Additionally, we gathered 201 mutations, labeled as LLPS-strengthening (n = 90) or LLPS-weakening (n = 111) from DrLLPS, PSMutPred^54^ and literature^45,55,92–97,98^. This resulted in a total of 2,529 determining mutations corresponding to 137 distinct proteins.

For the background mutations used in comparative analysis, 2,740 random mutations were generated by introducing 20 random amino acid substitutions per sequence across the same 137 proteins. To assess the ability of Phaseek to distinguish determining mutations from random ones, all mutations including the 2,328 missense, 90 LLPS-strengthening, and 111 LLPS-weakening substitutions were compared against the random mutation set. To evaluate whether Phaseek can differentiate LLPS-strengthening from LLPS-weakening mutations, only the 90 strengthening and 111 weakening mutations were used.

Three distinct scores were defined to evaluate model performance to detect determining mutations:

1. Δ(score): The difference in LLPS score between the mutated sequence and the corresponding wild-type sequence.
2. Δ(aa_score): The change in LLPS score for the single mutated residue before and after mutation.
3. Σ(Δ(aa_score)): The cumulative LLPS score difference across all residues whose individual LLPS propensity values changed as a result of the mutation, calculated only for positions where the absolute score difference exceeds the threshold of 1e-4.

Human Annexin A11 (P50995 · ANX11_HUMAN)^56,99^ was selected to illustrate the effect of point mutations on residue-level scoring (**Fig. 2, Supplementary Fig. 11**) and one strengthening and one weakening mutation were used for illustration.

### Residue-level LLPS profile of proteins

We implemented a sliding window whose size varies based on protein length, ranging from a minimum of 5 to a maximum of 50 amino acids. This window slides across the full sequence, centering on each residue so that the target amino acid of interest lies at the midpoint. Subsequently, the window is divided into segments, with a length of half the window size. Finally, the LLPS scores for each segment are calculated by the core model and processed by a weighted summation layer to produce the final residue-level score. Weights are generated and assigned using an adjusted Gaussian distribution, giving higher importance to residues closer to the central amino acid (**Supplementary Fig. 1**).

To assess the models ability to accurately distinguish LLPS-promoting regions from those with low-propensity, we defined a custom match scoring function focused on two key criteria: (i) the model accuracy to classify regions as LLPS-promoting or low LLPS propensity, and (ii) its ability to produce a significant difference in scores between these two region types.

Residues were classified as LLPS-positive or LLPS-negative based on a score threshold of 0.5. The corresponding accuracy score was calculated according to the experimentally validated LLPS profiles at the residue level.Additionally, to validate the significance of the differences in assigned LLPS scores, we calculated the average score difference between LLPS-promoting and low-propensity regions. Finally, the mean of the two scores was computed.

### Key region and random region derivation

To define the key regions, we first smoothed the residue-level scores of each sequence with a sliding window of size 7. Key regions were defined as consecutive segments where the smoothed scores remained at or above 0.7 for a minimum of 20 residues. If the average score within any 7-residue window dropped below 0.7, the region was considered discontinued.

Random regions, ranging from 20 to 250 residues in length, were sampled from each protein sequence across the proteomes.

### Physicochemical features

A score for aromatic clustering was calculated as in Holehouse et al.^100^

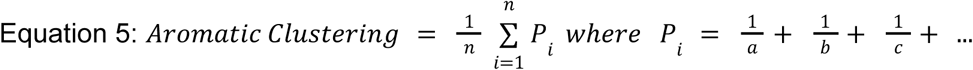

a, b, and c are the distances between aromatic residues. Proteome sequences with no aromatic residues present were excluded from the analysis.

Disorder scores were calculated using AIUPred^58^.

We used localCIDER^101^ to calculate kappa (*κ*)^102^, hydropathy, polyproline II helix (PPII) propensity, net charge, and amino acid fractions in different categories including negative residue fraction (negative), positive residue fraction (positive), glycine and proline residue fraction (Gly/Pro), polar residue fraction (polar), aromatic residue fraction (aromatic), and hydropathic residue fractions (hydrophobic).

The charge blockiness parameter Kappa (*κ*)^102^ quantifies the extent of charged amino acid mixing (R, K, D, E) and is defined as

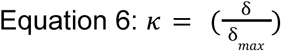

where δ defines the average deviation from charge symmetry across sequence regions (blobs) using a sliding window (size of 5), and δ_*max*_ is the maximum δ for the sequence given its composition. A *κ* value of 0 indicates charge disparity, while 1 represents maximum charge blockiness. A negative *κ* value (*e.g*., −1, see the minor peak in **Fig. 4d**) corresponds to uncharged sequences.

Hydropathy is based on the Kyte-Doolittle hydrophobicity scale, where each amino acid is assigned a hydropathy index reflecting its water-vapor transfer free energy change and its side-chain distribution between the protein interior and exterior^103^.

PPII helix propensity is based on the Hilser scale, whereby each amino acid is assigned a polyproline propensity score measured using a peptide host-guest system and isothermal titration calorimetry^104^.

### Secondary structure derivation

Predicted structures of proteins for 10 proteomes were obtained in PDB format from the AlphaFold Protein Structure Database (https://alphafold.ebi.ac.uk/download). Secondary structure information was extracted using the DSSP (Dictionary of Secondary Structure in Proteins) package^105,106^. Structural elements were categorized into five groups: helices (H, G, I, P), sheets (B, E), turns (T), bends (S), and irregular or flexible regions (-), with the latter classified as disordered regions (**Supplementary Table 2**).

### Functional enrichment

To associate the LLPS phenomenon with biological processes, we enriched LLPS-positive proteins for Gene ontology terms, which hierarchically represent biological processes. We reduced the list of results using semantic similarity and compared the filtered terms across species.

We retrieved the proteomes predictions of all organisms but *Plasmodium falciparum*, whose annotation package is not maintained (**Supplementary Table 3**) and considered both manually curated sequences from SwissProt and computationally predicted sequences from TrEMBL database. For each organism, we filtered proteins with an LLPS score equal or higher than 0.7 (distributions of LLPS scores and percentages of the proteome selected for each organism; **Supplementary Fig. 35-51**).

To map genes and GO terms, we used for each organism the corresponding OrgDb package containing versioned implementations of the GO processes to ensure the full reproducibility of the results (**code availability**). We then performed simple hypergeometric tests as implemented in the gostats^107^ package (v2.70.0) (top-10 GO terms enriched for each organism represented as lollipop plots; **Supplementary Fig. 35-51**).

To summarize the list of enriched terms, we applied semantic similarity using the rrvgo^108^ package (v1.16.0) with default method (Rel) and threshold (0.7). This package mimics the core functionality of the REVIGO webserver ^62^ which reduces the redundancy within a list of GO terms (for each organism, summarized GO terms are represented as treemaps; **Supplementary Fig. 35-51**). The results are provided as a notebook together with the code to reproduce this analysis (**Data availability**).

### Ortholog analysis

The coding sequences (CDS) of 18 organisms, including *E. coli*, *P. falciparum*, *S. cerevisiae*, *A. thaliana*, *A. gambiae*, *D. melanogaster*, *C. elegans*, *D. rerio*, *X. laevis*, *G. gallus*, *R. norvegicus*, *M. musculus*, *C. lupus familiaris*, *B. taurus*, *S. scrofa*, *M. mulatta*, *P. troglodytes*, and *H. sapiens,* were retrieved from UniProt proteome database (**Accession IDs and versions listed Supplementary Table 3**).

For all organisms, we used a unique manually curated sequence from SwissProt for each gene. The LLPS scores of these sequences were predicted by Phaseek and assigned as phase-separating with a threshold of 0.7. Using OrthoFinder^73–75^, 328,887 genes (93.7% of the total) were grouped into 24,438 orthogroups. Half of the genes belonged to orthogroups containing 18 or more genes (G50 = 18) (see **Data availability**). Among these, 216 orthogroups contained sequences from all species, including 9 orthogroups composed exclusively of single-copy genes. The tree inferred by orthofinder in **Fig. 3a** was generated using Dendroscope^109^.

To assess the statistical significance of LLPS conservation across orthogroups, a binomial null model was employed. Each protein was assumed to have an independent probability *P* of being LLPS-positive, where *P* represents the global average LLPS rate across all proteins in the dataset based on Phaseek predictions. Under this null model, the probability that an orthogroup considering *N* organisms contains exactly *k* LLPS-positive members follows a Binomial(*N*, *P*) distribution. For each (*k*, *N*) combination, the expected number of orthologs under the null was calculated, and enrichment was quantified as log2(Observed/Expected + 1). A two-sided exact binomial test was then applied to determine whether the observed count in each tile significantly deviated from the null expectation. All p-values were corrected for multiple testing using the Benjamini–Hochberg false discovery rate (FDR) procedure. Results were visualized as a heatmap in which color intensity reflects the degree of enrichment or depletion relative to the null model, and asterisks denote cells with statistically significant deviations after FDR correction.

### LLPS investigation in Topoisomerase II enzyme orthologs

We explored the orthogroups with members from all 18 organisms to identify protein sequences with conserved LLPS properties across the tree of life. Only orthogroups containing a minimum of nine LLPS-positive members were selected, resulting in 45 candidates. Among these, the Topoisomerase II family stood out (min = 0.64, max = 0.76, median = 0.73). Given the fundamental role of this enzyme in resolving DNA supercoiling and genome organization during transcription and replication, we investigated whether the elevated LLPS scores might reflect a conserved role for phase separation in Topoisomerase II function. To investigate further, we conducted a literature search and found that Topoisomerase II orthologs in two evolutionarily distant organisms, *Saccharomyces cerevisiae* and *Homo sapiens*, have been shown to undergo phase separation^77^.

### Phase separating peptide generation

To design novel peptides with high predicted LLPS propensity, we employed a differentiable sequence optimization strategy inspired by SeqProp (https://github.com/johli/seqprop)^110^. This framework enables gradient-based optimization of amino-acid distributions using a pretrained Transformer classifier as the scoring function.

Sequences were modeled using a logit matrix 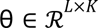 where L is sequence length and K=20 amino acids. To reflect natural initiation, the first position was fixed to methionine (M), while the remaining logits were randomly initialized. Discrete amino-acid choices were relaxed into a differentiable probability matrix P using a temperature-controlled Gumbel–Softmax:

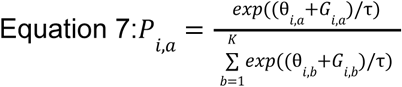

where *G_i,a_*, denotes Gumbel noise and τ is the annealing temperature. This relaxation enables gradient flow through sequence choices while still converging to discrete solutions. The continuous sequence representation was projected into the model embedding space via:

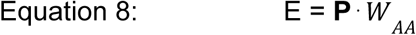

Where *W_AA_* is the frozen amino-acid embedding matrix from the Transformer. The resulting embedding was fed through our pre-trained classifier to obtain a differentiable LLPS score.

Sequence parameters were optimized to maximize the predicted LLPS score while penalizing high-entropy distributions:

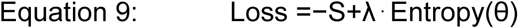

where λ controls the sharpness of the emerging sequence profile. Optimization was performed using gradient ascent with Adam optimizer. The temperature τ was linearly decreased from a high initial value to a low final value to support early exploration and late-stage convergence.

Final peptide sequences were obtained by argmax decoding across amino-acid probabilities. An optional discrete refinement step (local hill climbing) was used to further maximize model-predicted LLPS propensity. The Gumbel–Softmax temperature was linearly annealed from 2.0 at the beginning of optimization to 0.1 at the final iteration. The initial Gumbel-noise scale was set to 1.0 and was also linearly reduced to zero, allowing stochastic exploration during the early iterations and increasingly deterministic sequence selection toward the end.

### Plasmid construction and DNA preparation

The DNA fragments encoding the 20 de novo peptide and the necessary primers were synthesized by Integrated DNA Technologies (IDT). The constructs were built under the control of the IPTG-inducible sigma 70-based promoter J23101 from the iGEM collection in a plasmid with pMB1 origin of replication and kanamycin resistance gene. To be able to visualize condensate formation, the candidate peptides were fused to the C-terminus of superfolder GFP (sfGFP) via a GGS linker. Cloning was performed using Golden Gate assembly (BsaI-HF®v2 NEB #R3733L, T4 DNA ligase NEB #M0202T, T4 DNA Ligase Reaction Buffer 10X NEB #B0202S, and DpnI NEB #R0176L). PCR amplifications for cloning inserts were performed using Q5® High-Fidelity 2X Master Mix (NEB #M0492L). Following amplification, DNA fragments were purified using the Monarch PCR & DNA Cleanup Kit (NEB #T1030L). Correct assembly was verified by sequencing. For downstream experiments, plasmids were purified using the Monarch® Spin Plasmid Miniprep Kit (NEB #T1110L).

### Sample preparation and fluorescence microscopy

*E. coli* strain TOP10 was used for all imaging experiments. Cells transformed with the expression plasmids were cultured in liquid LB medium supplemented with kanamycin at 37°C. Upon reaching an optical density at 600 nm (OD600) of 0.4–0.5, protein expression was induced by the addition of 0.1 mM IPTG. The cultures were subsequently incubated overnight at 37°C with shaking. For imaging, 2 µL of the overnight culture was dispensed onto a microscope slide and covered with a 1% agarose pad to immobilize the cells. Imaging was performed using a Zeiss Axio Observer 7 inverted microscope equipped with a 100x oil immersion objective. The fluorescence of the sfGFP-fused peptides was recorded using an excitation wavelength of 475 nm. Image processing and analysis were performed using Fiji software.

## Data availability

The training and test data is available on https://github.com/AMIRMOHAMMAD-OSS/Phaseek, along with the detailed pathway enrichment analysis data for each organism (associated with **Fig. 3c**). The predicted LLPS score of the 18 proteomes (associated with **Fig. 3, 4**), physicochemical and structural data (**Fig. 4, Supplementary Fig. 15-18**) are available on https://zenodo.org/records/16728261. Data underlying **Fig. 1b-g**, **Fig. 2a-e**, **Fig. 3c**, **Fig. 4b**, **Supplementary Fig. 2-4**, **7-13** are provided as **Source Data File**.

## Code availability

The code associated with the model, along with the trained model, is provided on https://github.com/AMIRMOHAMMAD-OSS/Phaseek with a license for academic and non-commercial use. The code for pathway enrichment analysis is also available on the same GitHub repository. A user-friendly Google Colab notebook of Phaseek is available through this [link](https://colab.research.google.com/github/AMIRMOHAMMAD-OSS/Phaseek/blob/main/phaseek_colab.ipynb).

## Acknowledgments

This work was supported by ATIP-Avenir young group leader program by INSERM-CNRS, Emergence Sorbonne University grant, and MOPGA (Make Our Planet Great Again) young researcher fellowship by the French government (AP), and the Fondation Bettencourt Schueller (AP and ABL). The authors wish to thank Mostafa Elraies, Negin Hadisadegh, and Roxana Ghasemi for their help in collecting initial training data and Amir Reza Aliakbari and Sadrodin Barikbin for revising the code developed in this work.

## Author contributions

AMM developed the models. AMM and HT gathered data and results and together with AP and VG analyzed the results. VG performed enrichment analyses together with HM. AMC, JB, DAR and YA performed wet lab experiments. AMM and HT prepared the draft of the manuscript with contributions from all authors. RN, ABL, VG and AP supervised the work.

