## Supplementary information for "A hybrid model for multiscale prediction and generation of phase separating protein regions"

**FEGS graphs: from sequence to pairwise geometry**

Given a protein sequence s=($s_1$,..., $s_L$) with $s_i$ in the 20-letter amino-acid alphabet, the FEGS framework maps s into a family of motif-conditioned geometric walks in R3. From each walk it derives a pairwise matrix M ∈ $\mathbb{R}^{L\times L}$ that captures relative geometric relations between residue positions. Classical FEGS summarizes each matrix by a scalar (its dominant eigenvalue), yielding a fixed-length descriptor. In our work we used the pre-eigenvalue matrices themselves as graph-like objects.

**Motif set and anchor geometry**

Let $\{M_j\}_1^K$ denote a fixed set of motif alphabets over amino acids (we use K=158 one per physicochemical descriptor). Define 20 anchor points on the unit circle with a constant offset in the third coordinate:

$$P_a = (cos(\frac{2\pi i}{20}), sin(\frac{2\pi i}{20}), 1) \in \mathbb{R}^3 \qquad a = 0,\dots,19$$

Define micro-steps between anchors with a scale parameter $\gamma$>0 :

$$V_{ab} = P_a + \gamma(P_b - P_a) \in \mathbb{R}^3 \qquad a,b \in \{0,\dots,19\}$$

**Motif-conditioned geometric walk**

For each motif $M_j$, a 3D walk $C_j$ =($c_0^j$ ,..., $c_L^j$) is constructed with initialization $c_0^j$ = 0 and a momentum-like drift $d_i^{(j)} \in \mathbb{R}^3$ . At position i define a one-hot vector $x \in \{0,1\}^{20}$ over the motif alphabet: $x_a = 1$ iff residue $s_i$ matches anchor $a$ within $M_j$ ( else $x = 0$ ).
Drift update (encodes smooth transitions between consecutive matches):

$$d_i^{(j)} = \begin{cases} \frac{\mathrm{i}-1}{\mathrm{i}} d_{i-1}^{(j)} + \frac{1}{\mathrm{i}}(\mathrm{V}_{\mathrm{a(i-1),a(i)}} - \mathrm{P}_{\mathrm{a(i-1)}}), & \text{if x} \neq 0 \text{ and previous match y} \neq 0 \\ \frac{\mathrm{i}-1}{\mathrm{i}} d_{i-1}^{(j)} & \text{otherwise} \end{cases}$$

Position update:

$$\mathrm{c}_i^{(j)} = \begin{cases} \mathrm{c}_{i-1}^{(j)} + \boldsymbol{x}^{\mathrm{T}} \boldsymbol{P} + d_i^{(j)}, & \text{if x} \neq 0 \\ \mathrm{c}_{i-1}^{(j)} + (0,0,1) + d_i^{(j)}, & \text{if x} \neq 0 \end{cases}$$

We discard the initial origin and denote the resulting polyline by:

$$W^{(j)} = (c_1^{(j)},\dots,c_L^{(j)}) \in \mathbb{R}^{L\times 3}$$

**Pairwise matrix:**

From $W_j$ we compute:

1.Euclidean distance matrix

$$E_{uv}^{(j)} = ||W_u^{(j)} - W_v^{(j)}||_2 , \qquad 1 \le u,v \le L$$

2. Cumulative along-path distance(distance when traversing only adjacent residues):

$$\mathrm{S}_{\mathrm{uv}}^{(j)} = \begin{cases} 0, & \text{if u} = \text{v} \\ \sum_{\mathrm{t=u}}^{\mathrm{v}-1} \mathrm{E}_{\mathrm{t,t+1}}^{(\mathrm{j})} & \text{if u} < \text{v} \\ \mathrm{S}_{\mathrm{vu}}^{(\mathrm{j})}, & \text{if v} > \text{u} \end{cases}$$

Finally, the FEGS pairwise matrix is then calculated as :

$$L^{(j)} = E^{(j)} \oslash D^{(j)} \in \mathbb{R}^{L\times L} \qquad which\, D^{(j)} = S^{(j)} + I$$

where $\oslash$ denotes elementwise division. Intuitively, $L^{(j)}$ is large where residues are far in straight-line distance relative to their stepwise separation along the walk, and small where they are correspondingly close, encoding geometry induced by motif $M_j$ .(**Supplementary Fig.1**)
We set $\gamma \in$ {1/20,1/4} and use K =158. For a sequence of length L, forming $E^{(j)}$is O($L^2$ ); computations are parallelized across motifs/sequences. The addition $I$ in $D^{(j)}$ avoids division-by-zero on the diagonal and improves numerical stability.

**Attention with a FEGS-derived pairwise prior**
For each sequence, we compute the ten selected FEGS matrices, each representing pairwise relationships between residues derived from the corresponding motif-conditioned graph construction. Let

$$L^{(j)} \in \mathbb{R}^{n\times n}, \qquad j = 1, \ldots, 10$$

denote the j-th matrix for a sequence of length n. Each matrix is standardized over its valid, unpadded residue–residue region:

$$\tilde{L}^{(j)} = \frac{L^{(j)} - \mu\left(L_v^{(j)}\right)}{\sigma\left(L_v^{(j)}\right) + \varepsilon}$$

where Lv(j) denotes the valid n×n region and ε prevents numerical instability. The standardized matrix is then zero-padded to the model context size T×T, with T=512. In the current implementation, a matrix with near-zero variance is replaced by zeros rather than divided by an unstable standard deviation.
Rather than combining the ten matrices into a single shared prior, Phaseek learns a separate mixture for each attention head. For head h, the mixture weights are

$$\pi_{h,j} = \mathrm{softmax}_j\left(\frac{\alpha_j + \delta_{h,j}}{\tau}\right)$$

where α∈R10 contains global matrix logits, δh∈R10 contains head-specific offsets, and τ>0 is the mixture temperature. The resulting bias for head h is

$$B_h = \sum_{j=1}^{10} \pi_{h,j}\, \tilde{L}^{(j)} \in \mathbb{R}^{T\times T}$$

The default Phaseek v2 configuration uses a shared mixture across Transformer layers, meaning that the same head-specific mixture weights are reused in every block. The implementation also supports a layer-specific alternative in which the mixture weights depend on both layer and head.
Before being added to the attention logits, each head-specific bias is standardized again over valid residue–residue pairs:

$$\hat{B}_h = \frac{B_h - \mu(B_{h,v})}{\sigma(B_{h,v}) + \varepsilon}$$

This second normalization is performed within the graph-biased attention module and prevents differences in the scale of the mixed FEGS biases from directly controlling their influence on attention. Values outside the valid residue-pair region are set to zero.
For Transformer layer ℓ and attention head h, the pre-softmax attention logits are

$$A_{\ell,h} = \frac{Q_{\ell,h} K_{\ell,h}^{\mathrm{T}}}{\sqrt{d_k}} + \beta_{\ell,h} \hat{B}_h + M_{\mathrm{pad}}.$$

where βℓ,h is a learned head-specific scaling parameter within block ℓ. Thus, each attention head can independently control how strongly it uses the FEGS-derived pairwise prior relative to the sequence-derived query–key similarity.
Padded key positions are masked before the softmax operation. Outputs corresponding to padded query positions are subsequently multiplied by the valid-token mask and set to zero. Therefore, padded positions neither contribute as attended keys nor propagate nonzero representations through subsequent Transformer blocks.
This formulation is conceptually related to Graphormer[ref], which incorporates graph-structural information into a Transformer through attention biases. It is also related at a broader architectural level to AlphaFold2, where learned pair representations influence residue-level attention. ALiBi similarly demonstrates the general principle of modifying query–key attention logits with an additive pairwise bias, although its bias is based on linear sequence distance rather than sequence-conditioned FEGS features

**Supplementary Table 1. Top Ten FEGS Features Used for The Derivation of Statistical Matrices.** SHAP values were calculated using a random forest model and the top 10 matrices were subsequently selected for implementation in Phaseek.

| Index | FEGS Feature Index | Feature Name |
|---|---|---|
| 1 | 21 | Aperiodic indices for alpha/beta-proteins |
| 2 | 82 | Weights for coil at the window position of 5 |
| 3 | 57 | 8Acontact number |
| 4 | 30 | Average relative probability of inner helix |
| 5 | 155 | TOTFT index |
| 6 | 42 | Retention coefficient in $NaClO_4$ |
| 7 | 7 | alpha-CH chemical shifts |
| 8 | 59 | Average non-bonded energy per atom |
| 9 | 50 | AAcomposition of CYT of single-spanning proteins |
| 10 | 19 | Hydrophobic parameter pi |

**Supplementary Table 2. Denotations of DSSP Secondary Structures.** DSSP (Dictionary of Secondary Structure of Proteins) is an algorithm developed by Wolfgang Kabsch and Chris Sander to standardize secondary-structure assignments based on atomic coordinates. Using the DSSP package, we analyzed the relationships between secondary structures and LLPS across ten proteomes downloaded from the AlphaFold Protein Structure Database.

| Code | Structure | Description |
|---|---|---|
| H | Alpha Helix (α) | Right-handed coil or spiral helix with 3.6 residues per turn. |
| B | Isolated Beta Bridge | Single pair of hydrogen-bonded beta strands, not part of a full beta sheet. |
| E | Extended Strand | Part of a beta sheet, forming extended strands that are parallel or antiparallel. |
| G | 3/10 Helix | Helix with three residues per turn, a tighter helical turn than an alpha helix. |
| I | Pi Helix (π) | Rare right-handed helix with approximately 4.1 residues per turn. |
| T | Turn | Sharp reversal in the polypeptide chain direction, often at the ends of helices or sheets. |
| S | Bend | Non-hydrogen-bonded bend in the protein chain, providing flexibility and affecting overall shape. |
| P | Polyproline II Helix | Left-handed helix found in proline-rich regions, does not rely on hydrogen bonds. |
| - | Loop or Coil | Irregular structure, not fitting into the regular secondary structure categories, often flexible regions. |

**Supplementary Table 3. Overview of UniProt reference proteomes analyzed in this study.** The table lists the UniProt proteome accession numbers, proteome versions, and total sequence counts for 18 organisms, ranging from *Escherichia coli* to *Homo sapiens*. For each proteome, sequence-level LLPS scores were calculated using Phaseek, followed by phylogenetic and functional enrichment analyses.

| Organism | UniProt Proteome Accession ID | Proteome Version | Gene Count | Alpha Fold | OrgDB |
|---|---|---|---|---|---|
| Escherichia coli (strain K12) | UP000000625 | v4 | 4,401 | ✓ | ✓ |
| Saccharomyces cerevisiae | UP000002311 | v4 | 6,060 | ✓ | ✓ |
| Plasmodium falciparum | UP000001450 | v4 | 5,361 | ✓ | |
| Arabidopsis thaliana | UP000006548 | v4 | 27,448 | ✓ | ✓ |
| Anopheles gambiae | UP000007062 | v4 | 13,017 | | ✓ |
| Drosophila melanogaster (Fruit fly) | UP000000803 | v4 | 13,822 | ✓ | ✓ |
| Caenorhabditis elegans | UP000001940 | v4 | 19,824 | ✓ | ✓ |
| Danio rerio (Zebrafish) (Brachydanio rerio) | UP000000437 | v4 | 25,985 | ✓ | ✓ |
| Xenopus laevis (African clawed frog) | UP000186698 | v4 | 36,175 | | ✓ |
| Gallus gallus (Chicken) | UP000000539 | v4 | 18,370 | | ✓ |
| Rattus norvegicus (Rat) | UP000002494 | v4 | 22,367 | ✓ | ✓ |
| Mus musculus (Mouse) | UP000000589 | v4 | 21,757 | ✓ | ✓ |
| Canis lupus familiaris (Dog) | UP000805418 | v4 | 20,991 | | ✓ |
| Bos taurus (Bovine) | UP000009136 | v4 | 26,942 | | ✓ |
| Sus scrofa (Pig) | UP000008227 | v4 | 22,833 | | ✓ |
| Macaca mulatta (Rhesus macaque) | UP000006718 | v4 | 21,893 | | ✓ |
| Pan troglodytes (Chimpanzee) | UP000002277 | v4 | 23,051 | | ✓ |
| Homo sapiens (Human) | UP000005640 | v4 | 20,650 | ✓ | ✓ |

**Supplementary Table 4. Experimentally tested designed LLPS peptides.** Amino acid (AA) and DNA sequence of the 20 designed peptides experimentally tested in this study, along with their score (with or without GXBoost module applied) and the score of the C-terminal fusion of the peptides to sfGFP.

| Peptide ID | AA sequence | #AA (−M) | DNA sequence | LLPS score (-XGBoost) | LLPS score (+XGBoost) | LLPS score C-terminal GFP fusion (+XGBoost) | worked |
|---|---|---|---|---|---|---|---|
| Phaseek_Gen_1 | MERFGYCKGKASAKKAADSISCVAVS | 25 | ATGGAACGCTTTGGGTATTGTAAGGGGAAAGCCTCCGCTAAGAAGGCCGCCGATAGCATCTCTTGTGTGGCAGTTTCC | 0.83 | 0.29 | 0.70 | no |
| Phaseek_Gen_2 | MYFCSKCRCMTVKATAQARTSMMVIC | 25 | ATGTATTTTTGTAGCAAATGCCGCTGCATGACAGTGAAGGCGACGGCTCAAGCGCGTACCTCCATGATGGTTATTTGC | 0.88 | 0.67 | 0.71 | yes |
| Phaseek_Gen_3 | MFTICPTQGMICTMVVQATTSTKACK | 25 | ATGTTCACGATTTGCCCGACCCAAGGGATGATTTGCACAATGGTTGTTCAGGCAACAACGAGTACGAAAGCCTGTAAG | 0.86 | 0.29 | 0.73 | yes |
| Phaseek_Gen_4 | MKDVKYKKTVMYYYHFDFYFYVKYTFFYKMTWTF | 34 | ATGAAAGACGTAAAGTACAAAAAGACGGTTATGTATTACTACCACTTCGATTTTTACTTCTATGTGAAGTACACCTTCTTTTACAAGATGACATGGACCTTT | 0.26 | 0.26 | 0.65 | yes |
| Phaseek_Gen_5 | MLMIRESFVQQPFIIYVFAYYAKFYMYVVHFVHHFF | 36 | ATGTTGATGATTCGCGAGAGCTTTGTACAACAGCCGTTTATTATTTACGTCTTCGCCTATTACGCTAAGTTTTACATGTACGTAGTCCATTTCGTTCACCATTTCTTT | 0.30 | 0.30 | 0.72 | yes |
| Phaseek_Gen_6 | MPVGPGEGPQEPPQEEGPGGEEEEGEGEPGEEEEDEPEYIHFKFYL | 45 | ATGCCTGTCGGACCTGGCGAGGGGCCCCAGGAGCCTCCTCAGGAAGAAGGCCCAGGTGGTGAGGAAGAAGAAGGGGAGGGTGAACCCGGAGAAGAGGAAGAAGATGAACCTGAATATATCCATTTCAAATTTTATTTG | 0.96 | 0.64 | 0.76 | yes |
| Phaseek_Gen_7 | MTPCKCSQEGQPGGCQPQGEGQPGPPEPEEGGGEPGESMPDYCRMGFF | 47 | ATGACCCCCTGTAAGTGCAGCCAGGAAGGCCAGCCAGGGGGATGCCAACCGCAAGGCGAAGGGCAACCGGGGCCGCCAGAGCCCGAAGAAGGTGGGGGTGAGCCAGGTGAGTCTATGCCTGACTACTGCCGCATGGGTTTTTTT | 0.94 | 0.94 | 0.76 | yes |
| Phaseek_Gen_8 | MQGMEPEEEGEPPESPEGGEEPPEPEEEPPQPGGPQKPKEMYEVYGVHFY | 49 | ATGCAAGGCATGGAACCAGAGGAAGAGGGTGAGCCCCCTGAAAGCCCAGAAGGTGGAGAGGAACCACCAGAGCCCGAGGAGGAGCCCCCGCAGCCAGGCGGGCCTCAAAAACCCAAAGAAATGTACGAGGTGTATGGGGTCCACTTTTAT | 0.96 | 0.95 | 0.78 | yes |
| Phaseek_Gen_9 | MQFPGGQGPEEEPGPPGEGPQGPGGEGGPGGGGGPGTTESYFEFVFFKMF | 49 | ATGCAGTTTCCTGGTGGCCAGGGGCCAGAGGAAGAACCAGGTCCGCCCGGTGAGGGACCTCAGGGCCCTGGTGGGGAAGGTGGACCCGGGGGCGGTGGTGGTCCTGGTACAACTGAGTCATATTTTGAGTTCGTTTTTTTTAAGATGTTT | 0.95 | 0.98 | 0.78 | yes |
| Phaseek_Gen_10 | MELGMGRLVRKAAVKAASKWEAVGVFFKFFVFAVQGAQKVKAGGDKAKKM | 50 | ATGGAATTGGGTATGGGCCGTTTAGTACGTAAAGCTGCAGTGAAAGCAGCGTCAAAATGGGAAGCTGTTGGGGTGTTCTTTAAATTCTTTGTGTTCGCGGTTCAGGGAGCCCAAAAGGTGAAGGCTGGGGGCGATAAAGCCAAGAAAATG | 0.30 | 0.30 | 0.70 | no |
| Phaseek_Gen_11 | MEGQEEQEGQEEPGSSQPPPEPGEPEGGEPEGEKEGKTYPYDMFDHFGMFRL | 51 | ATGGAGGGCCAAGAGGAGCAAGAGGGTCAAGAGGAACCAGGTTCTTCCCAGCCCCCACCCGAGCCCGGCGAACCTGAAGGAGGGGAGCCGGAGGGGGAGAAGGAAGGAAAAACATACCCATATGACATGTTCGACCATTTTGGCATGTTTCGTCTT | 0.94 | 0.94 | 0.78 | yes |
| Phaseek_Gen_12 | MKQILPEEEQQEQPGQQPGPGEPPGPEGGGGEPEEQKGEEDGLEFPVLAYPL | 51 | ATGAAGCAGATCCTTCCTGAAGAGGAACAACAAGAGCAACCTGGACAACAGCCGGGACCAGGTGAACCGCCTGGTCCTGAGGGCGGAGGCGGAGAGCCCGAGGAACAGAAGGGAGAAGAAGATGGATTAGAATTTCCGGTCTTGGCTTACCCACTT | 0.95 | 0.94 | 0.78 | no |

**Supplementary Table 4 (continued)**

| Peptide ID | AA sequence | #AA (−M) | DNA sequence | LLPS score (-XGBoost) | LLPS score (+XGBoost) | LLPS score C-terminal GFP fusion (+XGBoost) | worked |
|---|---|---|---|---|---|---|---|
| Phaseek _Gen_13 | MCQPEPPEE GPEPGPPPG EGQCPGPPG EEPEPPPGE EQEPMGEMP HMDHDPHHD PAFAYHK | 60 | ATGTGCCAGCCAGAACCTCCGGAGGAGGG CCCCGAACCAGGGCCGCCTCCTGGTGAAG GTCAATGTCCTGGGCCACCTGGTGAGGAAC CCGAGCCTCCACCCGGTGAAGAACAGGAAC CAATGGGTGAAATGCCCCATATGGACCATG ATCCGCATCATGACCCGGCGTTCGCATACC ATAAA | 0.94 | 0.93 | 0.78 | no |
| Phaseek _Gen_14 | MPYWVYYLH VKAHETFCS KVVVRESWF QLVVWTFGS GCVKVAGAM VKKHAQKQA YGCFEGESC | 63 | ATGCCCTACTGGGTCTACTACTTACACGTAA AGGCTCACGAGACGTTCTGCTCCAAGGTGG TGGTCCGCGAAAGCTGGTTTCAACTGGTTG TGTGGACATTCGGCTCCGGTTGTGTCAAGG TTGCAGGCGCGATGGTGAAAAAACACGCCC AGAAACAAGCCTATGGTTGTTTTGAAGGAG AATCATGC | 0.24 | 0.24 | 0.71 | yes |
| Phaseek _Gen_15 | MFGYQQGE GQQPPGGE GGGGGGEG PEPEPPGGE EEGGQKEDP ELFFFDWFD VHKRALAHM PYYDA | 64 | ATGTTCGGATATCAACAAGGTGAAGGCCAG CAGCCACCGGGAGGAGAAGGGGGTGGCG GTGGAGGTGAGGGTCCCGAGCCAGAACCT CCAGGCGGCGAGGAAGAAGGTGGACAAAA GGAAGATCCGGAGTTGTTCTTCTTCGATTG GTTCGACGTCCACAAGCGCGCACTTGCGCA CATGCCGTATTACGATGCG | 0.94 | 0.93 | 0.78 | no |
| Phaseek _Gen_16 | MRMGEGGE GEPEQQEQQ GGEEEPGPP GGPPPGPGG GGRKPKMPE GGFHLDDYF TFLGEEEDD GAPGE | 66 | ATGCGCATGGGCGAGGGAGGCGAAGGAGA ACCAGAGCAGCAGGAGCAACAGGGCGGAG AAGAGGAGCCAGGACCACCTGGAGGTCCA CCTCCTGGACCTGGCGGAGGTGGCCGCAA GCCTAAGATGCCAGAAGGTGGTTTTCACCT GGATGATTACTTCACCTTTTTGGGAGAAGAA GAGGACGACGGGGCGCCGGGGGAA | 0.95 | 0.94 | 0.79 | no |
| Phaseek _Gen_17 | MEYPGGGGE EQPPPEPGG PEEPEEEGP GGEPEEMEP EQEPKFRVR FMSWYKVHP RAEEKGGDE GDHDD | 67 | ATGGAATATCCGGGTGGTGGTGGTGAAGAA CAGCCGCCCCCCGAACCTGGCGGTCCGGA AGAGCCGGAGGAAGAGGGCCCTGGGGGAG AACCTGAAGAAATGGAACCAGAGCAGGAGC CTAAGTTCCGCGTTCGTTTTATGTCCTGGTA TAAGGTACACCCTCGCGCCGAAGAAAAAGG CGGGGACGAGGGAGACCACGATGAC | 0.95 | 0.95 | 0.78 | yes |
| Phaseek _Gen_18 | MQHTEQQQ GGEGGESGE GEGQEPGPP EGGPGPPPE GKSKGVYPV FLDDMDPIFY DARHMVRLT LVMDLVLKFD VTTDL | 77 | ATGCAACACACTGAGCAGCAACAAGGAGGT GAGGGTGGAGAAAGCGGAGAGGGAGAGGG GCAAGAACCTGGGCCACCTGAAGGGGGAC CAGGCCCGCCCCCGGAGGGAAAAAGCAAG GGCGTTTACCCCGTGTTCTTGGATGATATG GATCCGATTTTTTACGATGCCCGCCATATGG TTCGTCTGACCCTGGTGATGGATTTAGTTTT AAAATTTGACGTAACTACCGATTTA | 0.93 | 0.40 | 0.77 | yes |
| Phaseek _Gen_19 | MLDEQPLISV CAAVATEQIS QTATVKKVQ KVKVHVACD AKVAGGYHY GKKTYSVYK SEGQHQCSG SSKGMQYVK EFSM | 78 | ATGTTGGATGAGCAACCTTTAATTTCAGTAT GTGCAGCAGTAGCCACCGAGCAGATCTCTC AGACCGCCACGGTTAAGAAAGTGCAAAAGG TTAAAGTTCACGTTGCATGCGATGCCAAGGT AGCAGGAGGATACCACTACGGAAAGAAAAC ATATTCTGTTTACAAGTCGGAGGGTCAGCAT CAGTGTAGTGGTAGTAGCAAGGGAATGCAG TACGTAAAAGAGTTCTCAATG | 0.28 | 0.28 | 0.74 | yes |
| Phaseek _Gen_20 | MYGTQGQEE EGGPEQEQG GPGPPPPGG GGEGPGPEQ EGYKEGLQL LKHFLEWFTF LEGKPDDDP DGFGLPKEL DEAFDLEA | 80 | ATGTACGGAACTCAGGGACAAGAAGAAGAG GGTGGGCCCGAACAGGAGCAAGGGGGGCC TGGACCGCCTCCGCCAGGCGGTGGTGGCG AAGGACCGGGGCCGGAGCAAGAAGGTTAC AAAGAAGGATTACAATTGCTGAAGCACTTCC TTGAGTGGTTCACGTTCTTAGAGGGAAAAC CGGATGATGATCCCGACGGTTTTGGGTTAC CAAAAGAACTGGACGAAGCTTTCGATTTAGA AGCA | 0.92 | 0.91 | 0.78 | yes |

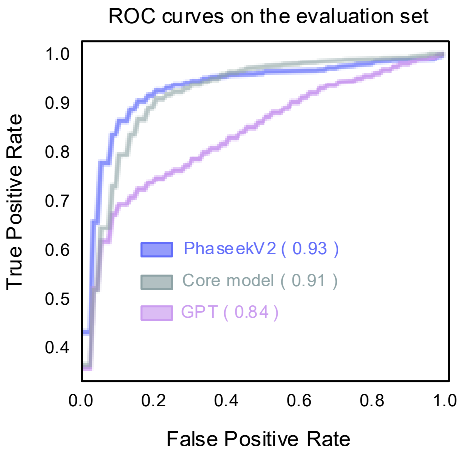


| | Accuracy | Specificity | Sensitivity | F1 Score |
|---|---|---|---|---|
| Phaseek | 0.88 | 0.86 | 0.90 | 0.87 |
| Core model | 0.75 | 0.51 | 0.96 | 0.66 |
| GPT | 0.74 | 0.48 | 0.96 | 0.63 |

**Supplementary Fig. 1.** Evaluation of FEGS-matrix incorporation and region-boosting on GPT model performance. We compared three models on the evaluation dataset: (1) a baseline GPT architecture (XM parameters) trained on the original training data, (2) the same GPT model enhanced with FEGS matrices (GPT + FEGS, "core model"), and (3) the final Phaseek architecture(PhaseekV2). Receiver operating characteristic (ROC) curves and their corresponding area under the curve (AUC) scores are shown for each model. A summary table reports the models' accuracy, specificity, sensitivity, and F1 score on the evaluation data.

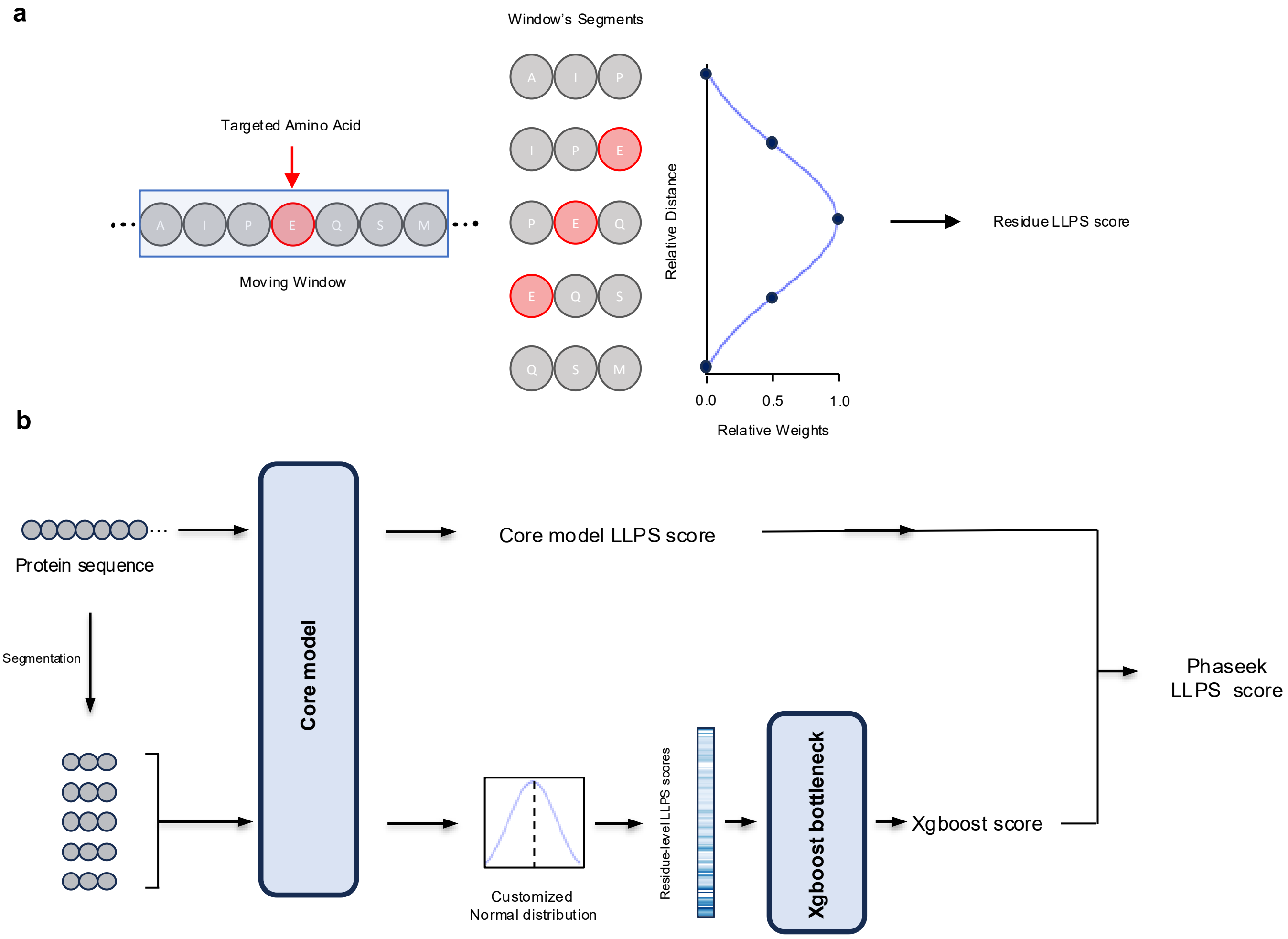


**Supplementary Fig. 2. Phaseek scoring and slicing mechanism for computing final and residue-level LLPS scores. (a)** The input sequence is segmented using a sliding window of fixed length. Each segment receives an LLPS score from the model and is weighted based on the position of the central residue using a Gaussian distribution (N(0,1)). Residue-level LLPS scores are calculated by aggregating the weighted segment scores (see *Methods*). **(b)** The residue-level LLPS scores are used as input features for the XGBoost bottleneck module, which produces an XGBoost score. The final Phaseek LLPS score is obtained by taking the median of the XGBoost and core model scores.

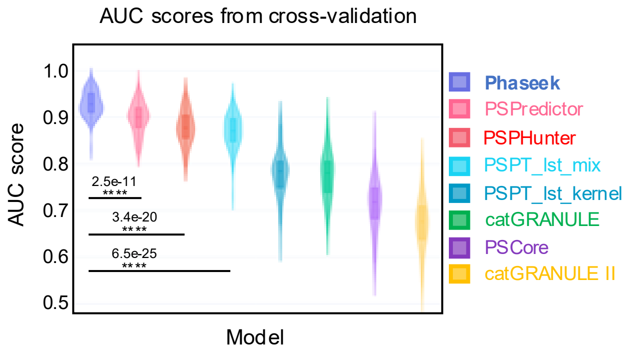


**Supplementary Fig. 3. Models benchmark on the validation dataset.** Violin plots illustrating ROC area under the curve (AUC) scores for Phaseek and other models, evaluated using Monte Carlo Cross-Validation (MCCV). Batches containing 40 phase-separating and 40 non-phase-separating sequences from the evaluation set were selected and tested over 100 iterations. The statistical significance is indicated by the one-sided Mann–Whitney U p-values (****p < 0.0001). Violin plots represent data distributions, and boxes indicate the interquartile range (Q1–Q3) with the median marked by the center line. Models are color-coded in the figure: Phaseek (periwinkle blue), PSPhunter (deep salmon), catGRANULE I (Caribbean green), PScore (medium purple), catGRANULE II (saffron yellow), PSPredictor (coral pink), PSTP-LST-mix (bright sky blue), and PSTP-LST-kernel (deep cerulean).

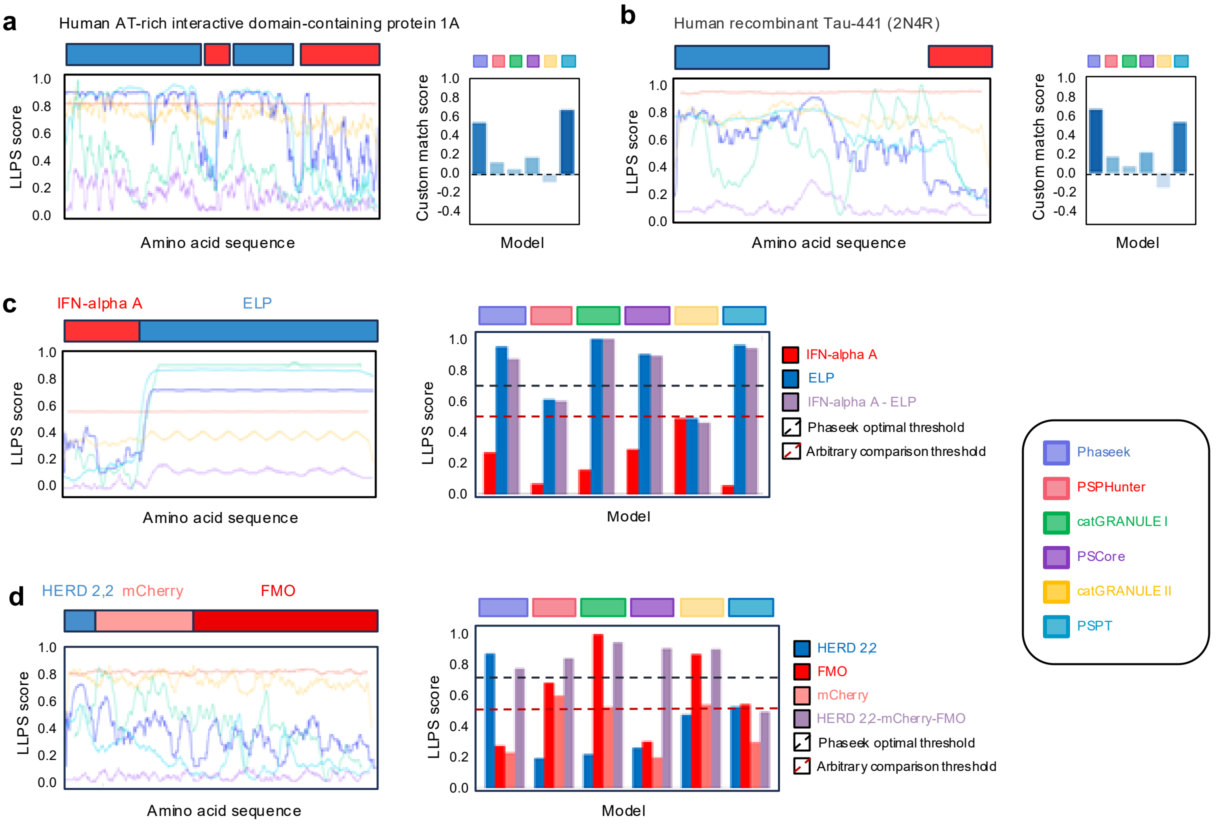


**Supplementary Figure 4. Phaseek identifies key LLPS regions and predicts the impact of peptide fusion on phase separation.** (a, b) Residue-level LLPS predictions by different models and their comparison using a custom match score (bar charts, see Methods) on two human proteins not seen during Phaseek training, with experimentally annotated phase-separating (blue) and non-phase-separating (red) regions. The proteins shown are (a) ARID1A (AT-rich interactive domain-containing protein 1A) and (b) recombinant Tau. (c, d) Whole-sequence LLPS score variations and residue-level profiles following sequence fusion with known or designed LLPS-prone components. Shown are: (c) the fusion of an elastin-like polypeptide (ELP) with Interferon-alpha A (IFN-αA), and (d) the fusion of the synthetic phase-separating peptide HERD 2,2 with mCherry and FMO (flavin-containing monooxygenase). Bar charts indicate the overall predicted LLPS propensity of the fusion constructs. To facilitate comparison across models, we applied an arbitrary threshold of 0.5, in addition to Phaseek's optimal threshold of 0.7. Both thresholds are indicated by dashed lines in the figure.

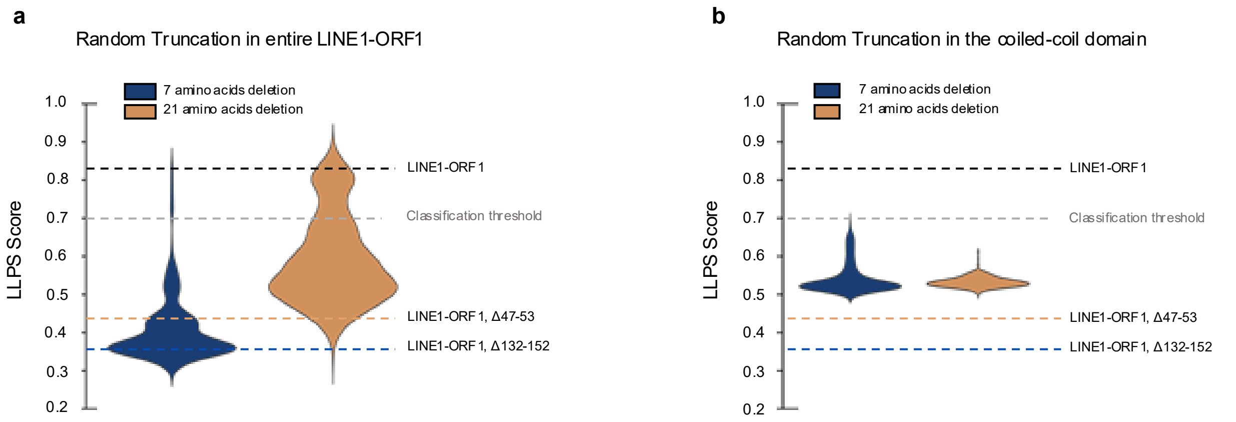


**Supplementary Figure 5. LLPS scores for random truncations in the LINE1-ORF1 sequence.** Violin plots show the predicted Phaseek LLPS scores for randomly truncated variants of the LINE1-ORF1 sequence, either across the full protein (a) or restricted to the coiled-coil region (b). Truncations of 21 amino acids are shown in light copper, and truncations of 7 amino acids are shown in blue. Horizontal lines indicate the LLPS scores of the wild-type LINE1-ORF1, as well as two designed truncation mutants: LINE1-ORF1-Δ47–53 and LINE1-ORF1-Δ132–152, for comparison against the distribution of random truncations. The model's optimal classification threshold is indicated at 0.7.

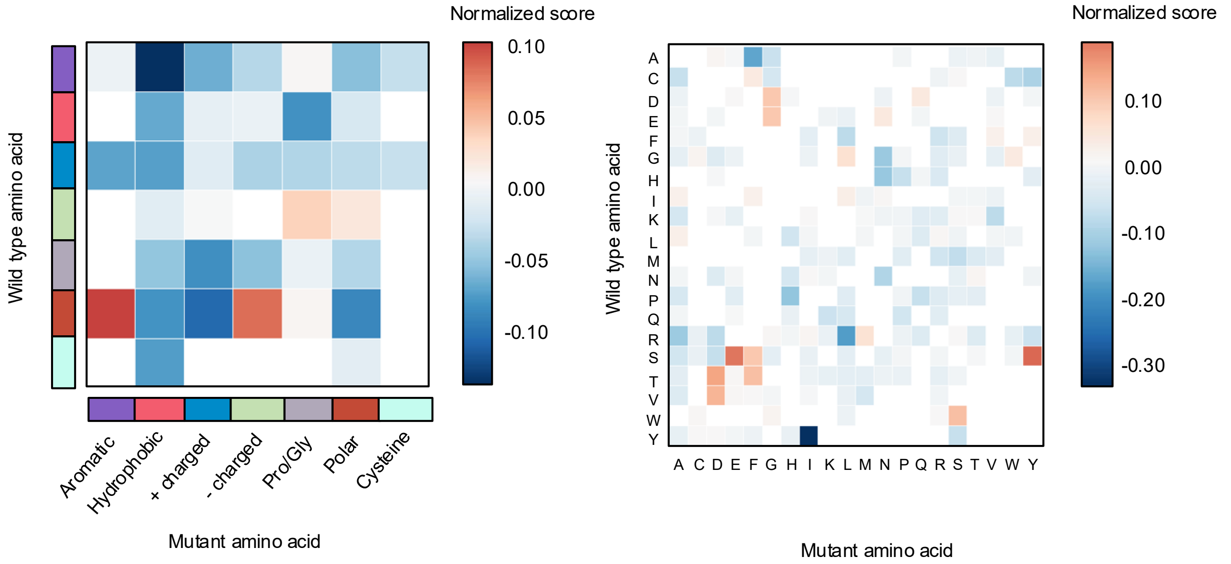


**Supplementary Figure 6.** Heatmaps presenting the normalized scores of mutation impacts, derived from collected impactful mutation data (2529 impactful mutations) to quantify the effect of amino acid substitutions by calculating a normalized score for each mutation type, based on the score difference (Δ(aa_Score)) predicted by Phaseek. The normalized score is computed as Normalized Score = $\frac{Avg_{neg}\cdot n_{neg} + Avg_{pos}\cdot n_{pos}}{n_{neg} + n_{pos}}$ where $n_{pos}$ and $n_{neg}$ are the counts of positive and negative score differences, and $Avg_{pos}$ and $Avg_{neg}$ are their respective mean values. The left heatmap groups wild-type and mutant amino acids into categories (Aromatic, Hydrophobic Negatively Charged, Positively Charged, Gly/Pro, Polar, and Cysteine), where Normalized Scores are computed to reflect the overall impact across groups. The right heatmap illustrates this analysis on individual wild-type and mutant amino acids. This analysis provides a balanced representation of mutation effects, accounting for both the frequency of occurrences and the magnitude of score changes.

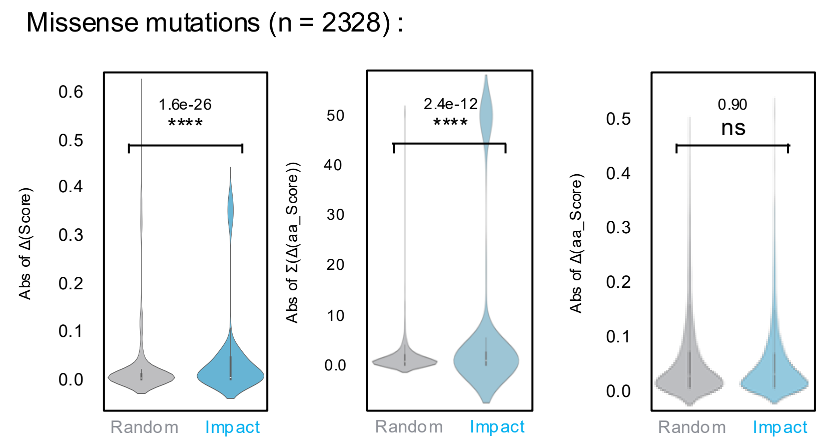


**Supplementary Figure 7. Phaseek accurately distinguishes missense mutations from random ones.** Violin plots showing the distribution of three custom metrics used to assess the impact of point mutations on LLPS: Δ(Score), Δ(aa_Score), and Σ(Δ(aa_Score)). These metrics reflect sequence-level, single-residue-level, and cumulative regional effects, respectively. Blue violins represent missense mutations (n=2328), while gray violins correspond to 2740 randomly generated mutations. P-values were calculated using one-sided Mann–Whitney U tests, with statistical significance indicated by asterisks (****p < 0.0001, ***p < 0.001, **p < 0.01, *p < 0.05, ns = not significant).

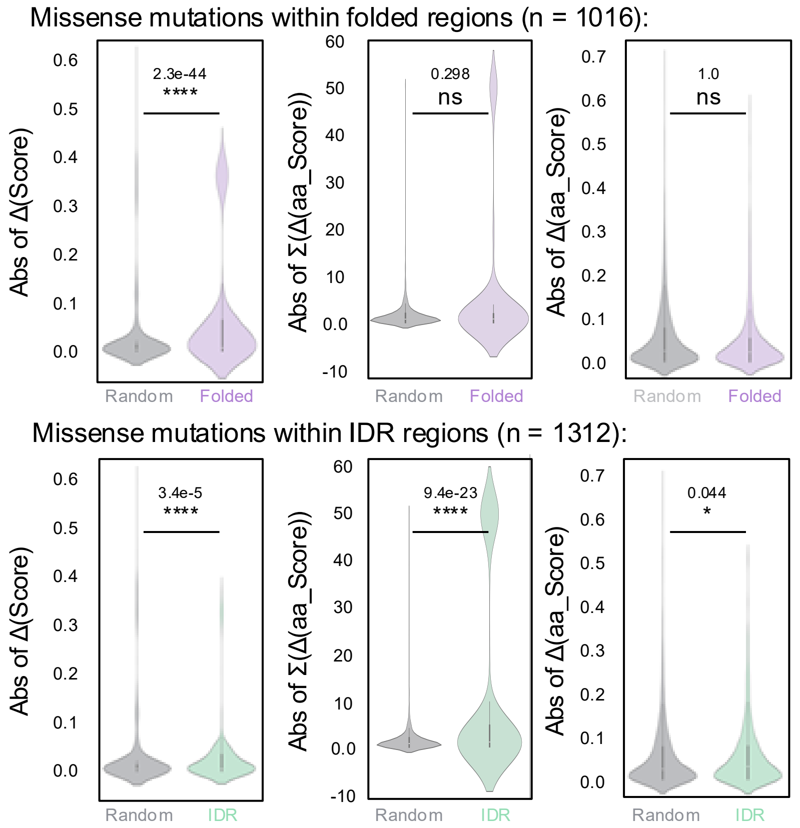


**Supplementary Figure 8. Phaseek identifies impactful (missense and labelled) mutations across both folded and IDR regions.** Violin plots show the distribution of three custom scores: Δ(Score), Δ(aa Score), and Σ(Δ(aa Score)) for all impactful mutations (green and purple) and randomly generated mutations (gray, n = 2740). Two analyses are presented: (a) all impactful mutations within folded regions (n = 1016), (b) all impactful mutations within IDR regions (n = 1312). Residues were categorized as folded or IDR based on the average AIUPred score across a 5-residue window centered on the mutation site, positions with an average score ≤ 0.5 were considered folded, and those with a score > 0.5 were considered IDR. With the exception of the Σ(Δ(aa Score)) metric, impactful mutations showed higher scores than random mutations, highlighting Phaseek's ability to detect impactful mutations within both folded and disordered regions. P-values were calculated using one-sided Mann–Whitney U tests, with statistical significance indicated by asterisks (****p < 0.0001, ***p < 0.001, **p < 0.01, *p < 0.05, ns = not significant).

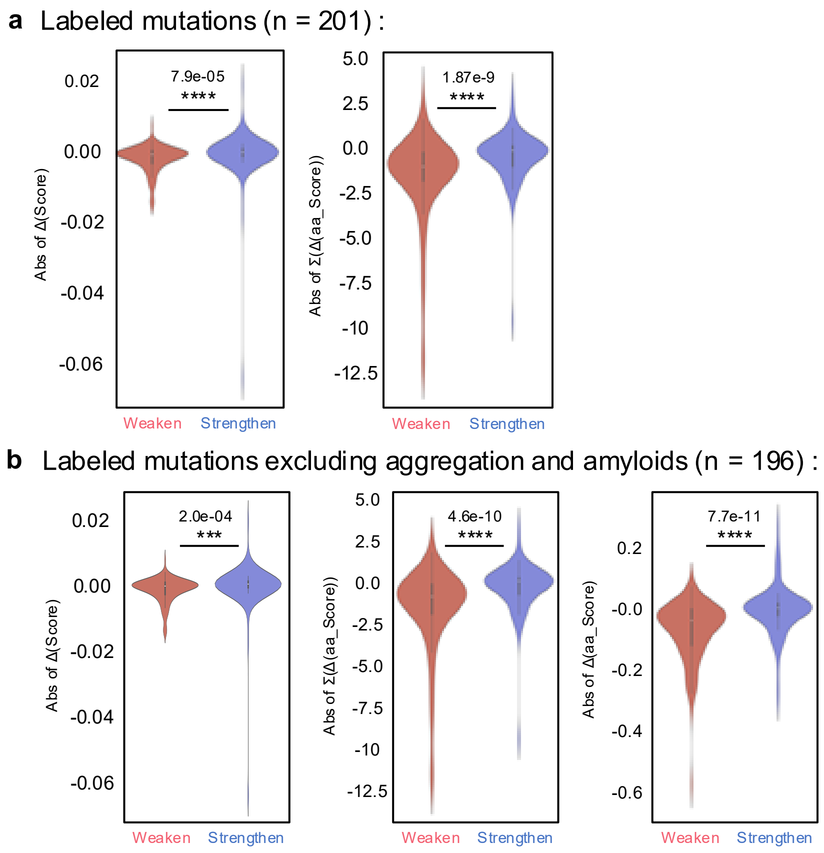


**Supplementary Figure 9. Phaseek effectively discriminates between LLPS-strengthening and LLPS-weakening mutations**. Violin plots show the distribution of three custom scores: Δ(Score), Δ(aa Score), and Σ(Δ(aa Score)) for LLPS-strengthening (blue) and LLPS-weakening (red) mutations. Two analyses are shown: (a) all labeled mutations (n = 93 strengthening, 114 weakening), (b) excluding mutations associated with aggregation or amyloids (n = 87 strengthening, 114 weakening). In both cases, all three scores are significantly higher for LLPS-strengthening mutations, demonstrating Phaseek's ability to distinguish mutations that enhance versus disrupt phase separation. P-values are from one-sided Mann–Whitney U tests, with statistical significance indicated by asterisks (****p < 0.0001, ***p < 0.001, **p < 0.01, *p < 0.05, ns = not significant).

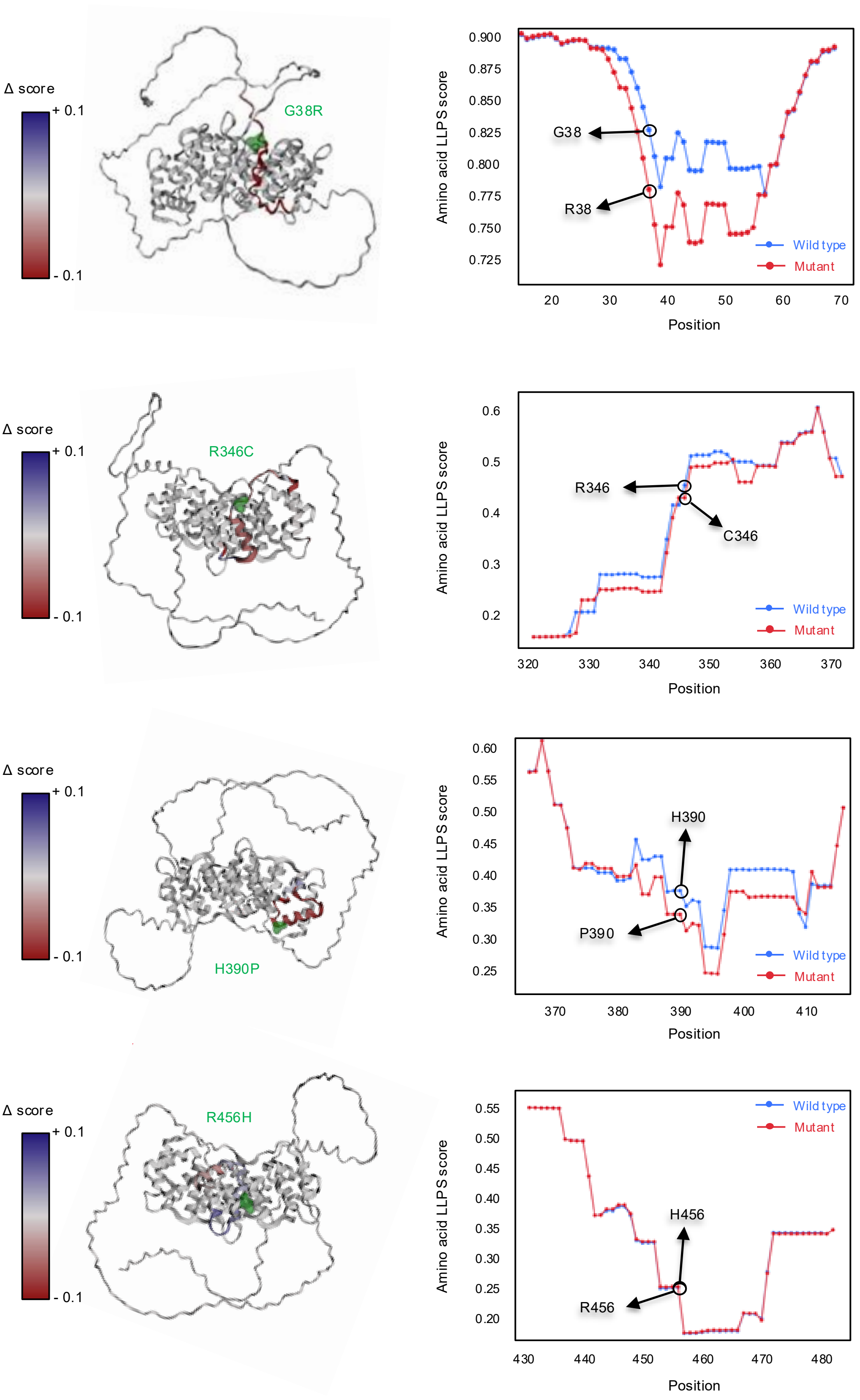


**Supplementary Figure 10. Residue-level LLPS score shifts induced by impactful point mutations in Human Annexin A11.** Left panels show the 3D structure of Annexin A11 predicted with Alphafold2 with the wild-type (red) structure and mutant residue is highlighted with green surface. Right panels plot the amino acid–level LLPS scores for both wild-type (blue) and mutant (red) sequences around the mutation site. Arrows mark the exact mutation positions. The structural coloring reflects the Δ(Score), demonstrating that the model's performance in identifying both LLPS-promoting and -disrupting mutations.

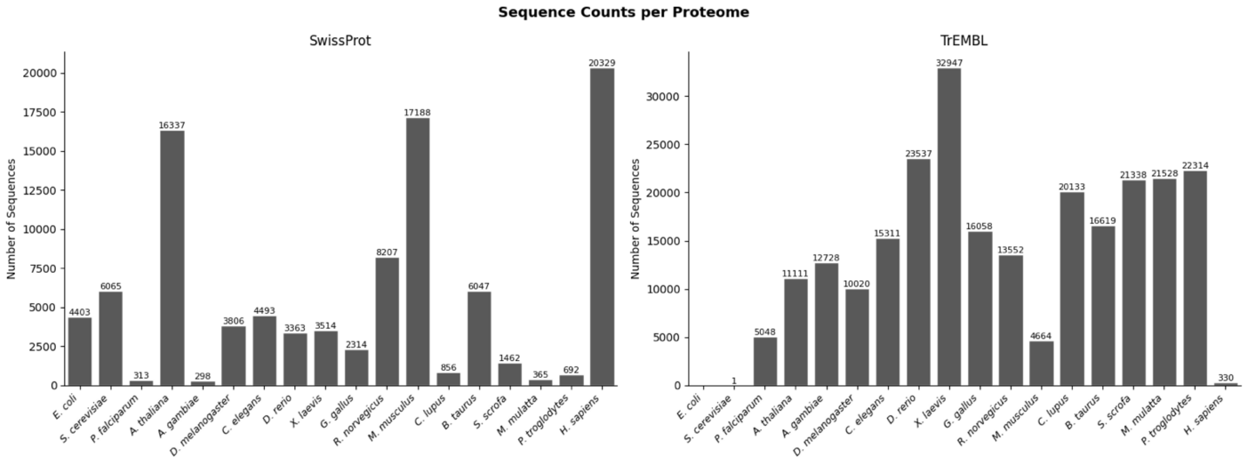


**Supplementary Figure 11. Number of coding sequences scored by Phaseek for all 18 organisms.** Left panel corresponds to manually curated sequences from SwissProt. Right panel corresponds to computationally predicted sequences from TrEMBL.

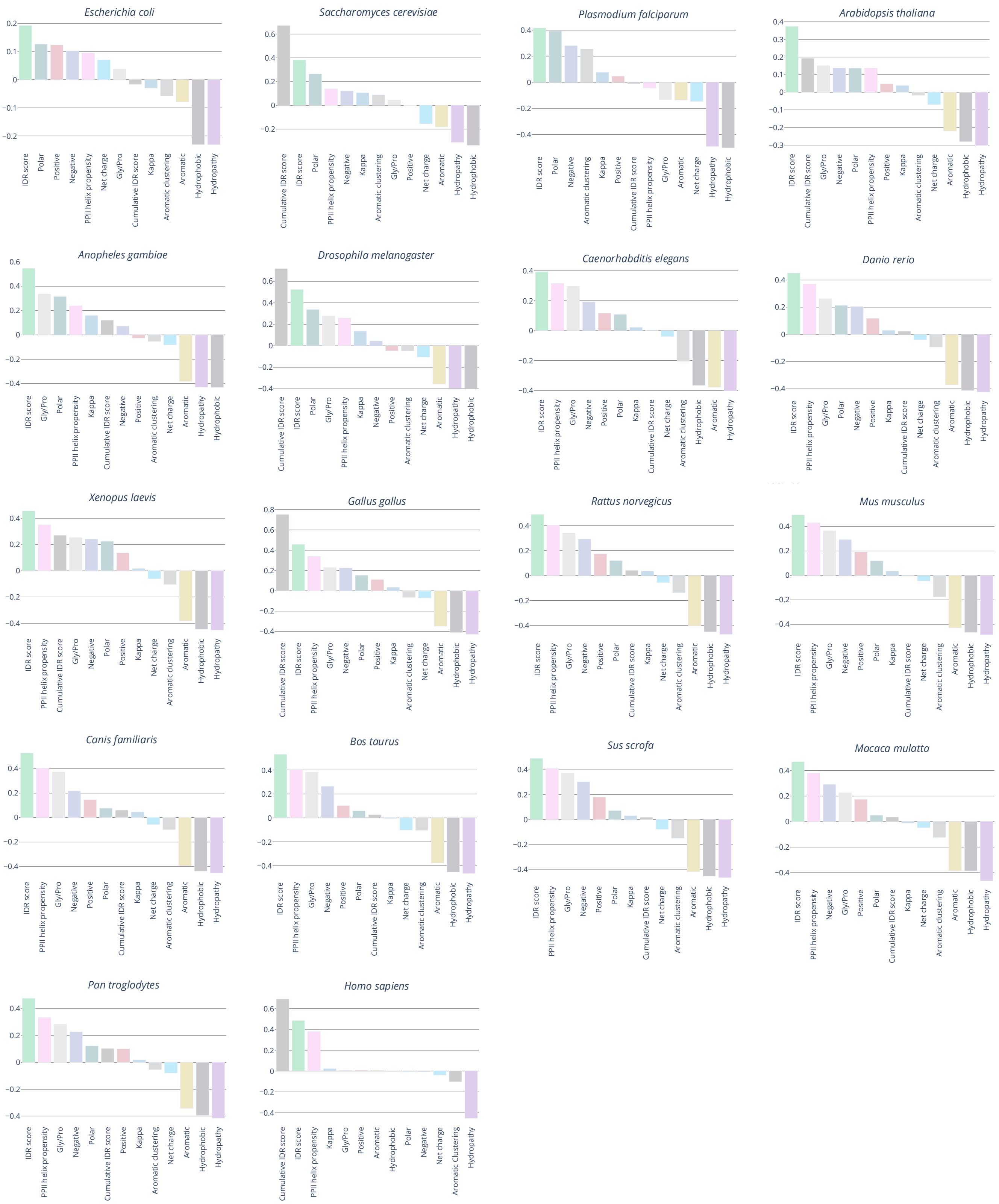


**Supplementary Figure 12. species-specific and combined analysis of predicted LLPS propensity and physicochemical features across 18 proteomes.** Spearman correlation coefficients between physicochemical properties and Phaseek-predicted LLPS scores are shown for each of the 18 individual proteomes, as well as for the combined dataset.

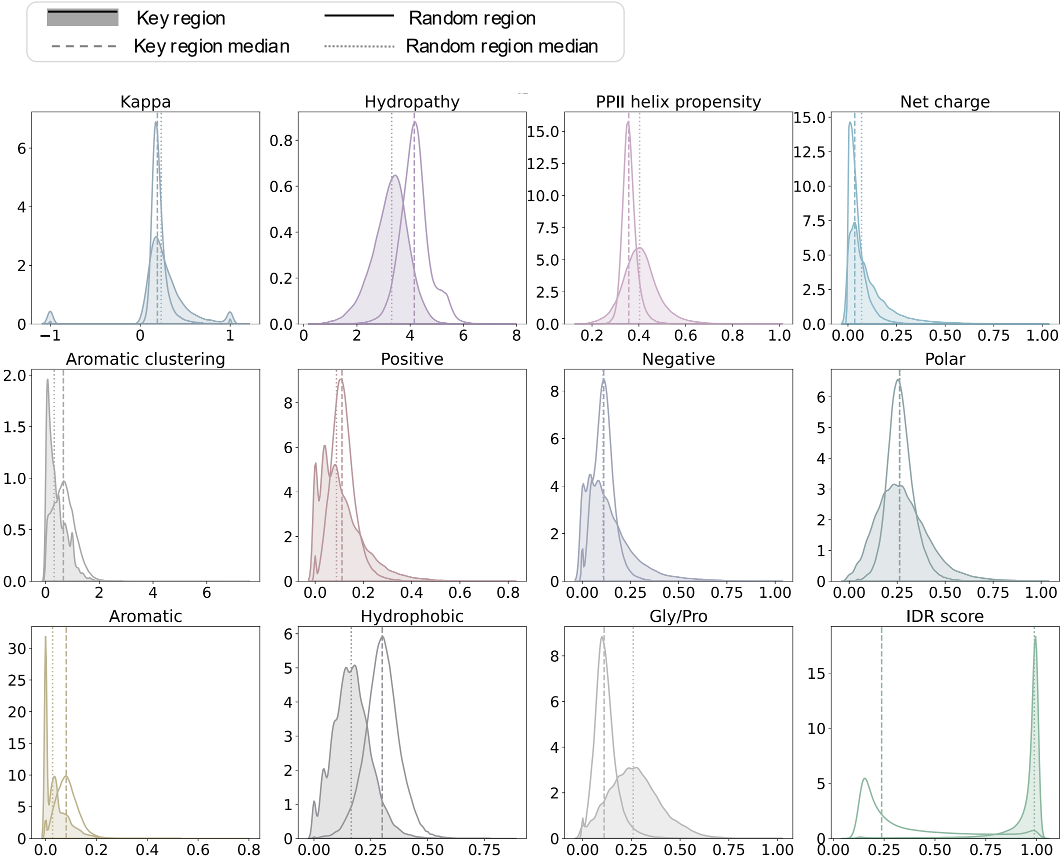


**Supplementary Figure 13. Physicochemical feature analysis in key and random regions of all 18 proteomes.** Density distributions of physicochemical features for random and key regions across all 18 proteomes analyzed in this study. Median for key and random regions are represented by dashed and dotted lines respectively.

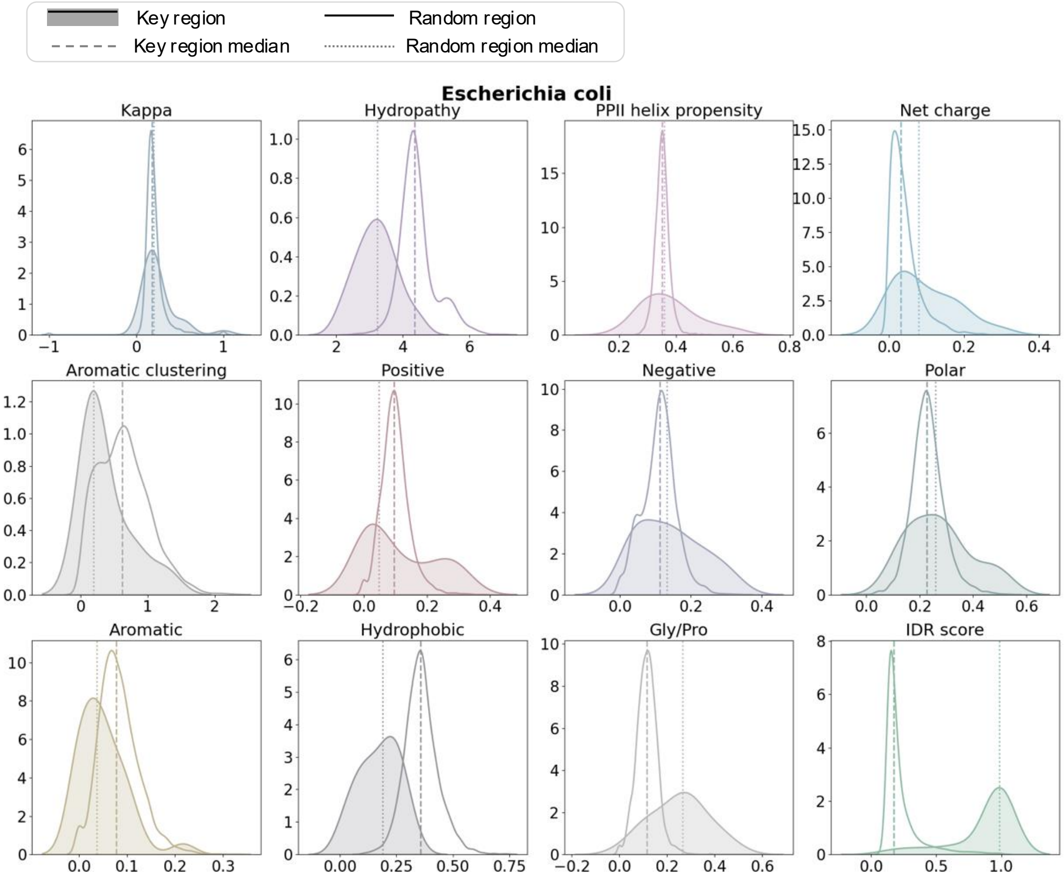


**Supplementary Figure 14. Physicochemical feature analysis in key and random regions of *Escherichia coli*.** Density distributions of physicochemical features for random and key regions across all 18 proteomes analyzed in this study. Median for key and random regions are represented by dashed and dotted lines respectively.

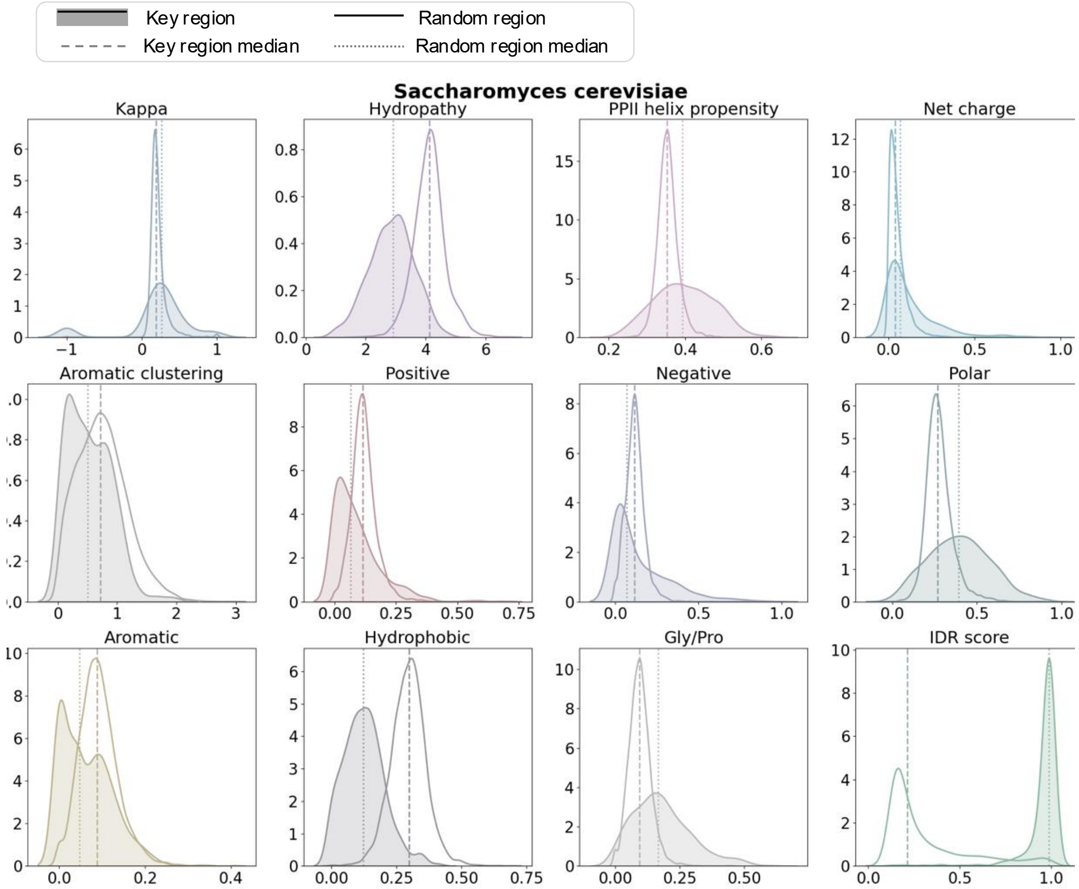


**Supplementary Figure 15. Physicochemical feature analysis in key and random regions of *Saccharomyces cerevisiae*.** Density distributions of physicochemical features for random and key regions across all 18 proteomes analyzed in this study. Median for key and random regions are represented by dashed and dotted lines respectively.

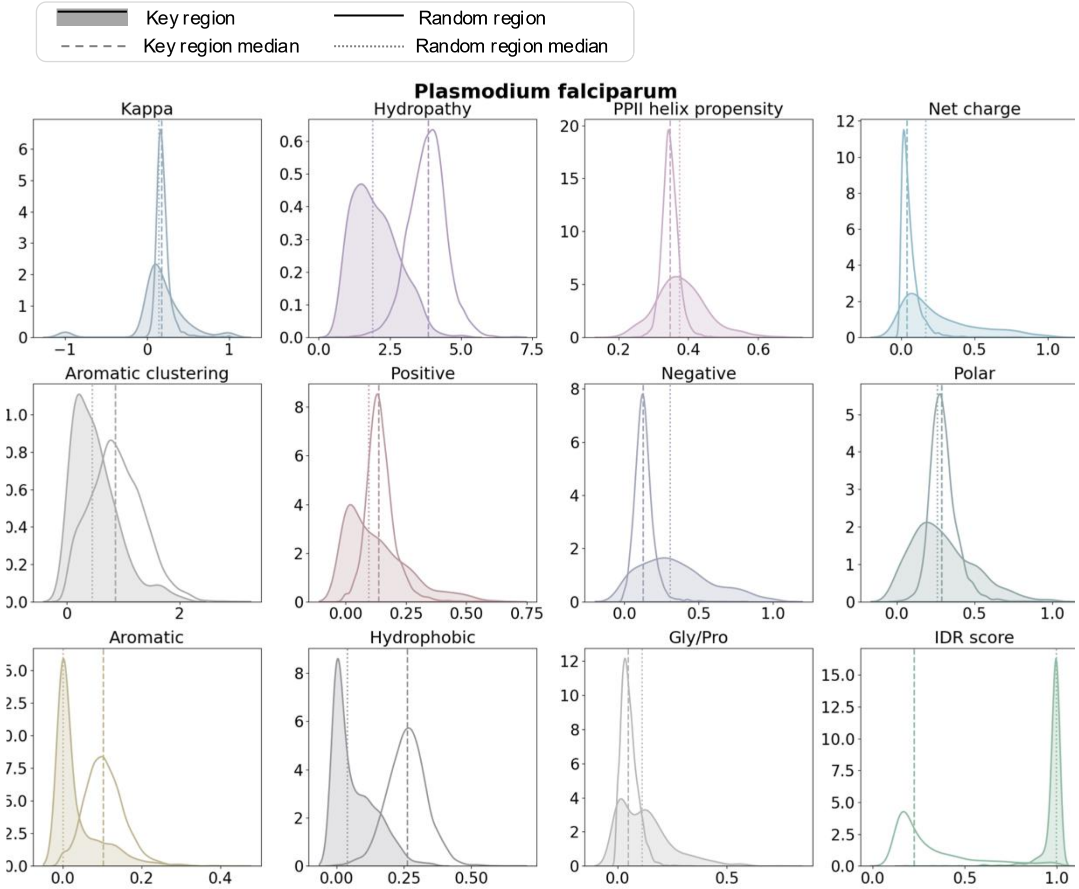


**Supplementary Figure 16. Physicochemical feature analysis in key and random regions of *Plasmodium falciparum*.** Density distributions of physicochemical features for random and key regions across all 18 proteomes analyzed in this study. Median for key and random regions are represented by dashed and dotted lines respectively.

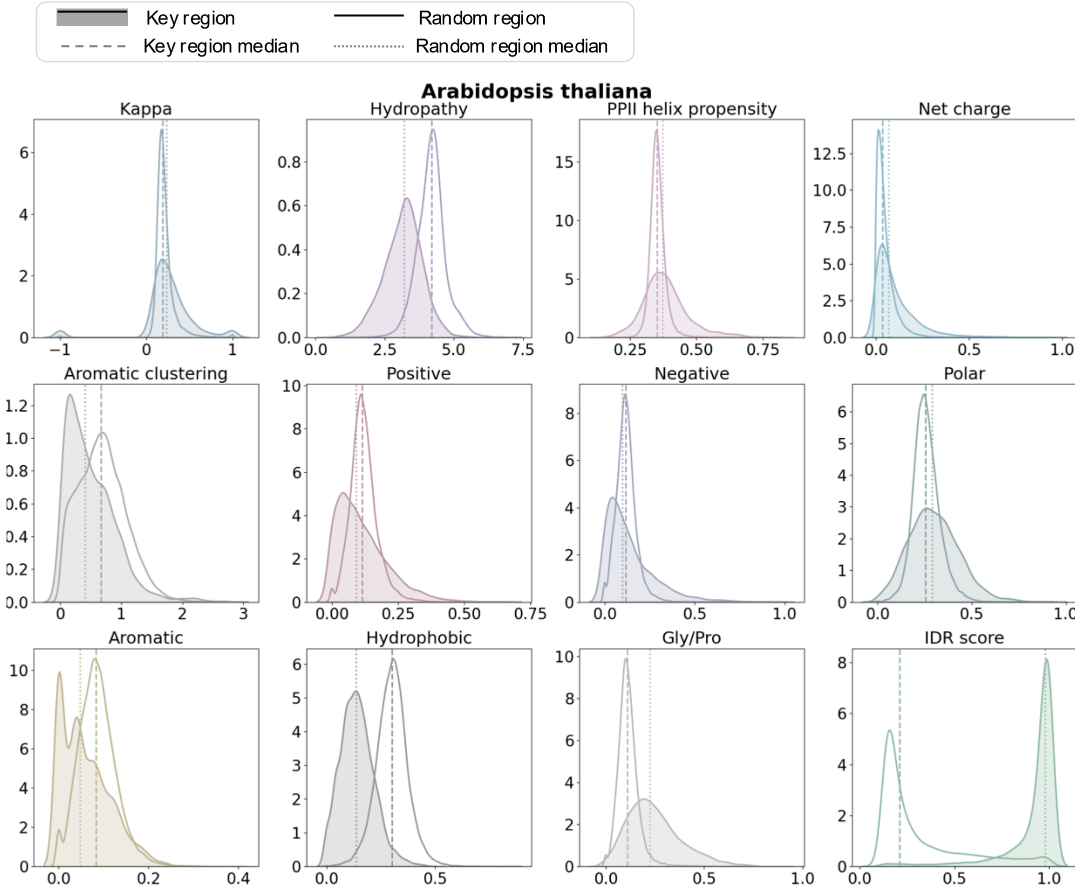


**Supplementary Figure 17. Physicochemical feature analysis in key and random regions of *Arabidopsis thaliana*.** Density distributions of physicochemical features for random and key regions across all 18 proteomes analyzed in this study. Median for key and random regions are represented by dashed and dotted lines respectively.

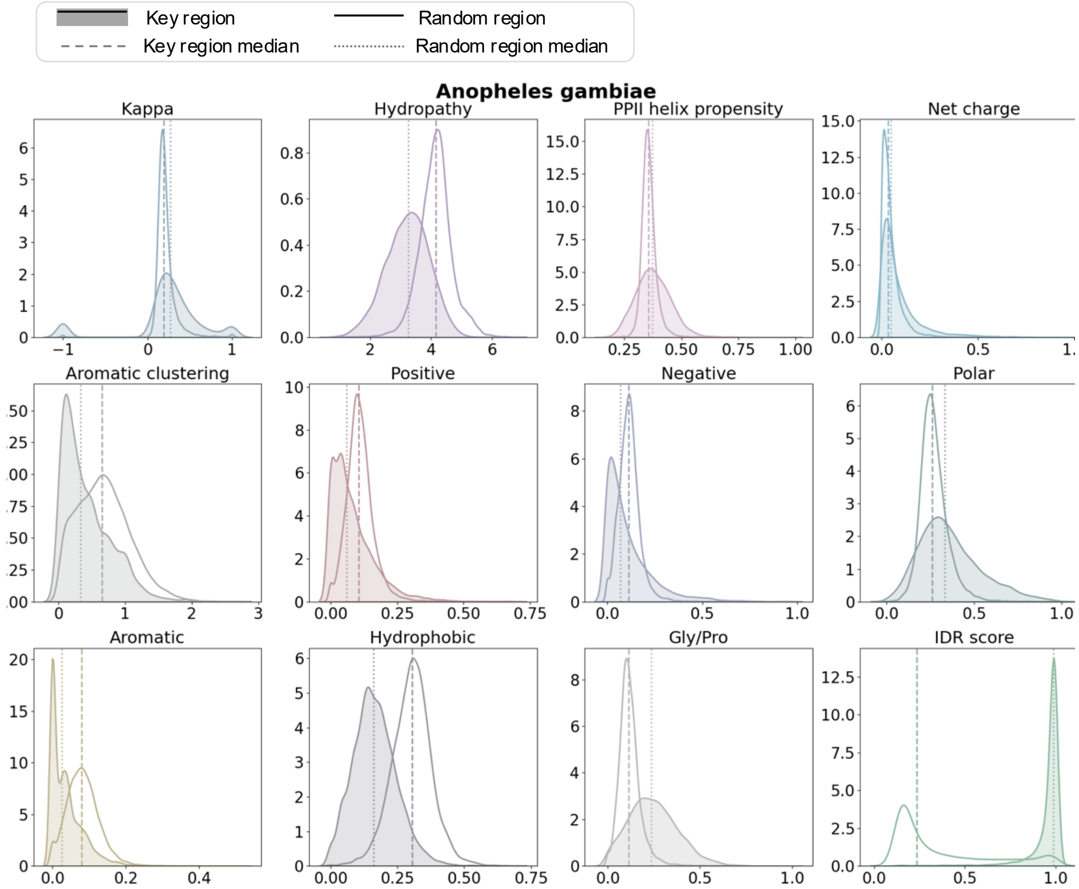


**Supplementary Figure 18. Physicochemical feature analysis in key and random regions of *Anopheles gambiae*.** Density distributions of physicochemical features for random and key regions across all 18 proteomes analyzed in this study. Median for key and random regions are represented by dashed and dotted lines respectively.

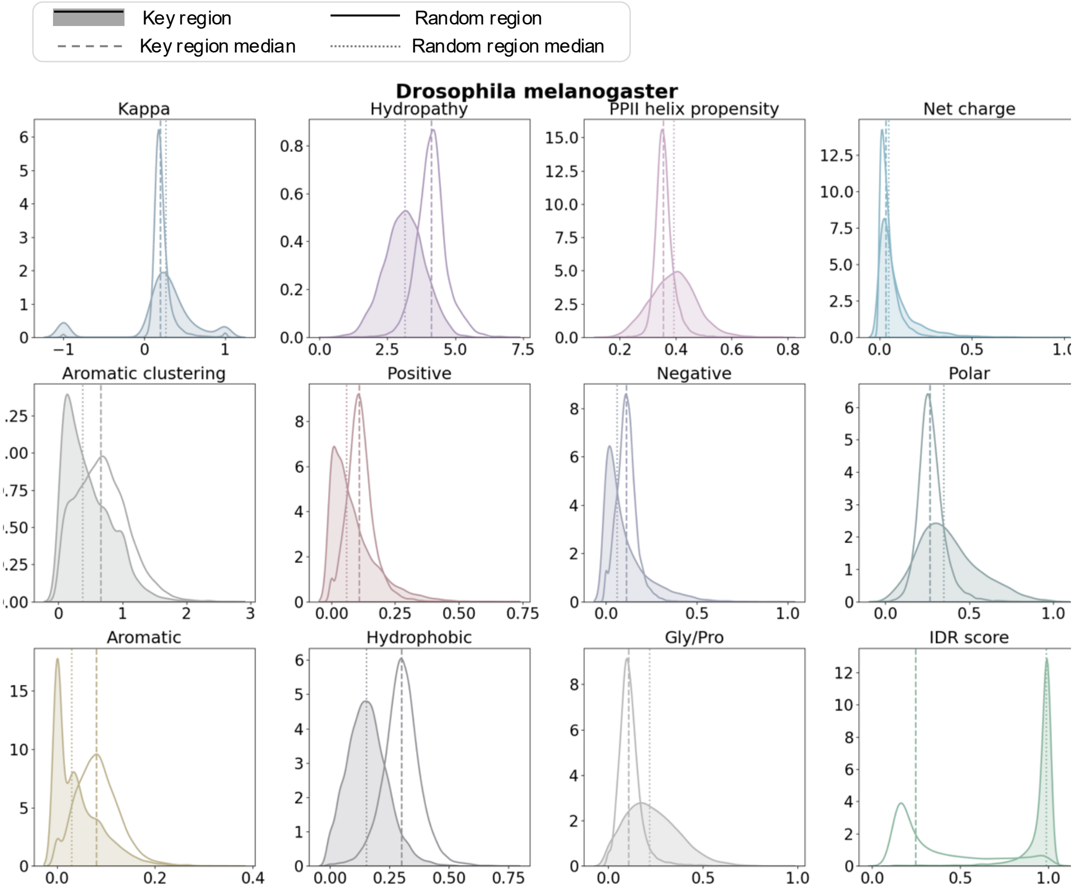


**Supplementary Figure 19. Physicochemical feature analysis in key and random regions of *Drosophila melanogaster*.** Density distributions of physicochemical features for random and key regions across all 18 proteomes analyzed in this study. Median for key and random regions are represented by dashed and dotted lines respectively.

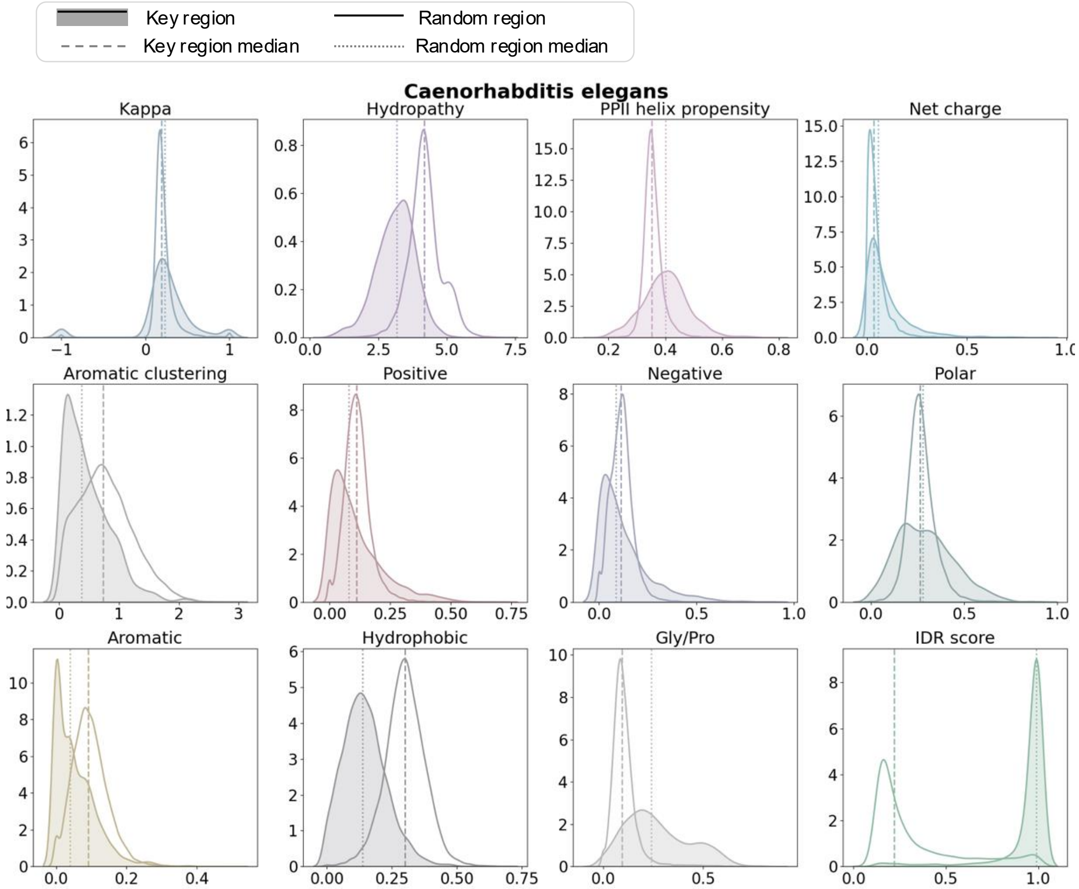


**Supplementary Figure 20. Physicochemical feature analysis in key and random regions of *Caenorhabditis elegans*.** Density distributions of physicochemical features for random and key regions across all 18 proteomes analyzed in this study. Median for key and random regions are represented by dashed and dotted lines respectively.

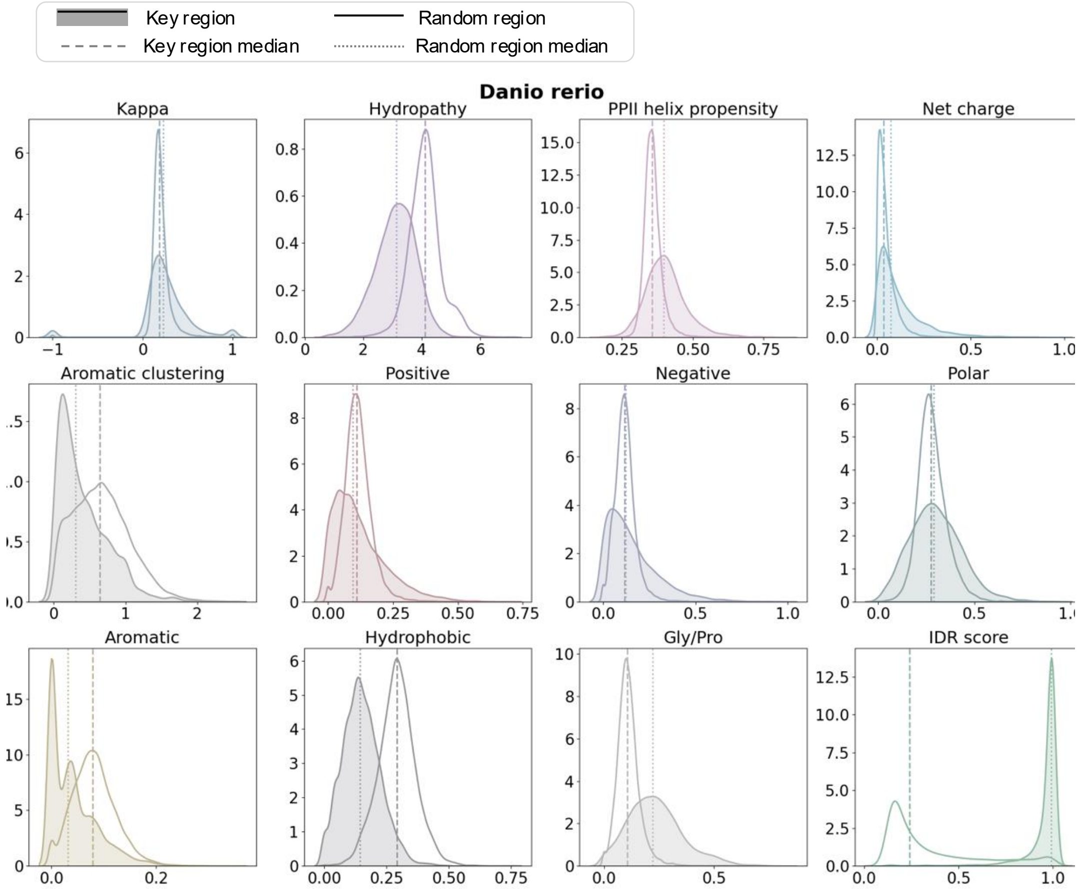


**Supplementary Figure 21. Physicochemical feature analysis in key and random regions of *Danio rerio*.** Density distributions of physicochemical features for random and key regions across all 18 proteomes analyzed in this study. Median for key and random regions are represented by dashed and dotted lines respectively.

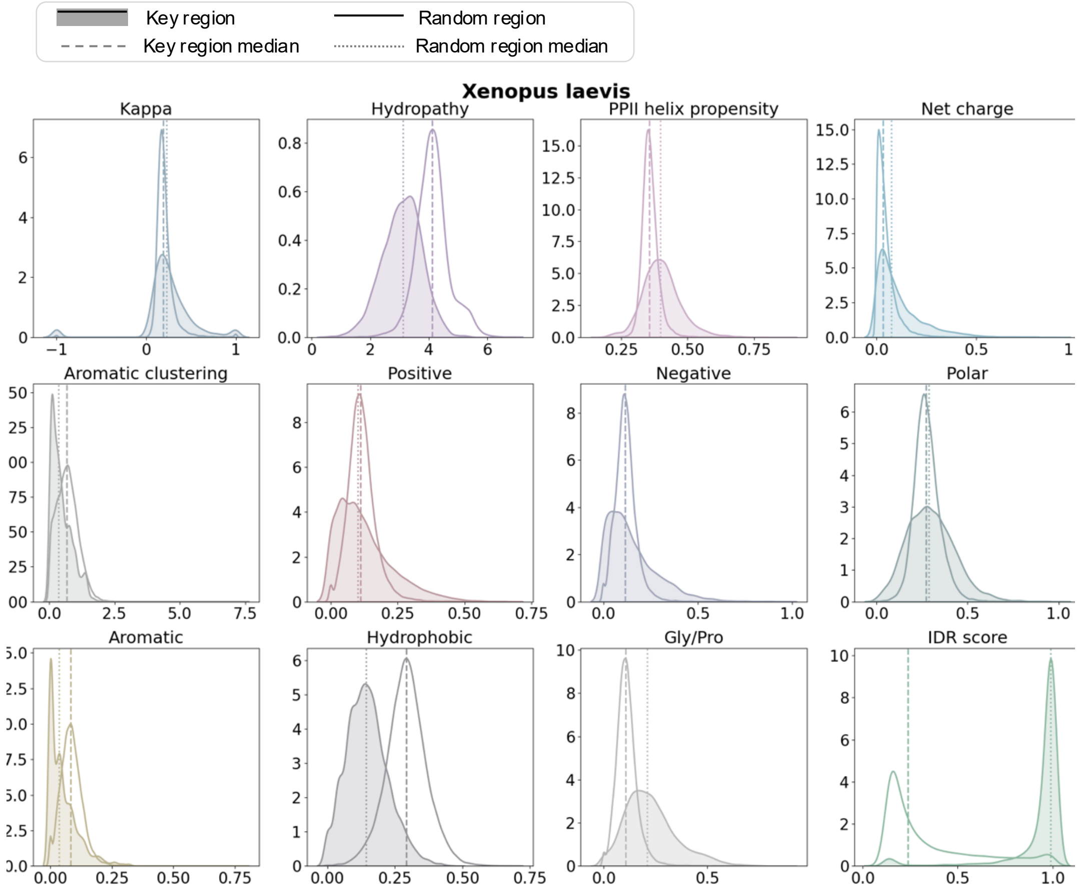


**Supplementary Figure 22. Physicochemical feature analysis in key and random regions of *Xenopus laevis*.** Density distributions of physicochemical features for random and key regions across all 18 proteomes analyzed in this study. Median for key and random regions are represented by dashed and dotted lines respectively.

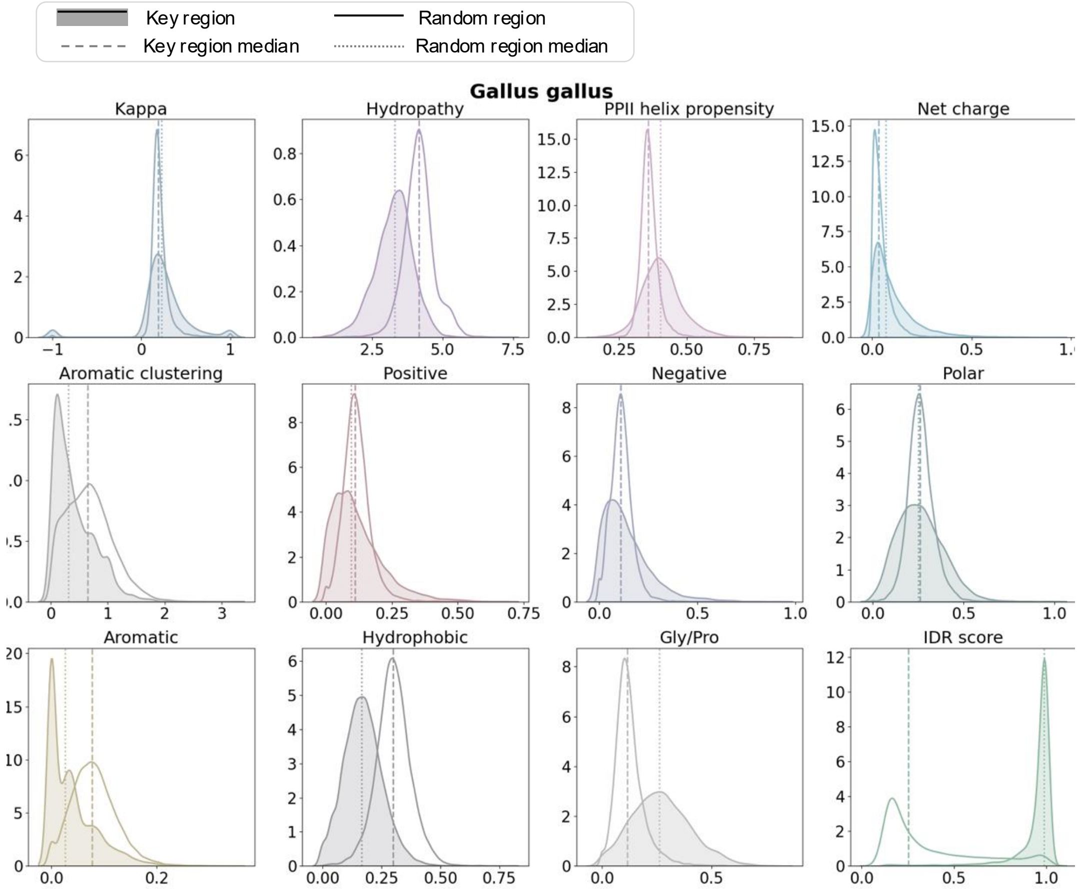


**Supplementary Figure 23. Physicochemical feature analysis in key and random regions of *Gallus gallus*.** Density distributions of physicochemical features for random and key regions across all 18 proteomes analyzed in this study. Median for key and random regions are represented by dashed and dotted lines respectively.

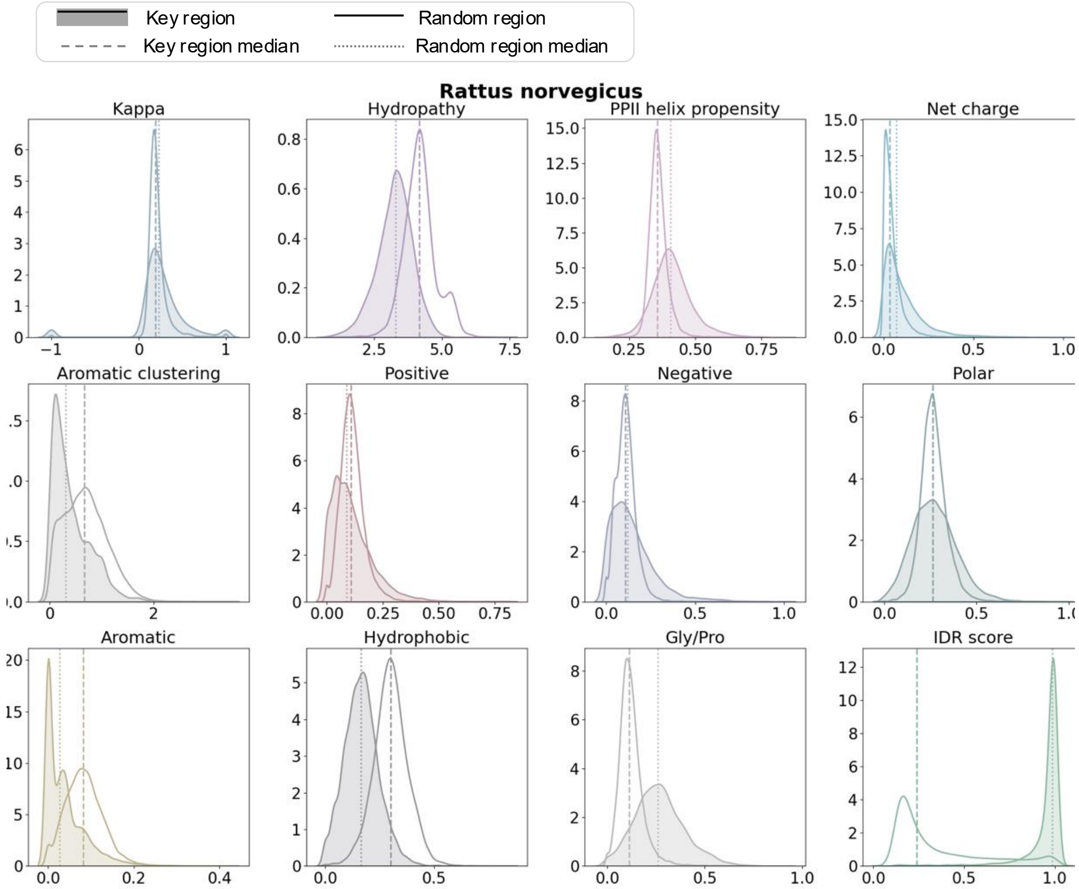


**Supplementary Figure 24. Physicochemical feature analysis in key and random regions of *Ratus norvegicus*.** Density distributions of physicochemical features for random and key regions across all 18 proteomes analyzed in this study. Median for key and random regions are represented by dashed and dotted lines respectively.

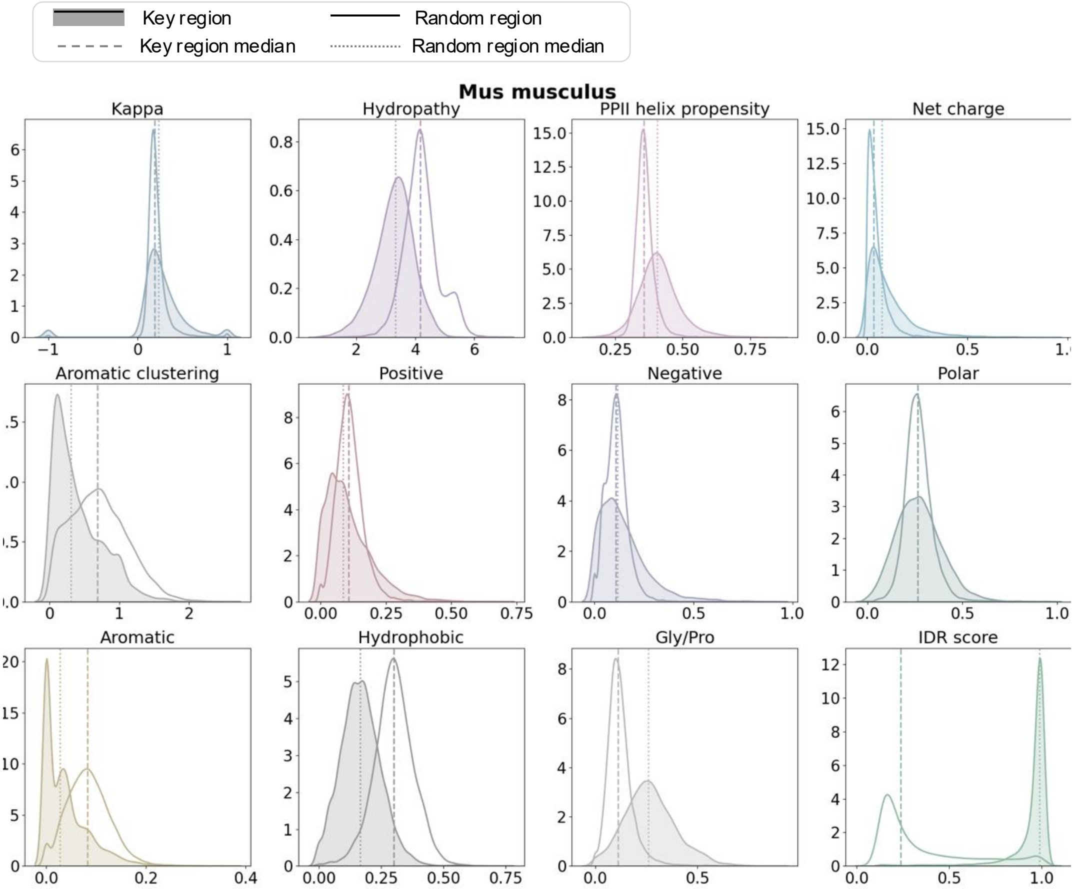


**Supplementary Figure 25. Physicochemical feature analysis in key and random regions of *Mus musculus*.** Density distributions of physicochemical features for random and key regions across all 18 proteomes analyzed in this study. Median for key and random regions are represented by dashed and dotted lines respectively.

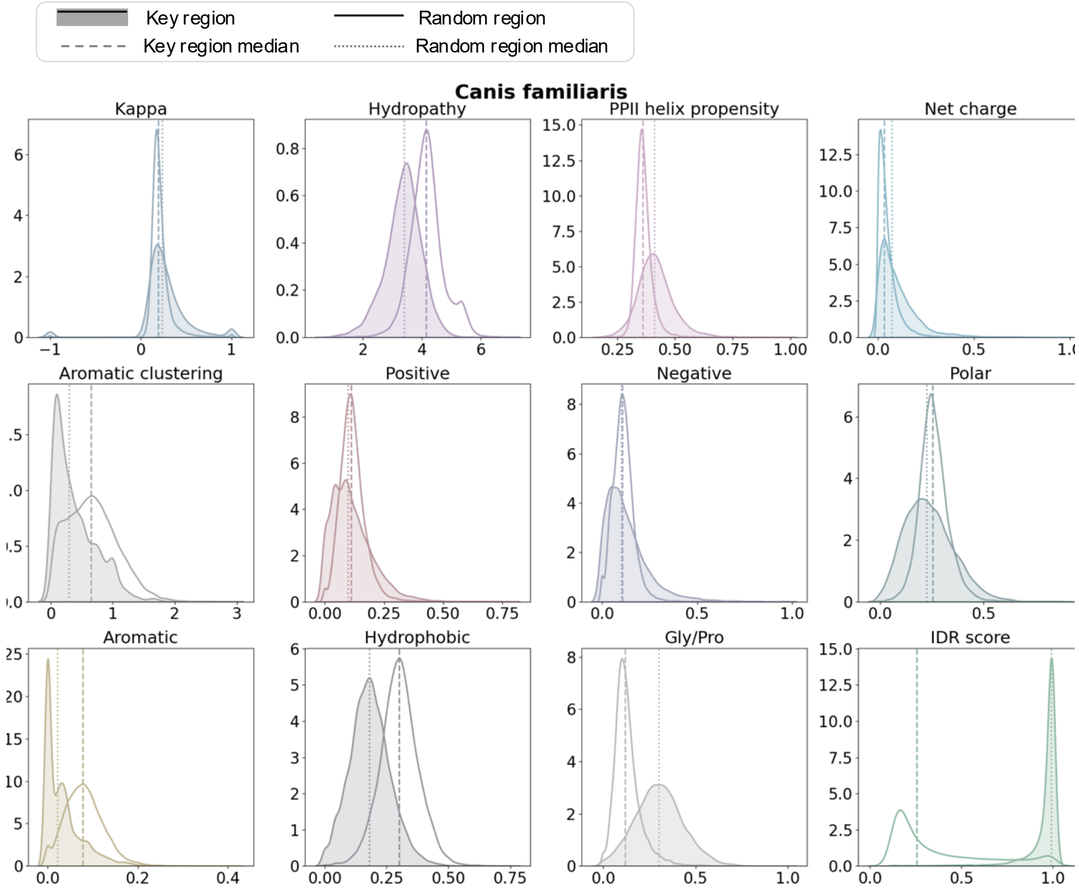


**Supplementary Figure 26. Physicochemical feature analysis in key and random regions of *Canis familiaris*.** Density distributions of physicochemical features for random and key regions across all 18 proteomes analyzed in this study. Median for key and random regions are represented by dashed and dotted lines respectively.

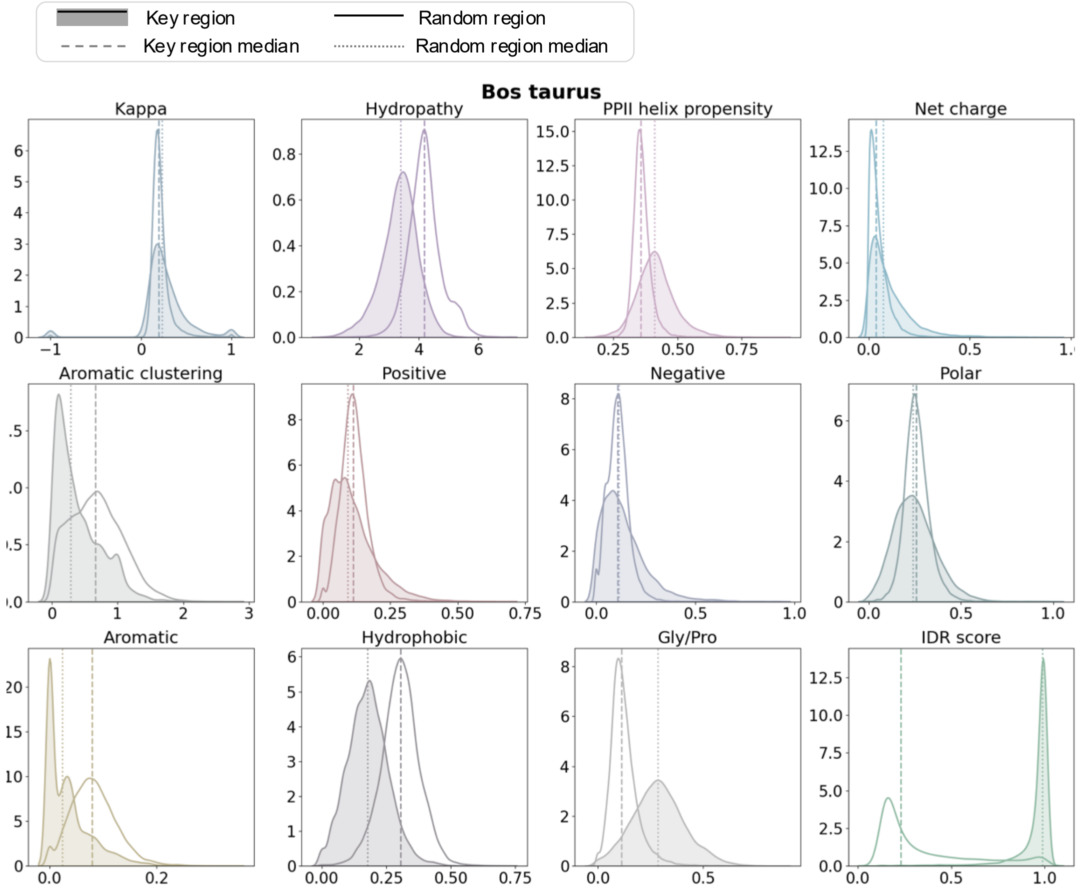


**Supplementary Figure 27. Physicochemical feature analysis in key and random regions of *Bos taurus*.** Density distributions of physicochemical features for random and key regions across all 18 proteomes analyzed in this study. Median for key and random regions are represented by dashed and dotted lines respectively.

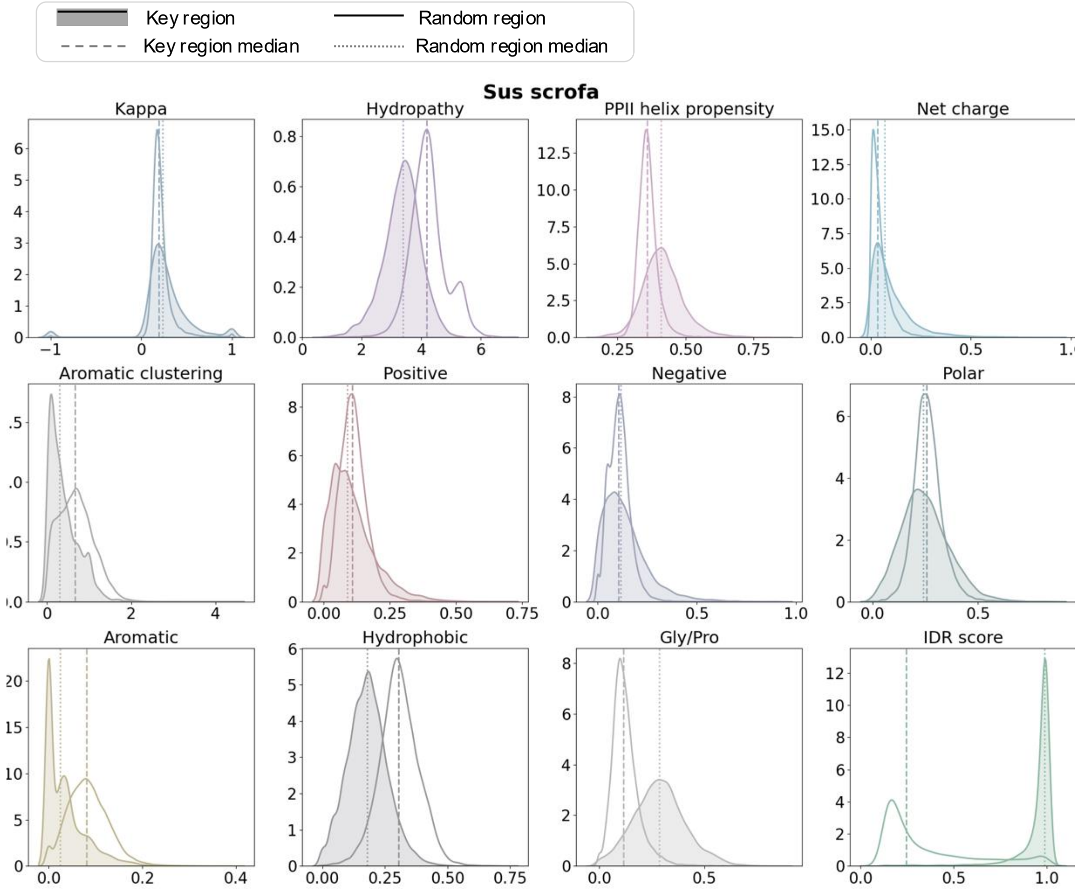


**Supplementary Figure 28. Physicochemical feature analysis in key and random regions of *Sus scrofa*.** Density distributions of physicochemical features for random and key regions across all 18 proteomes analyzed in this study. Median for key and random regions are represented by dashed and dotted lines respectively.

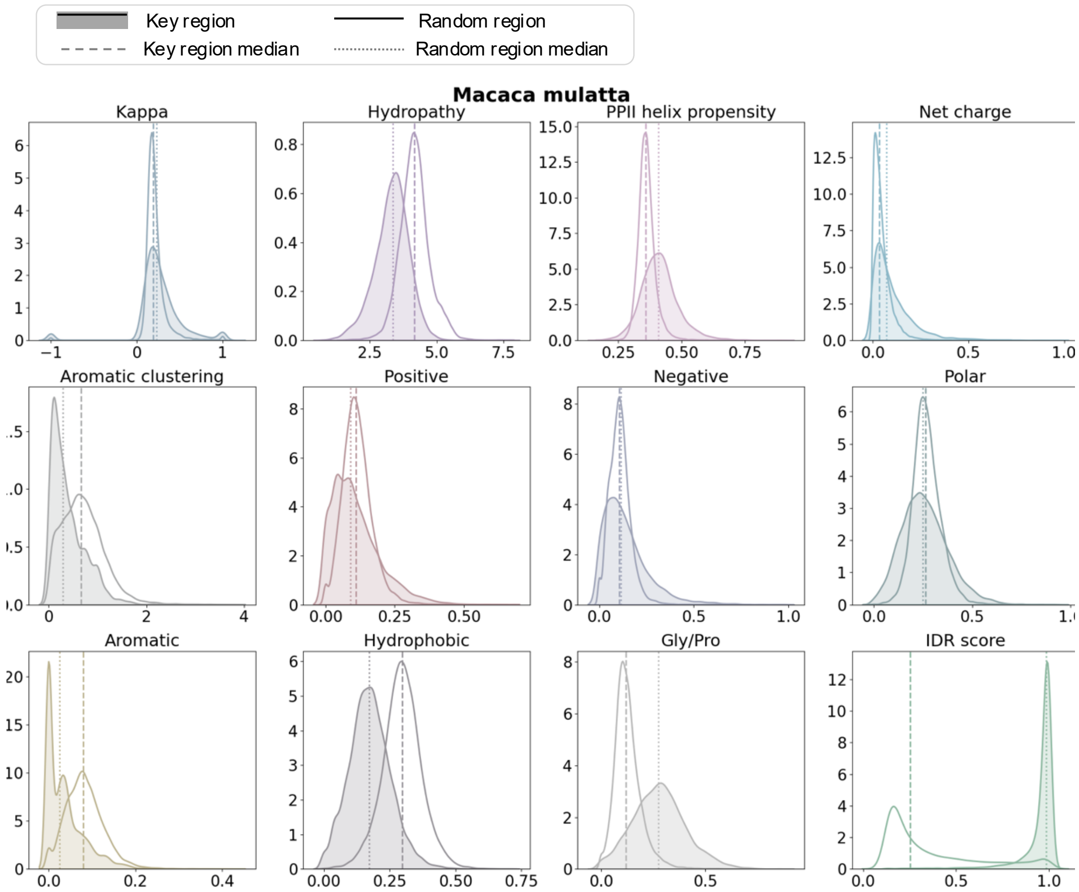


**Supplementary Figure 29. Physicochemical feature analysis in key and random regions of *Macaca mulatta*.** Density distributions of physicochemical features for random and key regions across all 18 proteomes analyzed in this study. Median for key and random regions are represented by dashed and dotted lines respectively.

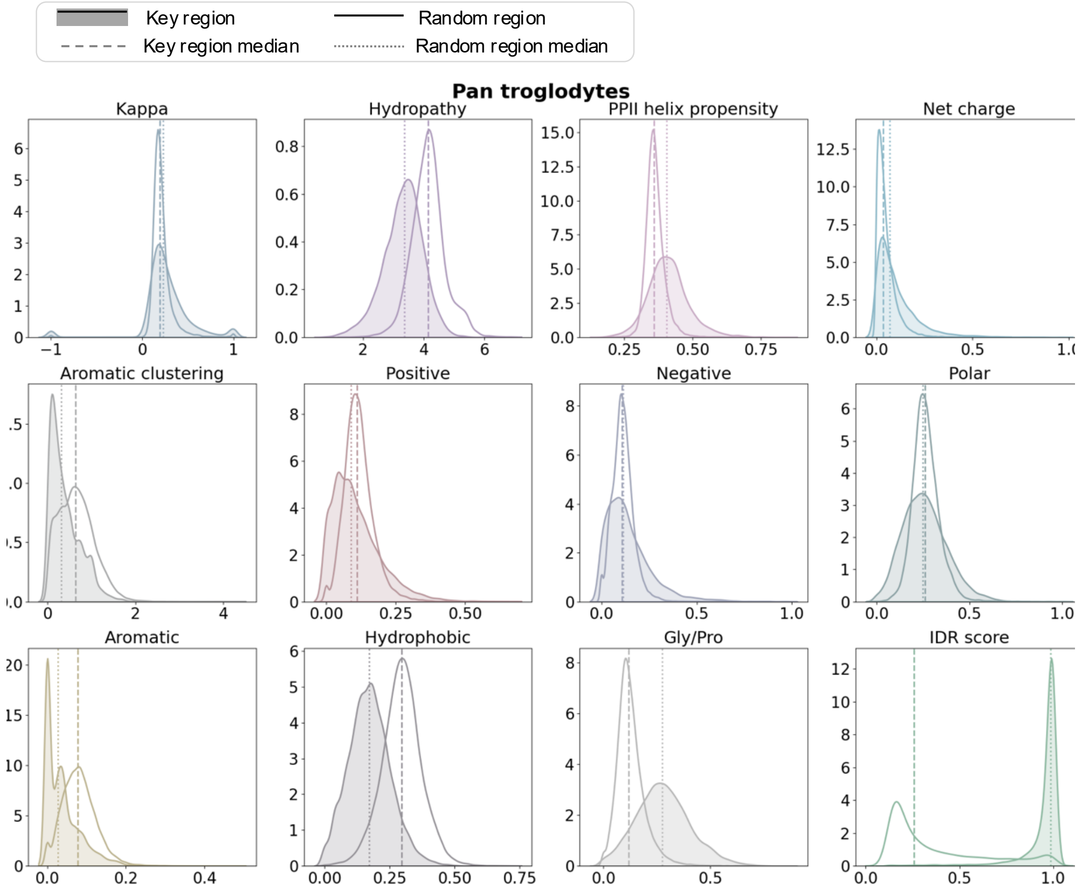


**Supplementary Figure 30. Physicochemical feature analysis in key and random regions of *Pan troglodytes*.** Density distributions of physicochemical features for random and key regions across all 18 proteomes analyzed in this study. Median for key and random regions are represented by dashed and dotted lines respectively.

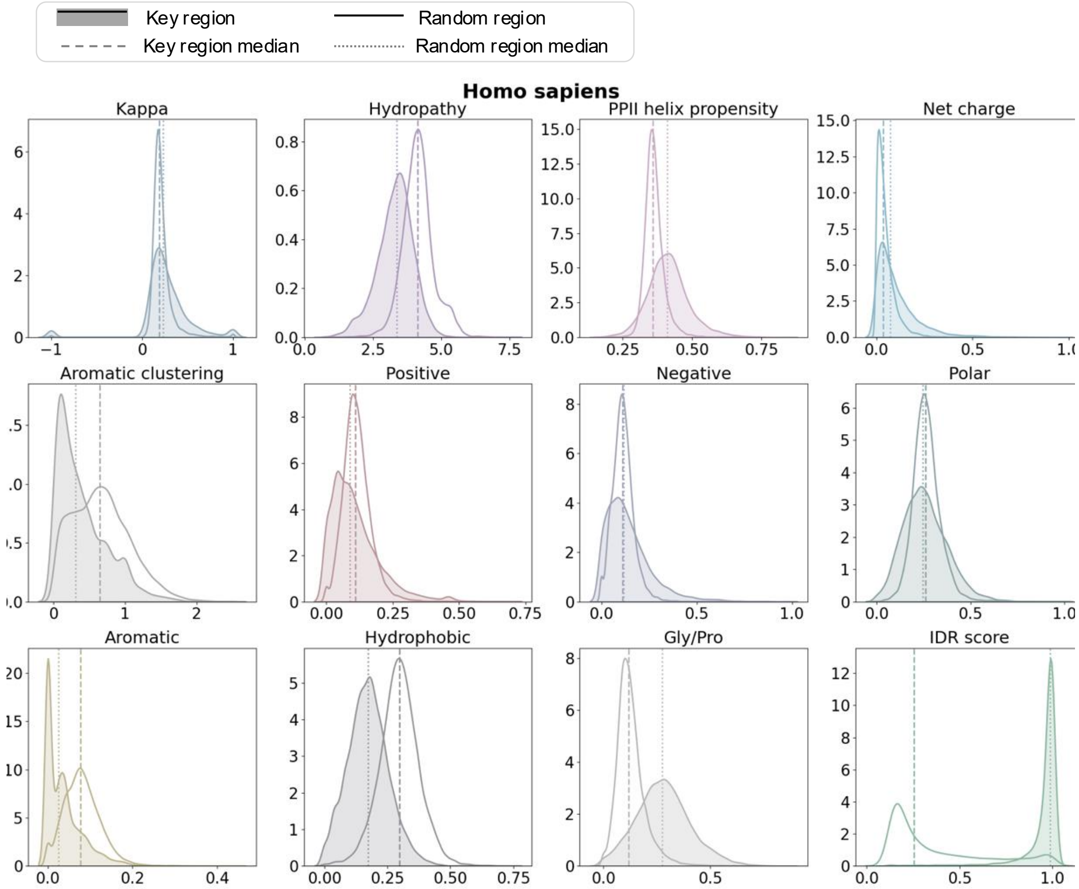


**Supplementary Figure 31. Physicochemical feature analysis in key and random regions of *Homo sapiens*.** Density distributions of physicochemical features for random and key regions across all 18 proteomes analyzed in this study. Median for key and random regions are represented by dashed and dotted lines respectively.

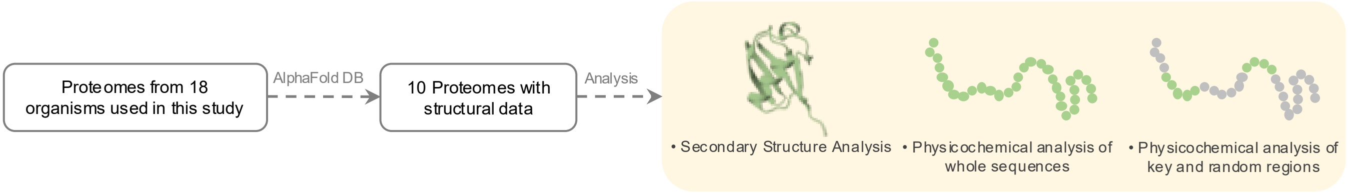


**Supplementary Figure 32. Structural features associated with LLPS score in key and random regions.** Only 10 out of the 18 proteomes used in this study had predicted secondary structures available in the AlphaFold Protein Structure Database (AlphaFold DB). We analyzed secondary structures (b), physicochemical features of full-length sequences (c), and physicochemical features of key versus random regions (d) to investigate how these properties correlate with the protein LLPS propensity.

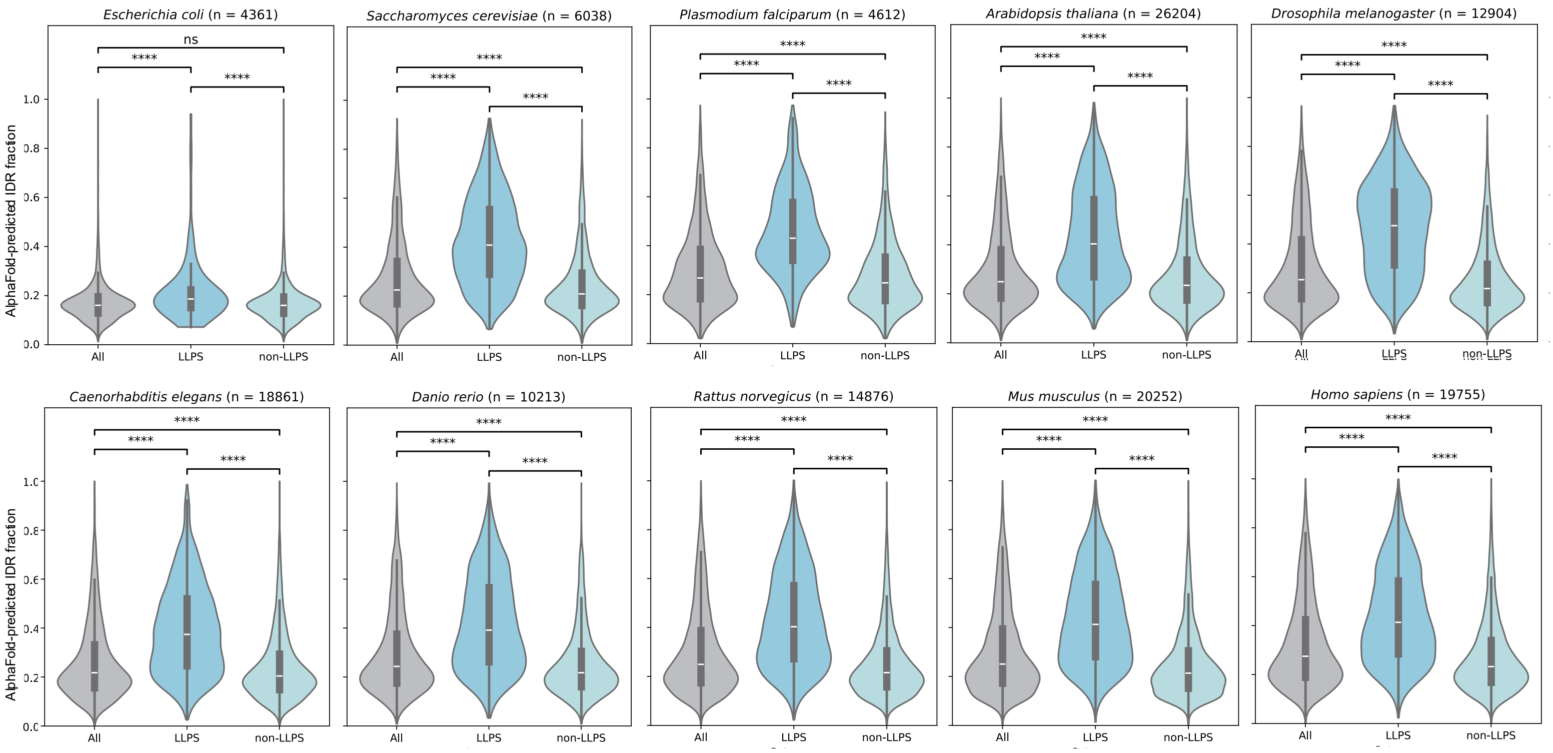


**Supplementary Figure 33. Species-specific AlphaFold-predicted IDR fraction distributions.** Violin plots show the distribution of IDR fractions across proteome sequences for ten species with available AlphaFold secondary structure predictions. For each proteome, sequences were classified as LLPS, non-LLPS, or All (LLPS + non-LLPS) based on Phaseek predictions. Proteins predicted to undergo phase separation (LLPS) exhibit significantly higher IDR fractions compared to non-LLPS proteins. Statistical significance was assessed using two-sided Mann–Whitney U tests and is indicated by asterisks (****$p < 0.0001$, ***$p < 0.001$, **$p < 0.01$, *$p < 0.05$; ns, not significant). n denotes the number of proteins in each proteome.

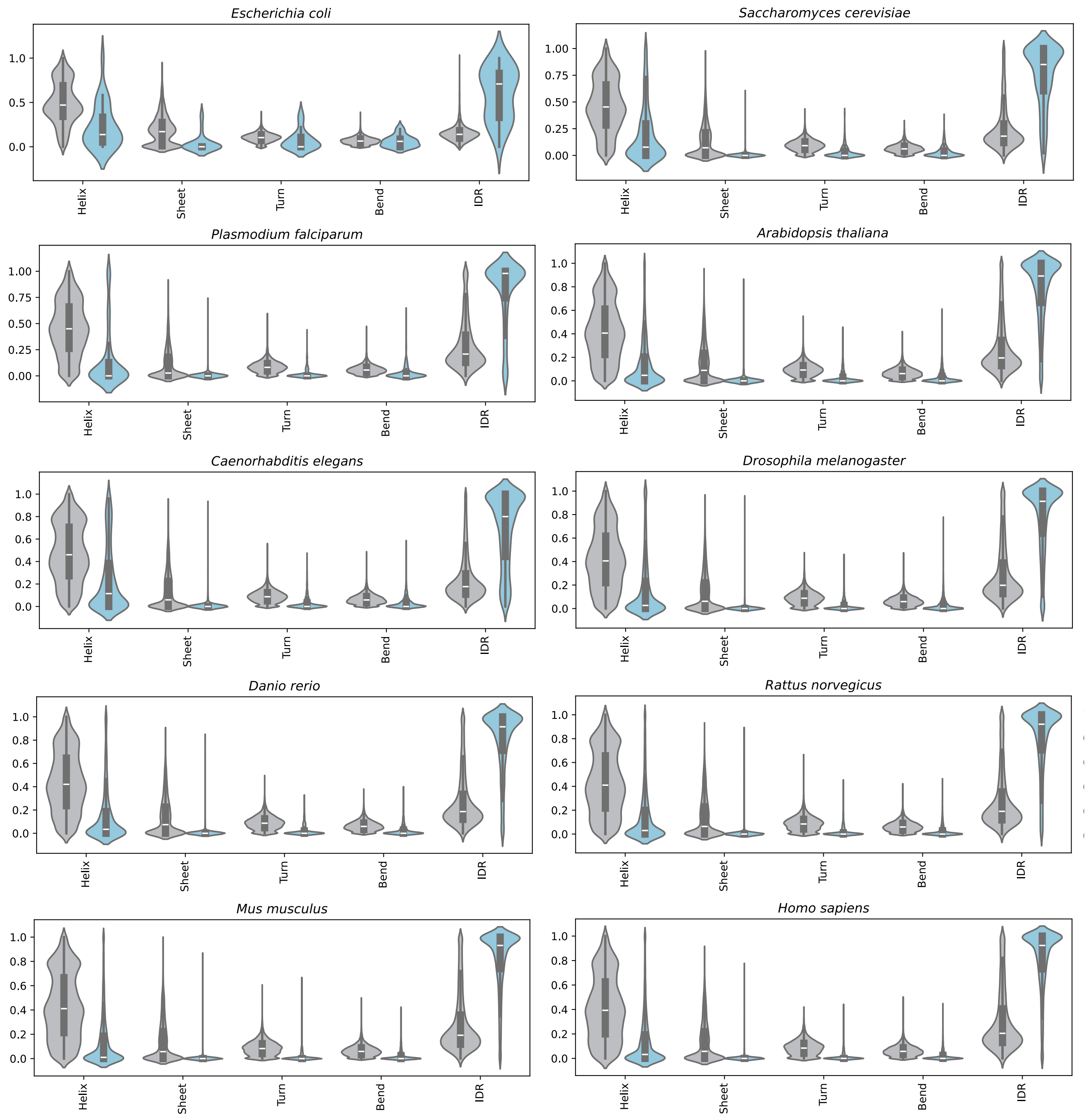


**Supplementary Figure 34: Secondary Structure Distribution of 10 Proteomes With Available Secondary Structures.** Violin plots representing for random and key regions, the distribution of secondary structures of 10 model organisms (AlphaFold-predicted structures). The one-sided Mann-Whitney U test was used for random and key region secondary structure distribution comparison, with ****p-value < 0.0001 (~0) for secondary structures of all organisms (not shown), except *E. coli*. The boxes show the interquartile range between the first and third quartiles, with the median as the middle line. Significance levels are: *p < 0.05, **p < 0.01, ***p < 0.001, ****p < 0.0001 (n.s. = not significant)

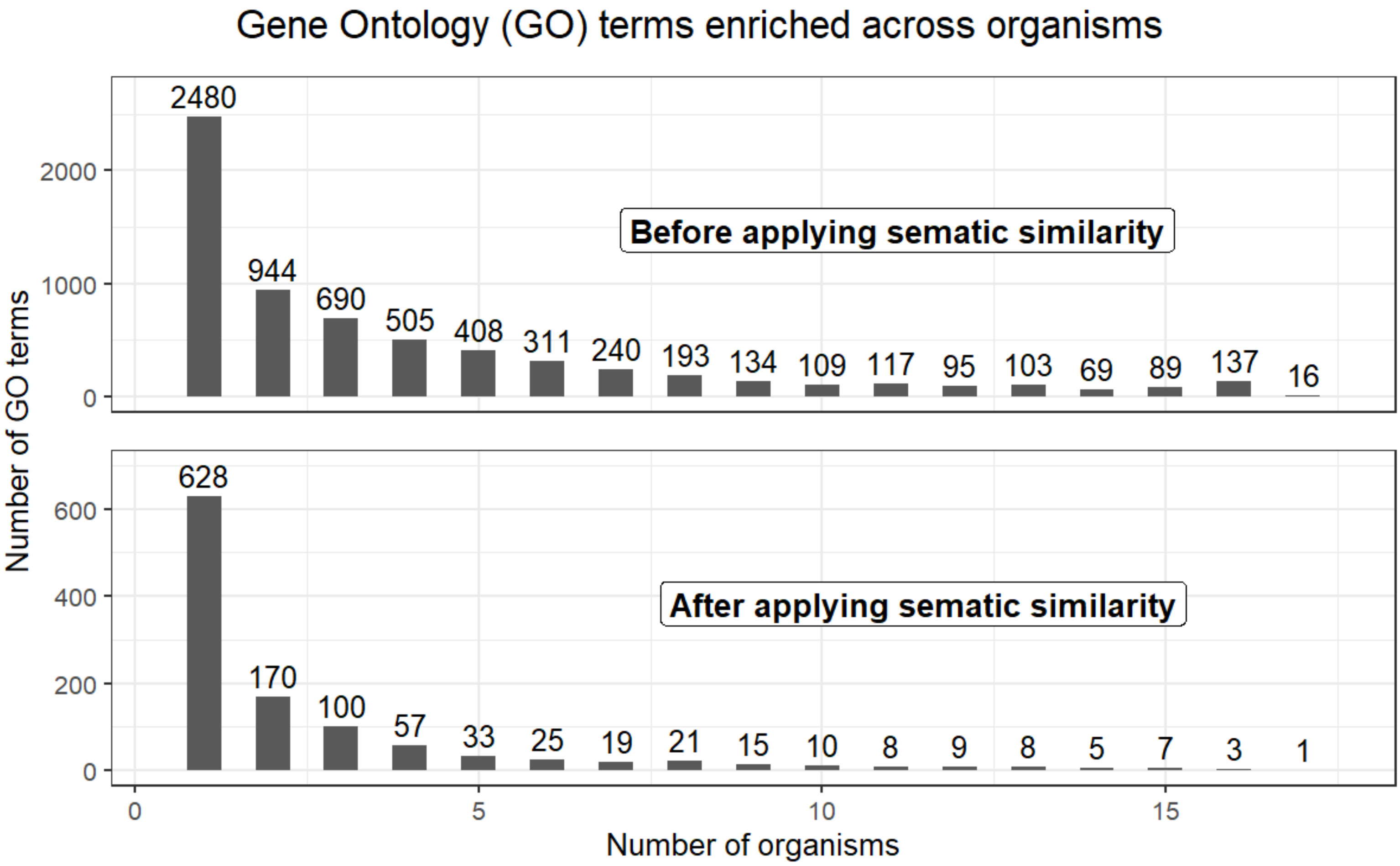


**Supplementary Figure 35: Distributions of Gene Ontology (GO) terms enriched across organisms before and after summarizing the results using semantic similarity.** Upper plot, histogram of enriched GO terms across organisms before applying semantic similarity. Lower plot, histogram of summarized results across organisms.

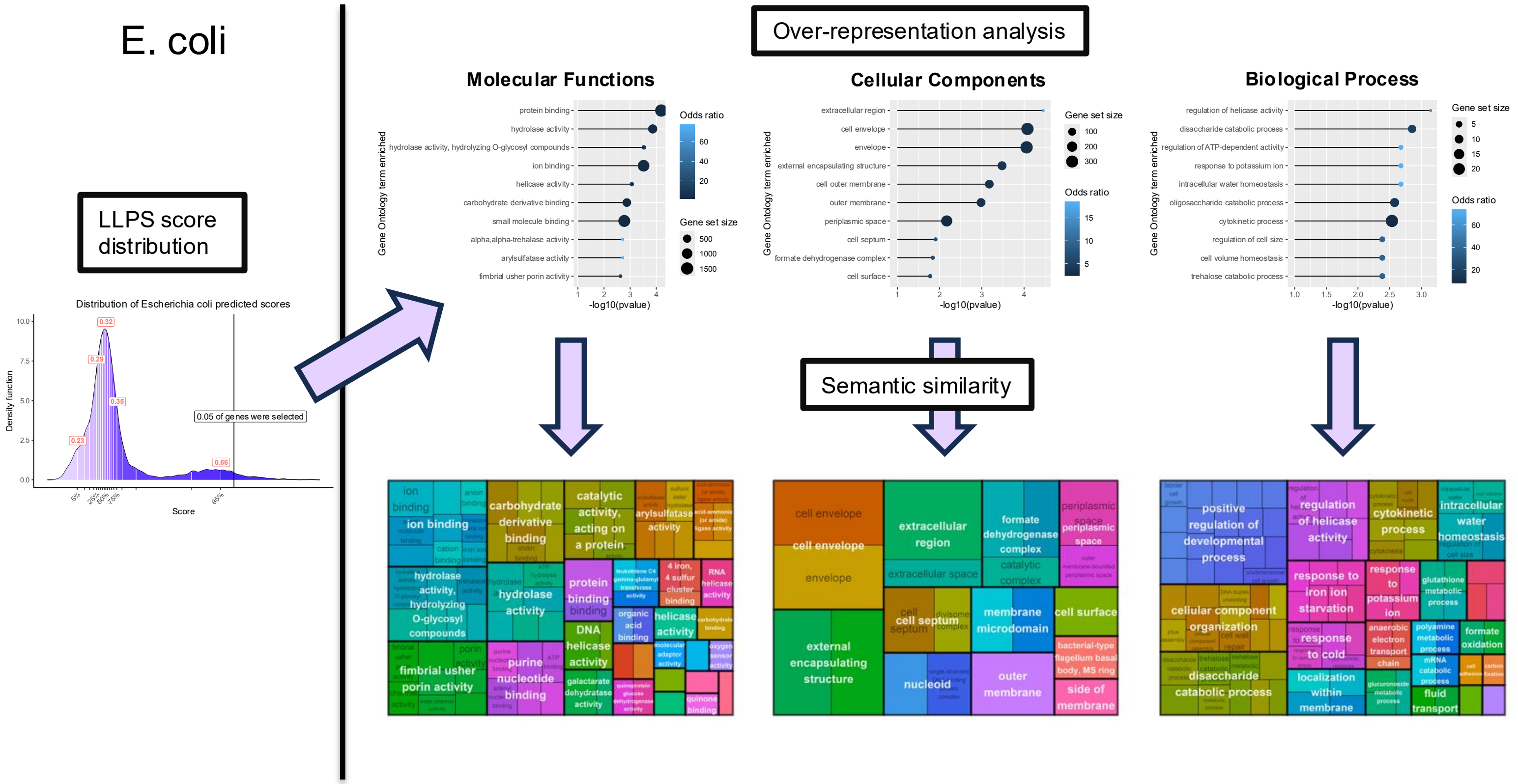


**Supplementary Figure 36: Gene Ontology (GO) enrichment results for Escherichia coli.** Left plot, distribution of LLPS scores within the proteome and fraction of total Coding Sequences (CDS) selected. Upper panel, lollipop plots representing the top-10 enriched terms for the three categories: Molecular Function, Cellular Component, Biological Process. Lower panel, treemap plots illustrating hierarchical relationships between GO terms and representative subset selected.

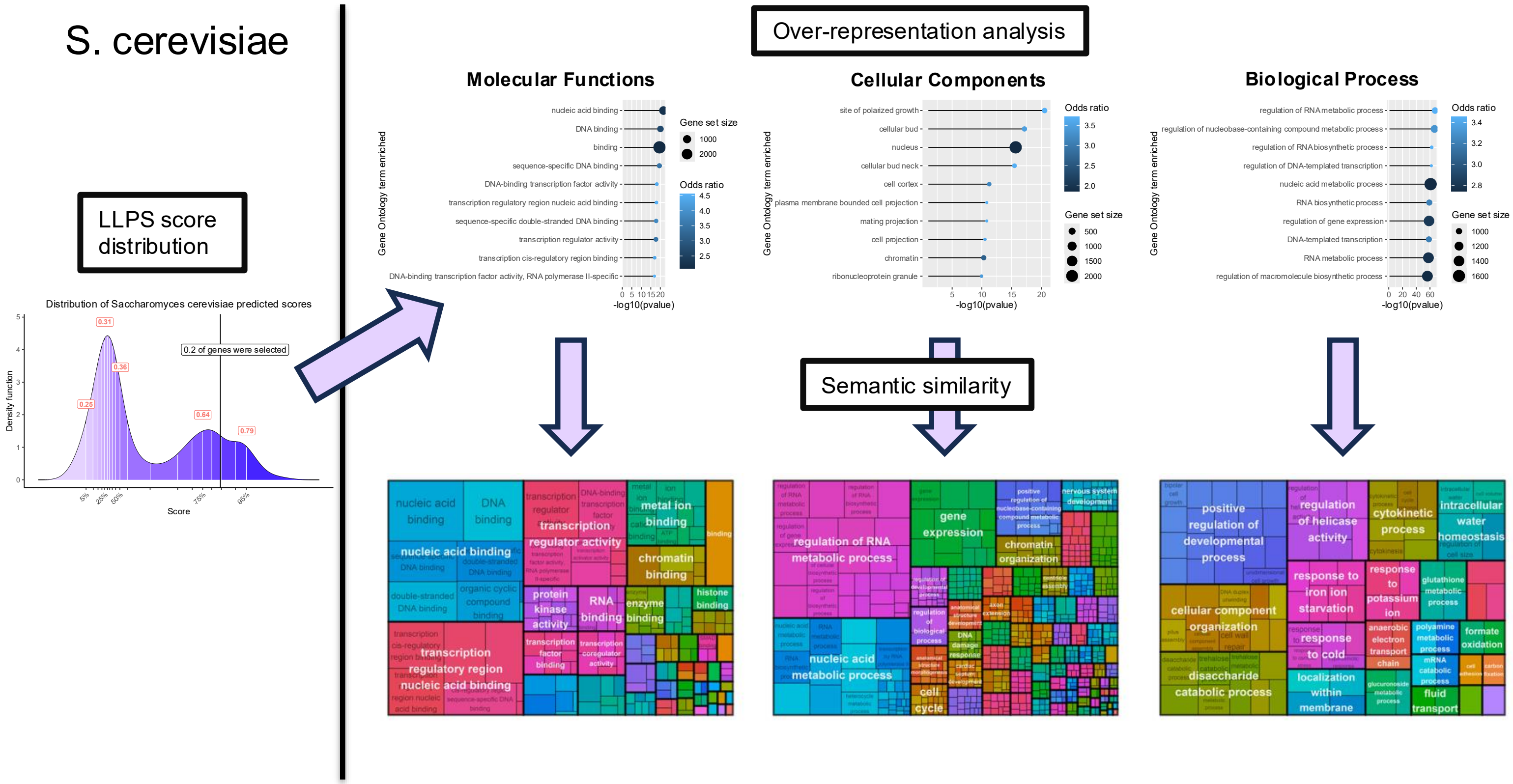


**Supplementary Figure 37: Gene Ontology (GO) enrichment results for Saccharomyces cerevisiae.** Left plot, distribution of LLPS scores within the proteome and fraction of total Coding Sequences (CDS) selected. Upper panel, lollipop plots representing the top-10 enriched terms for the three categories: Molecular Function, Cellular Component, Biological Process. Lower panel, treemap plots illustrating hierarchical relationships between GO terms and representative subset selected.

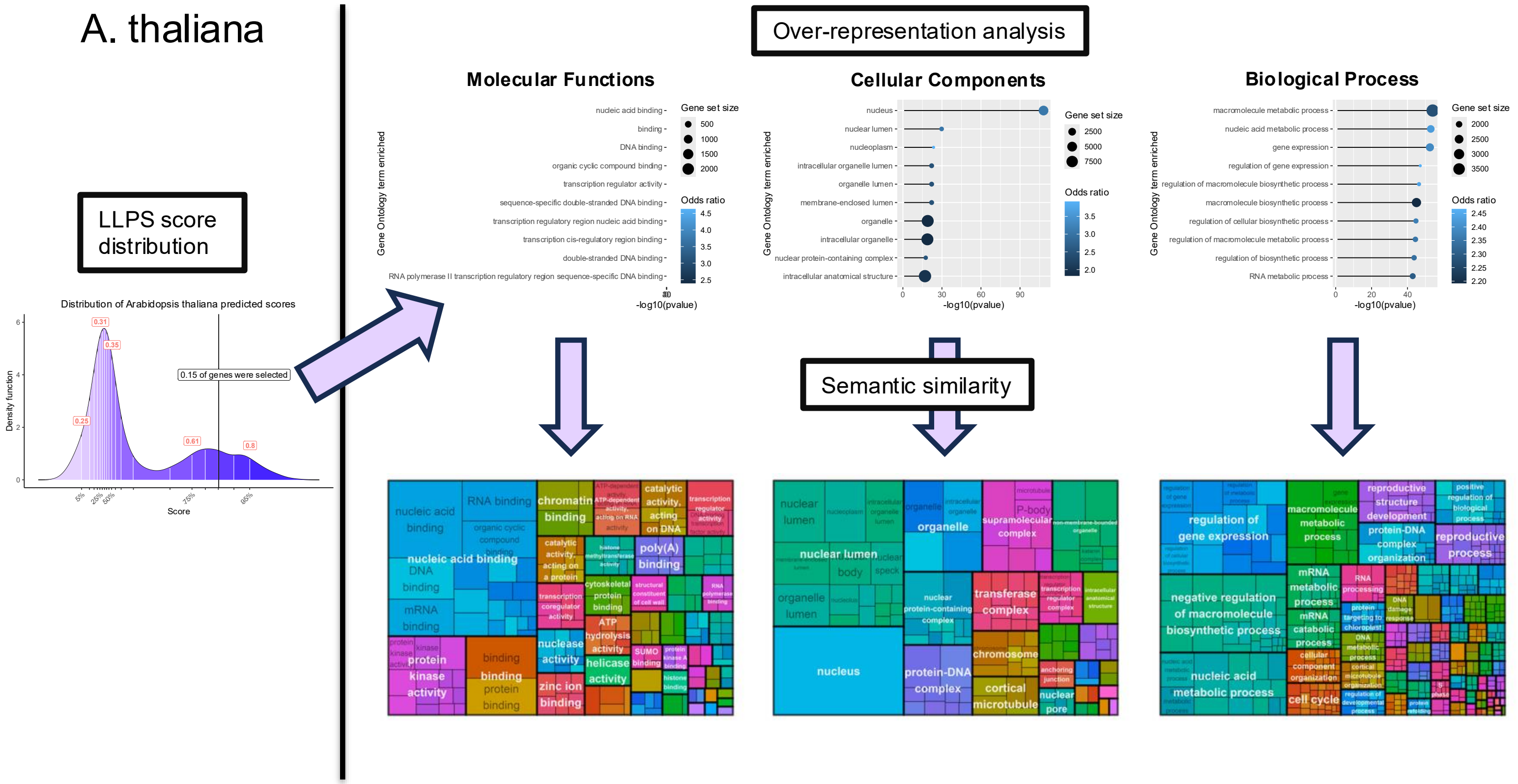


**Supplementary Figure 38: Gene Ontology (GO) enrichment results for Arabidopsis thaliana.** Left plot, distribution of LLPS scores within the proteome and fraction of total Coding Sequences (CDS) selected. Upper panel, lollipop plots representing the top-10 enriched terms for the three categories: Molecular Function, Cellular Component, Biological Process. Lower panel, treemap plots illustrating hierarchical relationships between GO terms and representative subset selected.

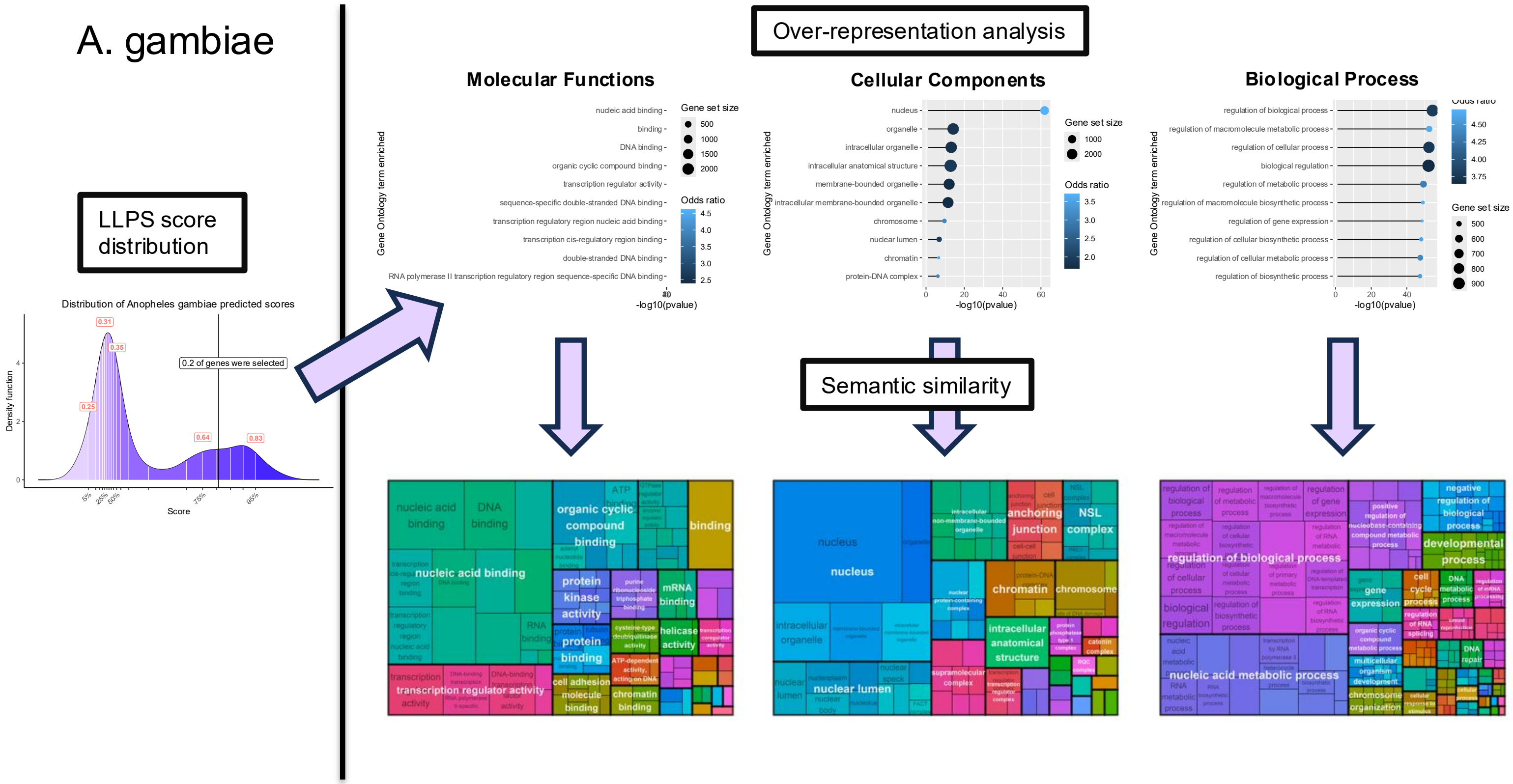


**Supplementary Figure 39: Gene Ontology (GO) enrichment results for Anopheles gambiae.** Left plot, distribution of LLPS scores within the proteome and fraction of total Coding Sequences (CDS) selected. Upper panel, lollipop plots representing the top-10 enriched terms for the three categories: Molecular Function, Cellular Component, Biological Process. Lower panel, treemap plots illustrating hierarchical relationships between GO terms and representative subset selected.

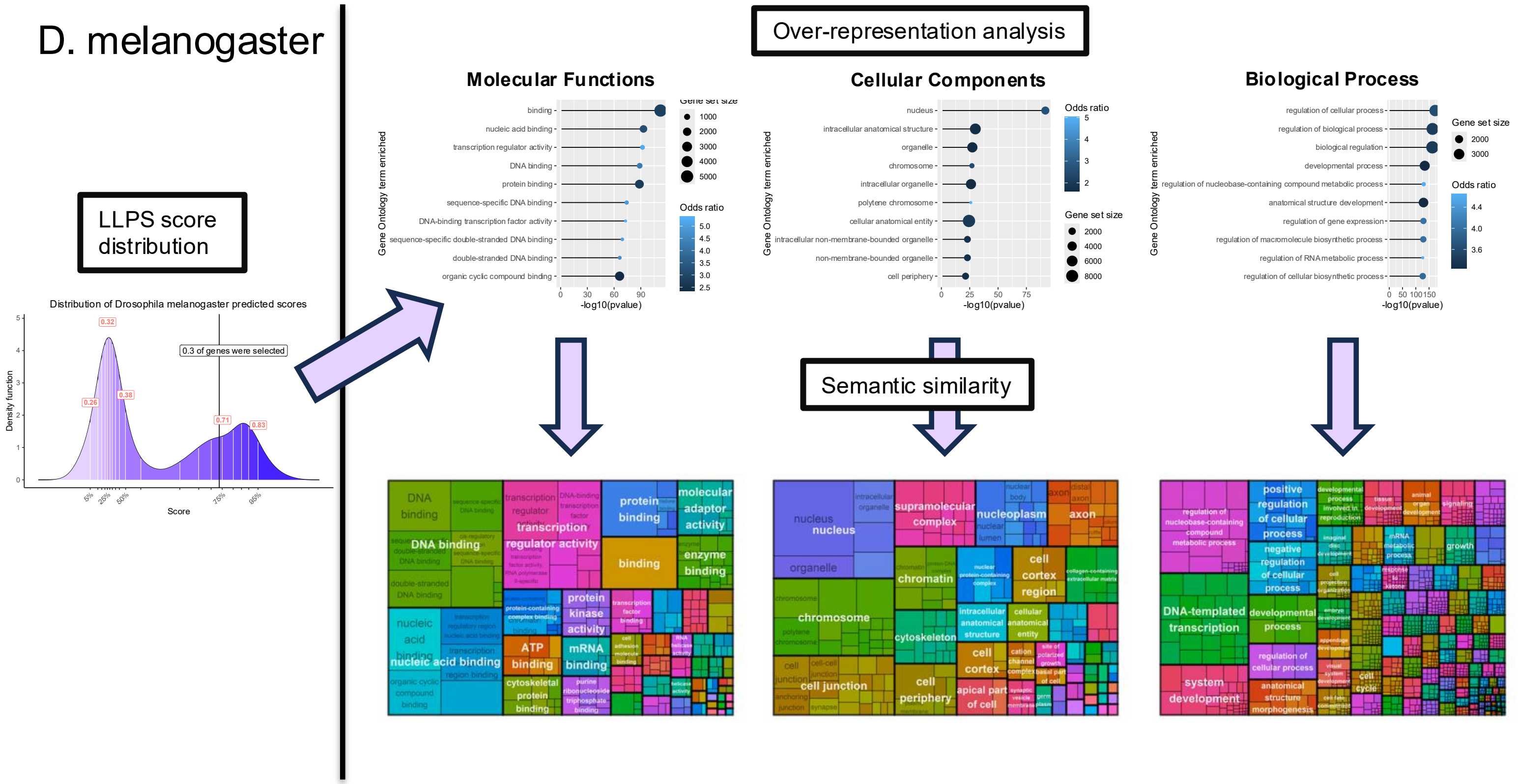


**Supplementary Figure 40: Gene Ontology (GO) enrichment results for Drosophila melanogaster.** Left plot, distribution of LLPS scores within the proteome and fraction of total Coding Sequences (CDS) selected. Upper panel, lollipop plots representing the top-10 enriched terms for the three categories: Molecular Function, Cellular Component, Biological Process. Lower panel, treemap plots illustrating hierarchical relationships between GO terms and representative subset selected.

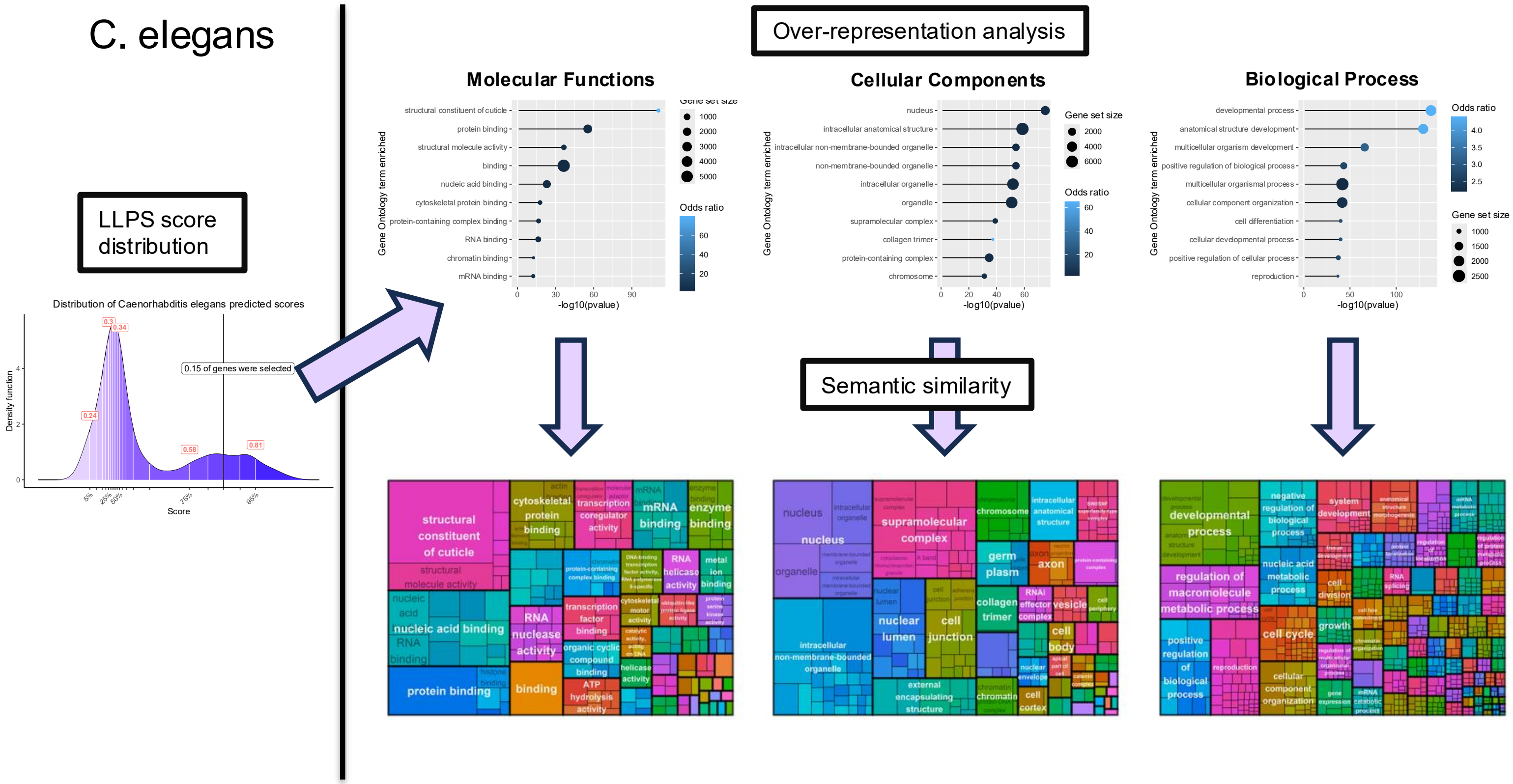


**Supplementary Figure 41: Gene Ontology (GO) enrichment results for Caenorhabditis elegans.** Left plot, distribution of LLPS scores within the proteome and fraction of total Coding Sequences (CDS) selected. Upper panel, lollipop plots representing the top-10 enriched terms for the three categories: Molecular Function, Cellular Component, Biological Process. Lower panel, treemap plots illustrating hierarchical relationships between GO terms and representative subset selected.

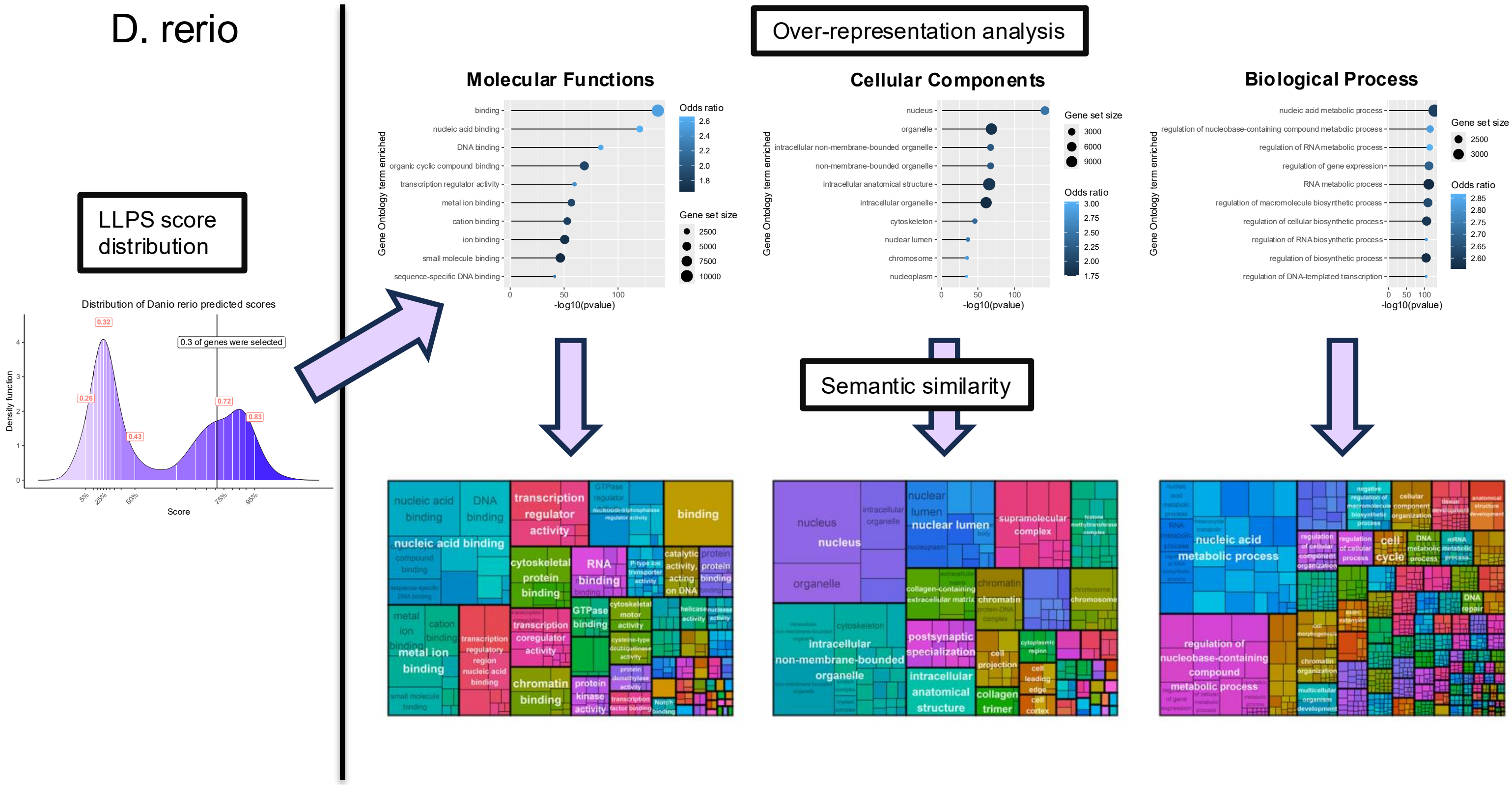


**Supplementary Figure 42: Gene Ontology (GO) enrichment results for Danio rerio.** Left plot, distribution of LLPS scores within the proteome and fraction of total Coding Sequences (CDS) selected. Upper panel, lollipop plots representing the top-10 enriched terms for the three categories: Molecular Function, Cellular Component, Biological Process. Lower panel, treemap plots illustrating hierarchical relationships between GO terms and representative subset selected.

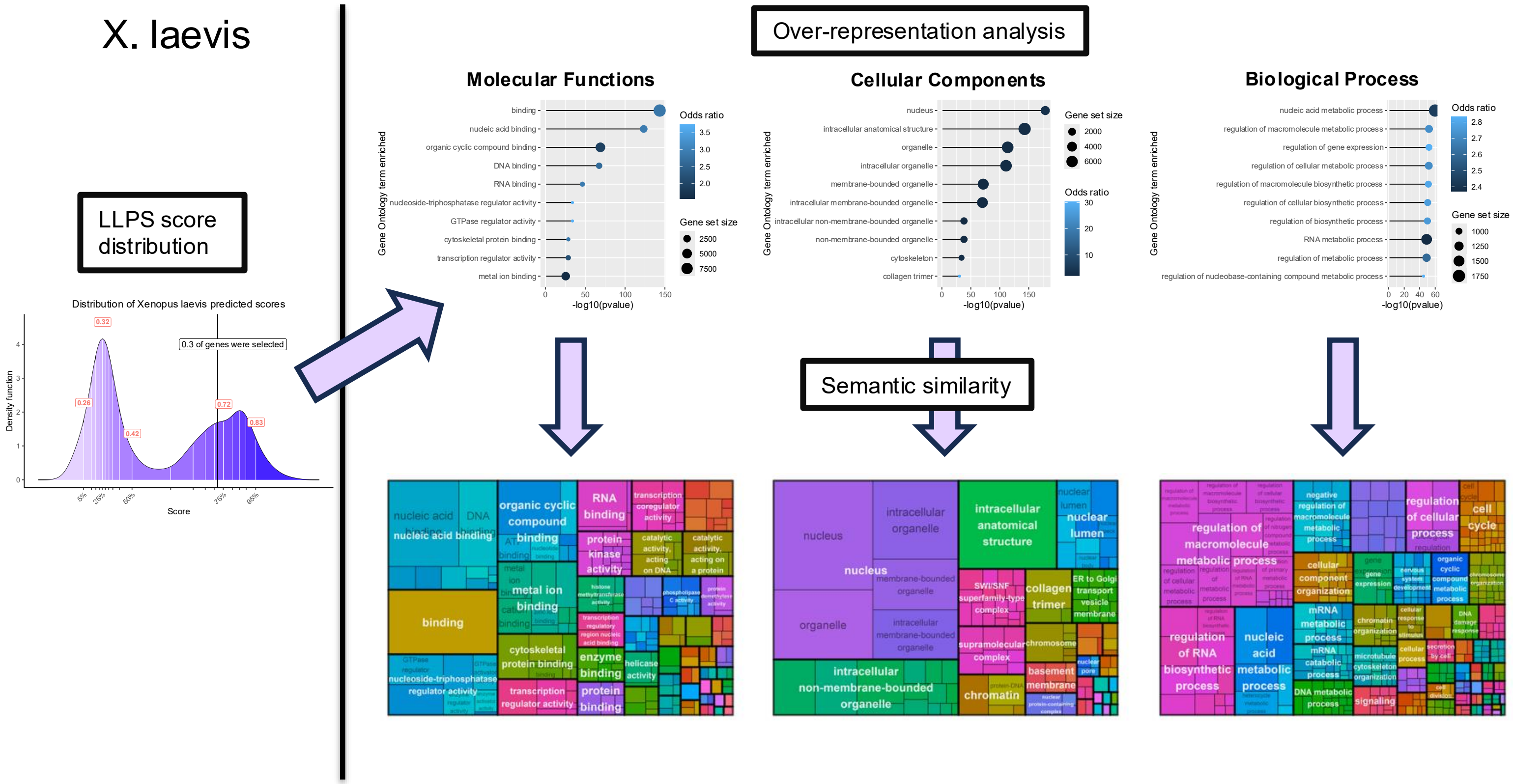


**Supplementary Figure 43: Gene Ontology (GO) enrichment results for Xenopus laevis.** Left plot, distribution of LLPS scores within the proteome and fraction of total Coding Sequences (CDS) selected. Upper panel, lollipop plots representing the top-10 enriched terms for the three categories: Molecular Function, Cellular Component, Biological Process. Lower panel, treemap plots illustrating hierarchical relationships between GO terms and representative subset selected.

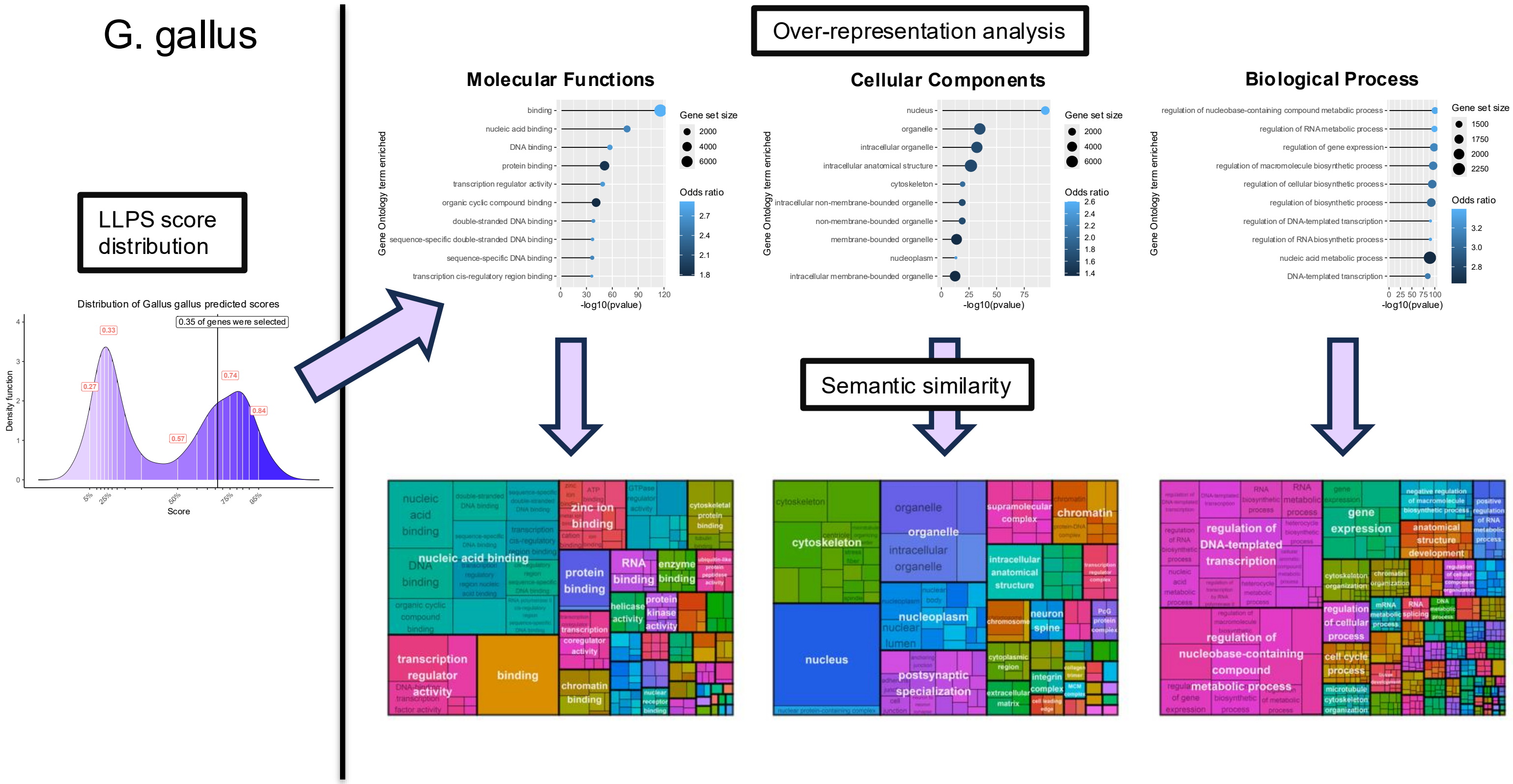


**Supplementary Figure 44: Gene Ontology (GO) enrichment results for Gallus gallus.** Left plot, distribution of LLPS scores within the proteome and fraction of total Coding Sequences (CDS) selected. Upper panel, lollipop plots representing the top-10 enriched terms for the three categories: Molecular Function, Cellular Component, Biological Process. Lower panel, treemap plots illustrating hierarchical relationships between GO terms and representative subset selected.

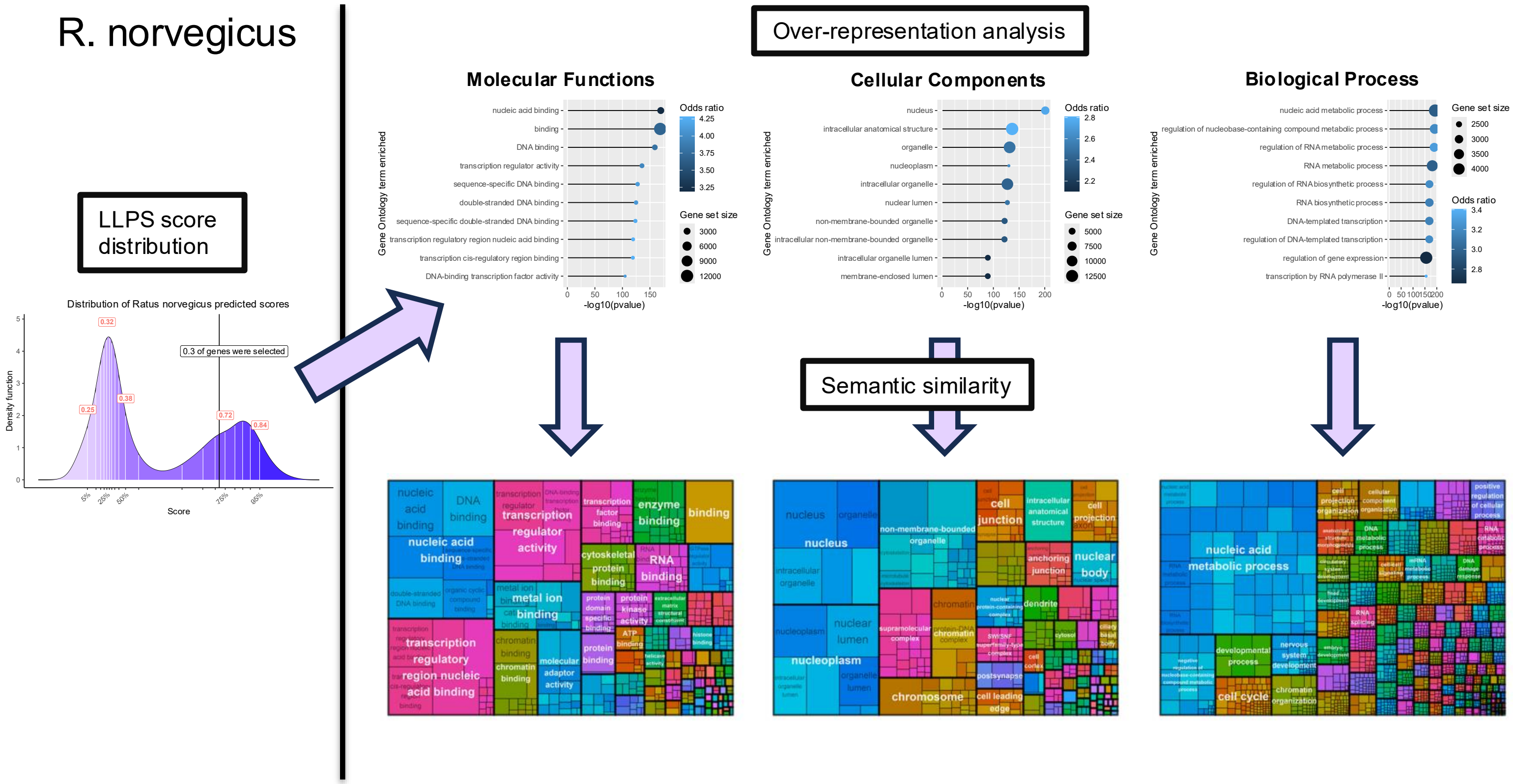


**Supplementary Figure 45: Gene Ontology (GO) enrichment results for Ratus norvegicus.** Left plot, distribution of LLPS scores within the proteome and fraction of total Coding Sequences (CDS) selected. Upper panel, lollipop plots representing the top-10 enriched terms for the three categories: Molecular Function, Cellular Component, Biological Process. Lower panel, treemap plots illustrating hierarchical relationships between GO terms and representative subset selected.

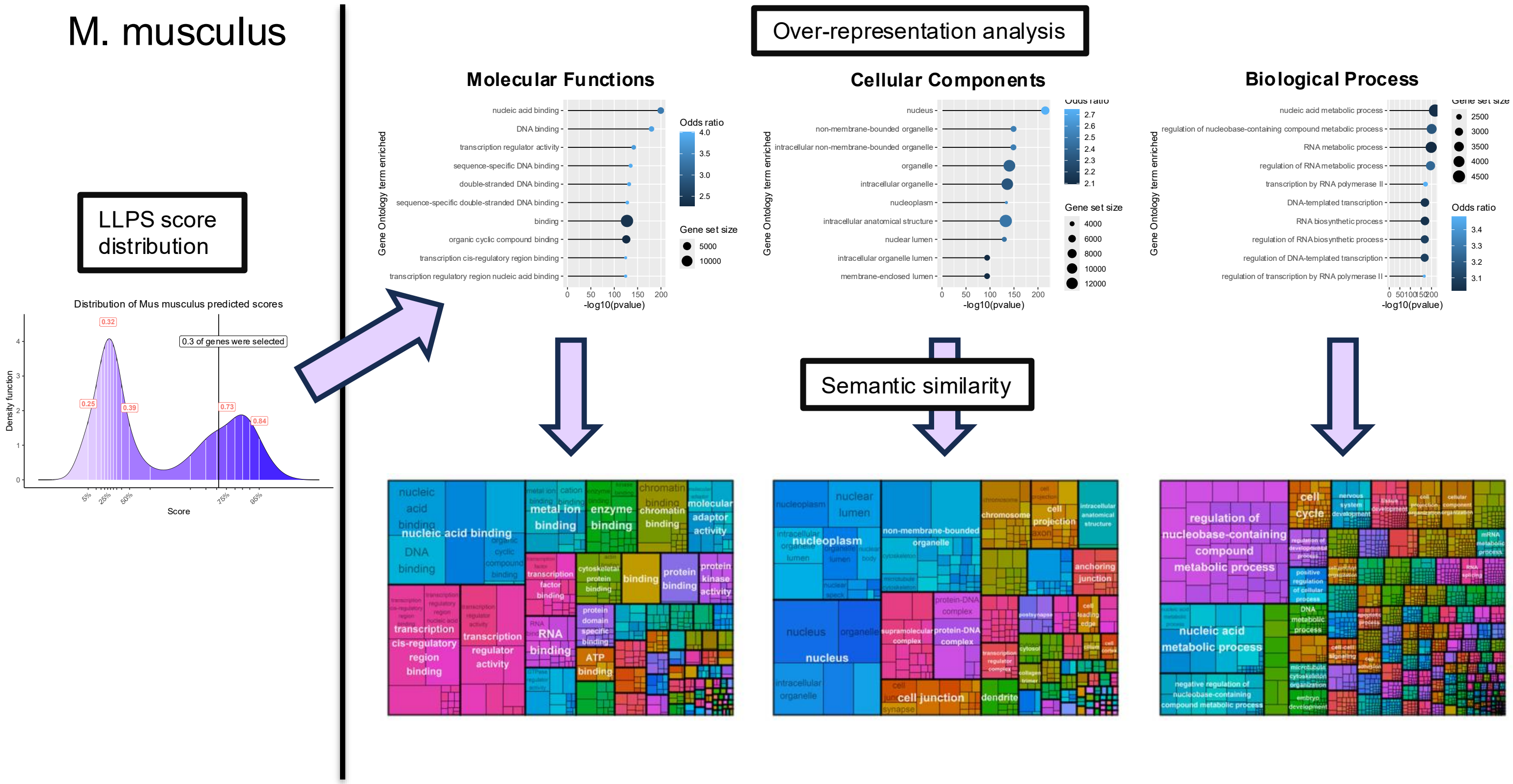


**Supplementary Figure 46: Gene Ontology (GO) enrichment results for Mus musculus.** Left plot, distribution of LLPS scores within the proteome and fraction of total Coding Sequences (CDS) selected. Upper panel, lollipop plots representing the top-10 enriched terms for the three categories: Molecular Function, Cellular Component, Biological Process. Lower panel, treemap plots illustrating hierarchical relationships between GO terms and representative subset selected.

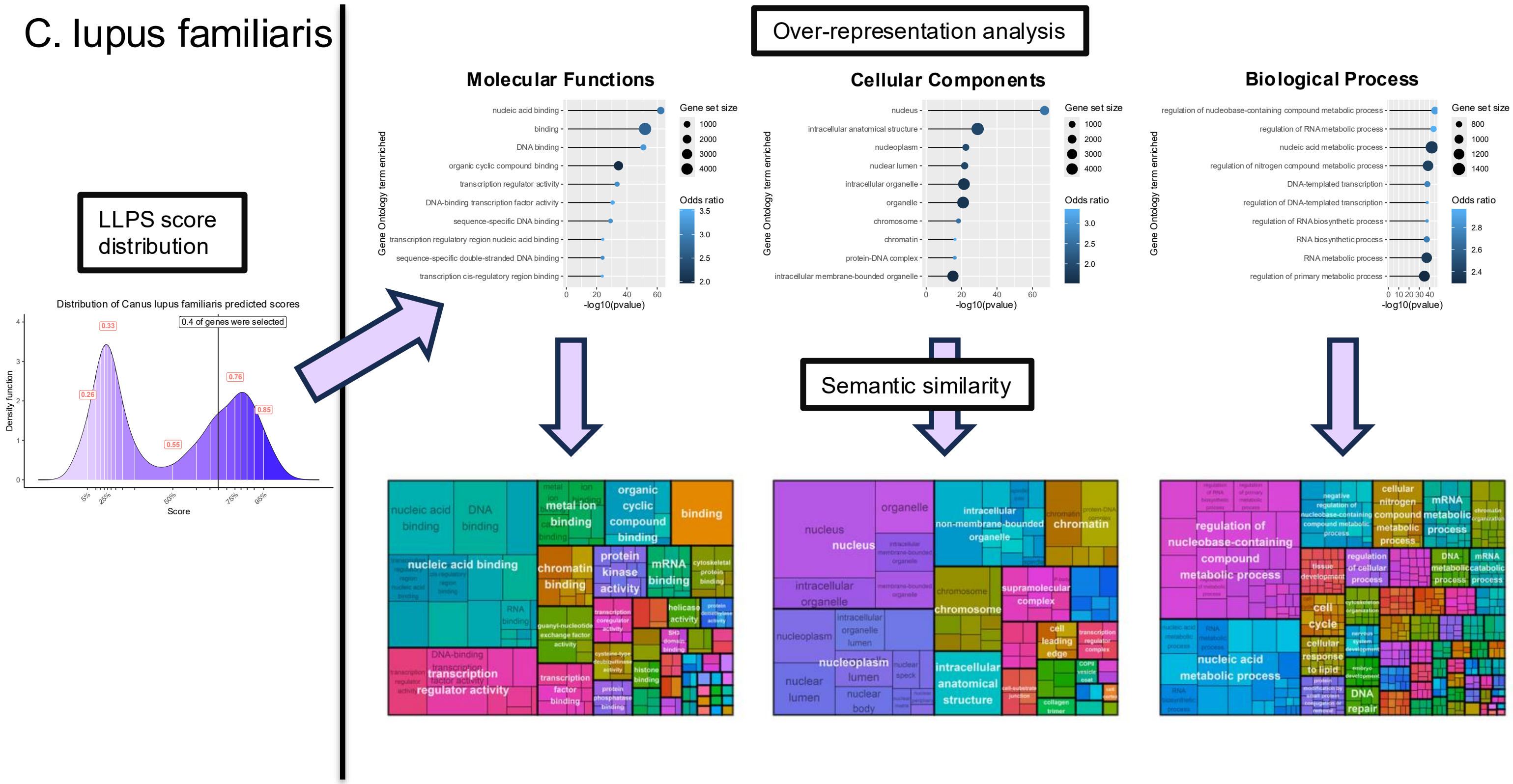


**Supplementary Figure 47: Gene Ontology (GO) enrichment results for Canus lupus familiaris.** Left plot, distribution of LLPS scores within the proteome and fraction of total Coding Sequences (CDS) selected. Upper panel, lollipop plots representing the top-10 enriched terms for the three categories: Molecular Function, Cellular Component, Biological Process. Lower panel, treemap plots illustrating hierarchical relationships between GO terms and representative subset selected.

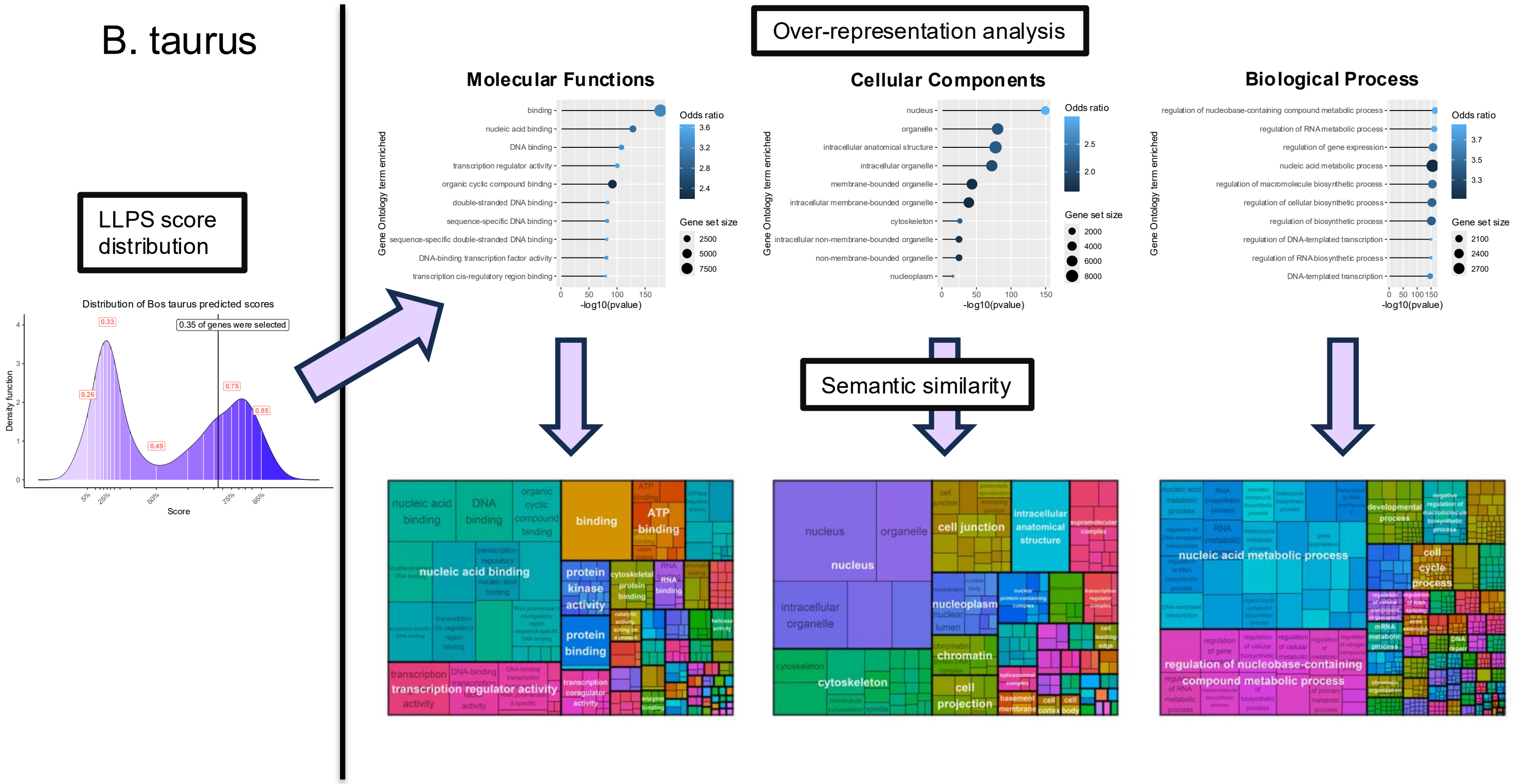


**Supplementary Figure 48: Gene Ontology (GO) enrichment results for Bos taurus.** Left plot, distribution of LLPS scores within the proteome and fraction of total Coding Sequences (CDS) selected. Upper panel, lollipop plots representing the top-10 enriched terms for the three categories: Molecular Function, Cellular Component, Biological Process. Lower panel, treemap plots illustrating hierarchical relationships between GO terms and representative subset selected.

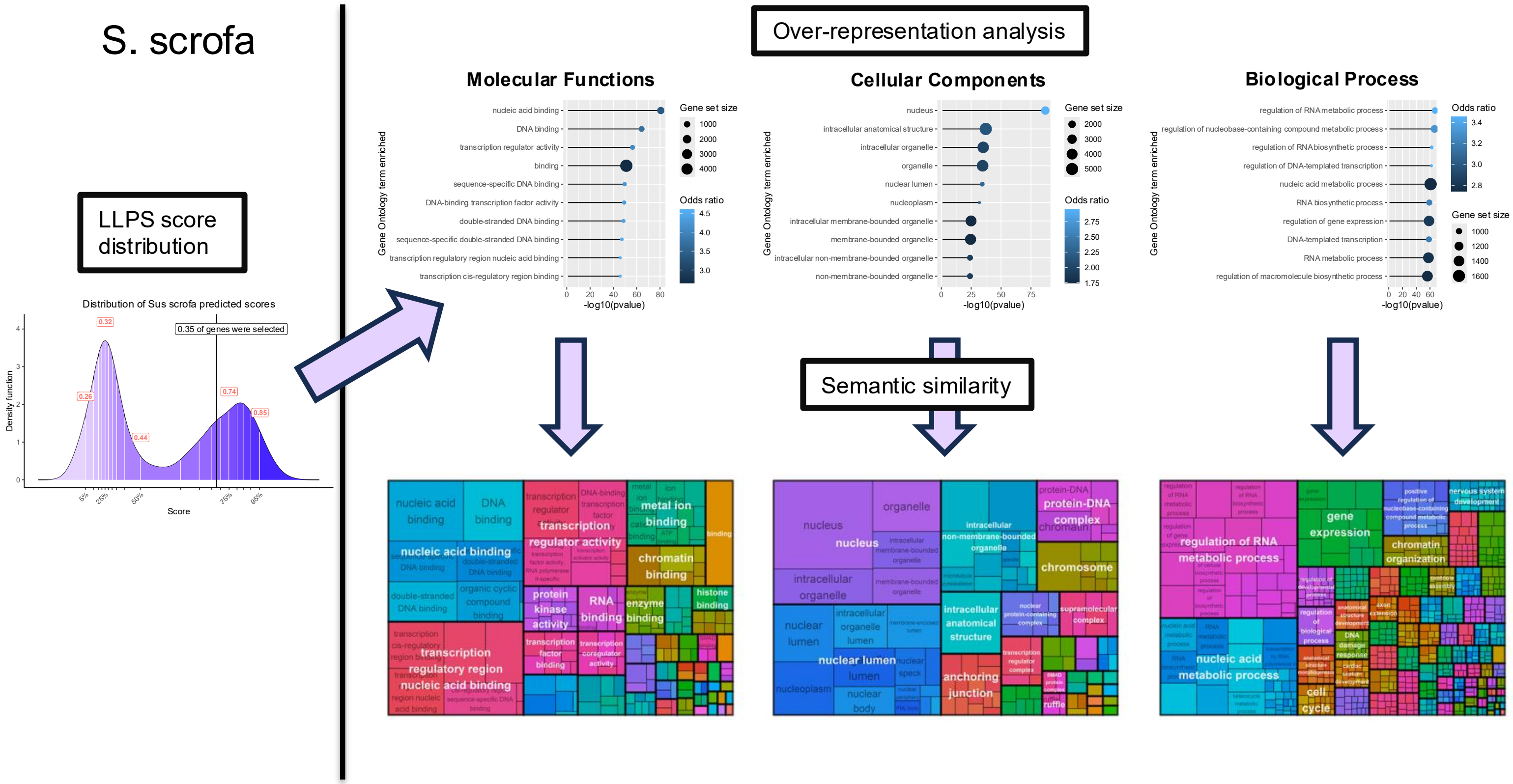


**Supplementary Figure 49: Gene Ontology (GO) enrichment results for Sus scrofa.** Left plot, distribution of LLPS scores within the proteome and fraction of total Coding Sequences (CDS) selected. Upper panel, lollipop plots representing the top-10 enriched terms for the three categories: Molecular Function, Cellular Component, Biological Process. Lower panel, treemap plots illustrating hierarchical relationships between GO terms and representative subset selected.

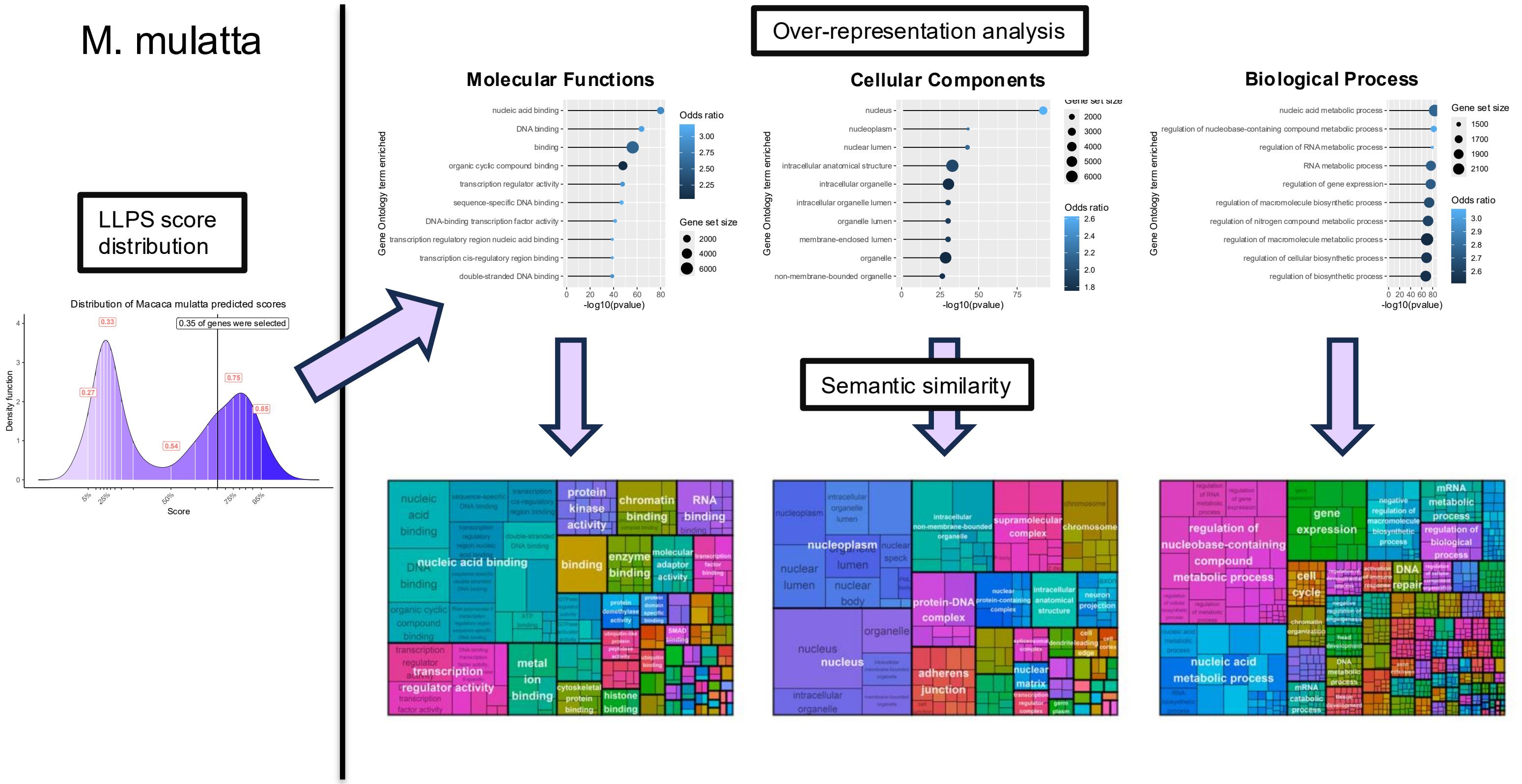


**Supplementary Figure 50: Gene Ontology (GO) enrichment results for Macaca mulatta.** Left plot, distribution of LLPS scores within the proteome and fraction of total Coding Sequences (CDS) selected. Upper panel, lollipop plots representing the top-10 enriched terms for the three categories: Molecular Function, Cellular Component, Biological Process. Lower panel, treemap plots illustrating hierarchical relationships between GO terms and representative subset selected.

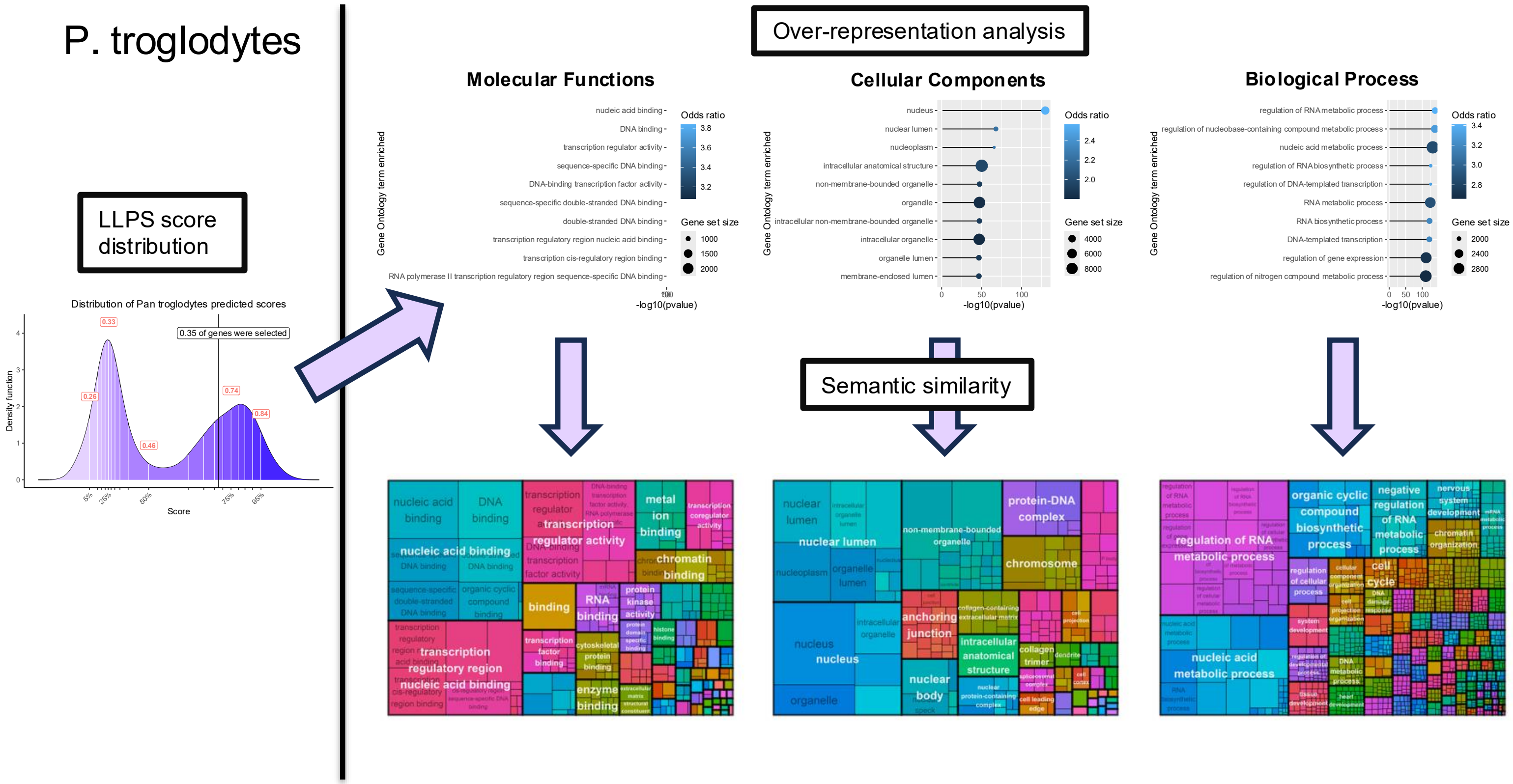


**Supplementary Figure 51: Gene Ontology (GO) enrichment results for Pan troglodytes.** Left plot, distribution of LLPS scores within the proteome and fraction of total Coding Sequences (CDS) selected. Upper panel, lollipop plots representing the top-10 enriched terms for the three categories: Molecular Function, Cellular Component, Biological Process. Lower panel, treemap plots illustrating hierarchical relationships between GO terms and representative subset selected.

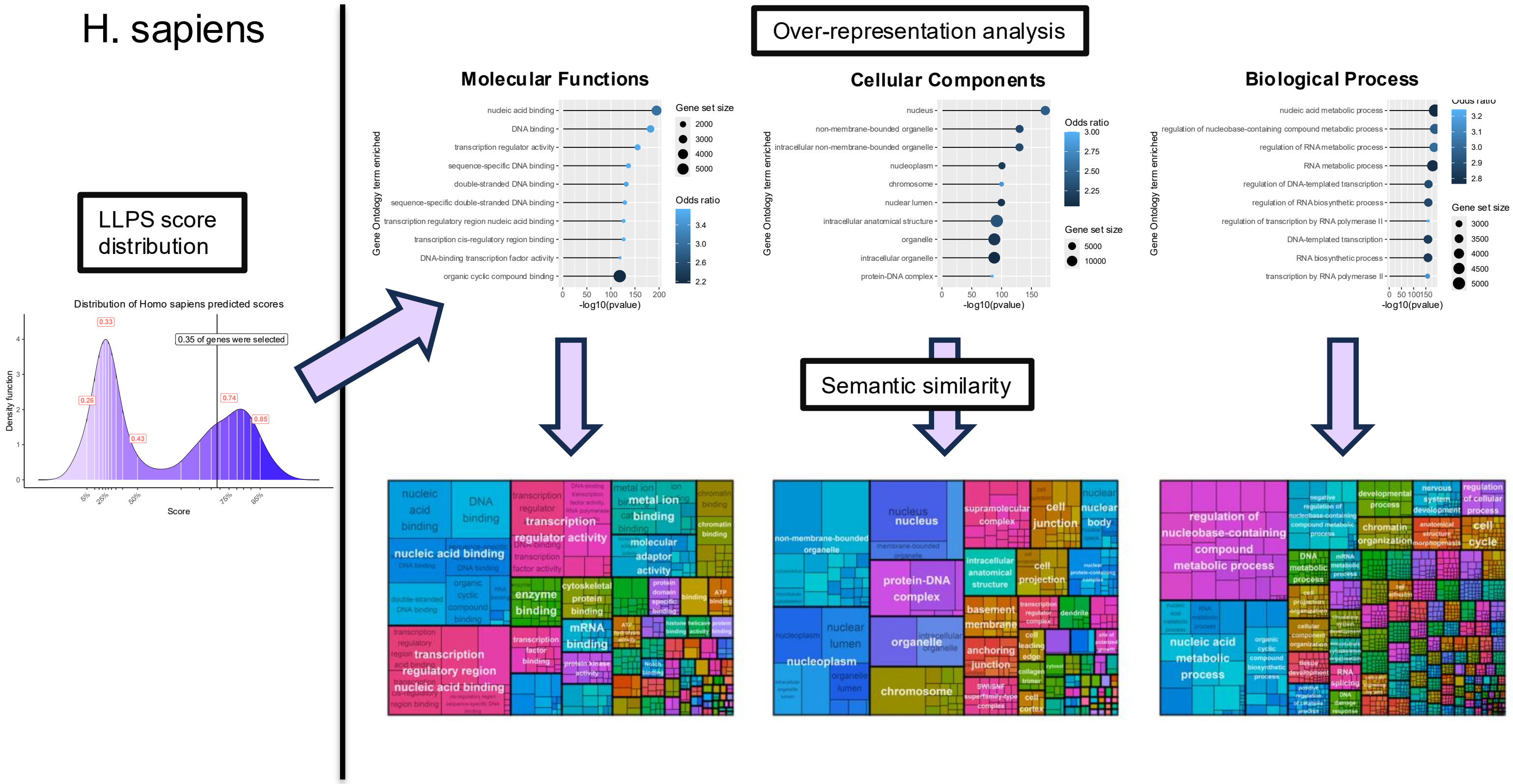


**Supplementary Figure 52: Gene Ontology (GO) enrichment results for Homo sapiens.** Left plot, distribution of LLPS scores within the proteome and fraction of total Coding Sequences (CDS) selected. Upper panel, lollipop plots representing the top-10 enriched terms for the three categories: Molecular Function, Cellular Component, Biological Process. Lower panel, treemap plots illustrating hierarchical relationships between GO terms and representative subset selected.

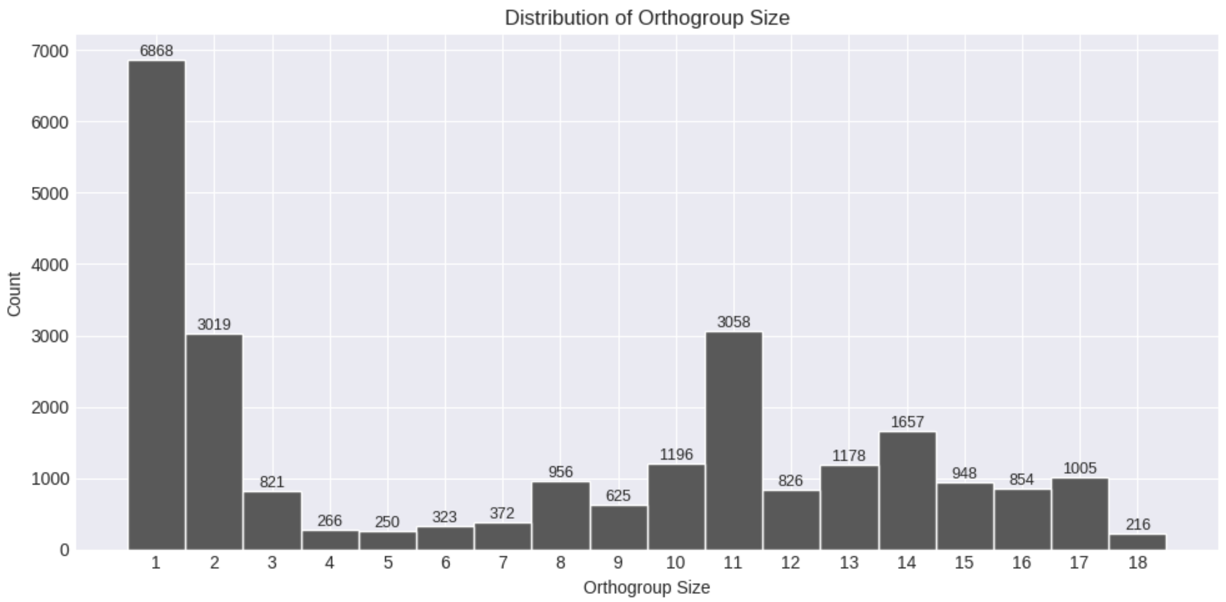


**Supplementary Figure 53: Distribution of orthogroup sizes identified by OrthoFinder.** The y axis represents the number of orthogoups which are sets of sequences with high similarity and thought to derive from the same ancestral gene. The x axis represents the orthogroup size which is the number of species having at least one sequence contributing to the orthogroup.

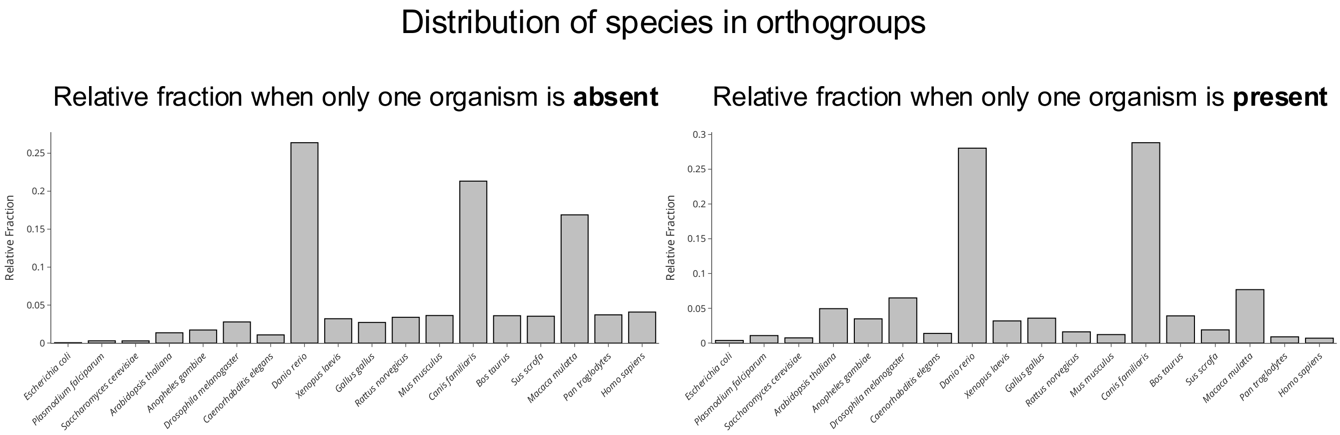


**Supplementary Figure 54: Identification of organisms with divergent LLPS phenotypes.** Normalized relative frequencies of organisms exhibiting outlier behavior in liquid-liquid phase separation (LLPS) phenotypes across orthogroups. **(a)** Organisms where the sole ortholog lacks LLPS capability. *Danio rerio* was identified as a significant outlier (one-sided Grubbs test, p = 0.0171). **(b)** Organisms where the sole ortholog displays LLPS capability. *Canis familiaris* was identified as a potential outlier (one-sided Grubbs test, p = 0.0169). Normalization accounts for genome size (ortholog count per species), enabling cross-species comparison.

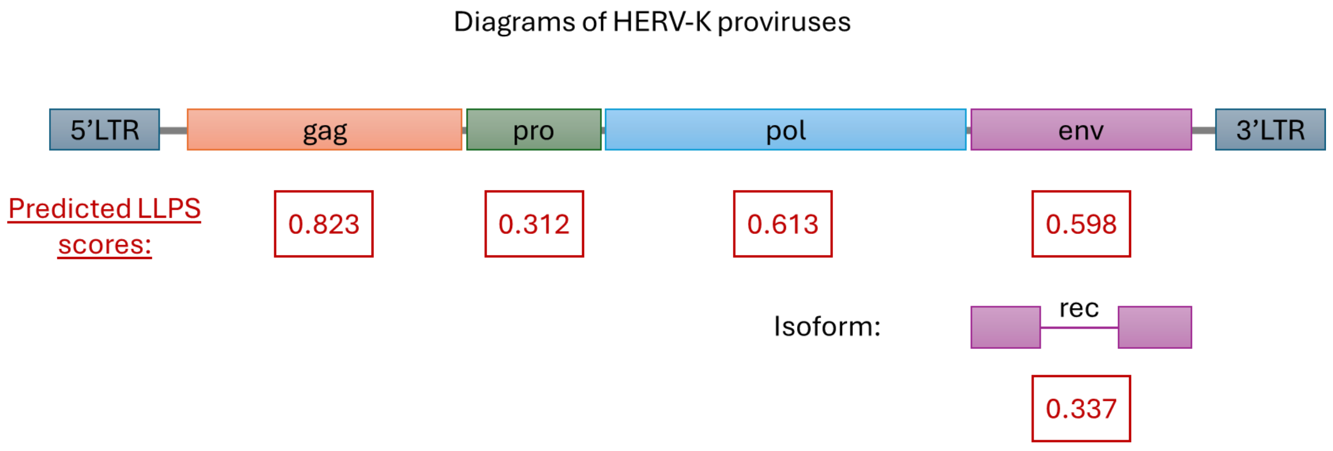


**Supplementary Figure 55: LLPS scores for five coding-sequences corresponding to the human endogenous retrovirus HERV-K113.** Human endogenous retroviruses are composed of the gag, pro, pol and env sequence, which further lead to a full length and a truncated sequence termed rec.
